# Coagulopathy signature precedes and predicts severity of end-organ heat stroke pathology in a mouse model

**DOI:** 10.1101/771410

**Authors:** Elizabeth A. Proctor, Shauna M. Dineen, Stephen C. Van Nostrand, Madison K. Kuhn, Christopher D. Barrett, Douglas K. Brubaker, Michael B. Yaffe, Douglas A. Lauffenburger, Lisa R. Leon

**Author notes:** Corresponding authors: Douglas A. Lauffenburger, 77 Massachusetts Ave 16-343, Cambridge MA 02139; Lisa R. Leon, 10 General Greene Ave, Bldg 42, Natick MA 01760.

## Abstract

Heat stroke is a life-threatening condition characterized by loss of thermoregulation and severe elevation of core body temperature, which can cause organ failure and damage to the central nervous system. While no definitive test exists to measure heat stroke severity, immune challenge is known to increase heat stroke risk, although the mechanism of this increased risk is unclear. In this study, we used a mouse model of classic heat stroke to test the effect of immune challenge on pathology. Employing multivariate supervised machine learning to identify patterns of molecular and cellular markers associated with heat stroke, we found that prior viral infection simulated with poly I:C injection resulted in heat stroke presenting with high levels of factors indicating coagulopathy. Despite a decreased number of platelets in the blood, platelets are large and non-uniform in size, suggesting younger, more active platelets. Levels of D-dimer and soluble thrombomodulin were increased in more severe heat stroke, and in cases presenting with the highest level of organ damage markers D-dimer levels dropped, indicating potential fibrinolysis-resistant thrombosis. Genes corresponding to immune response, coagulation, hypoxia, and vessel repair were up-regulated in kidneys of heat-challenged animals, and these increases correlated with both viral treatment and distal organ damage while appearing before discernible tissue damage to the kidney itself. We conclude that heat stroke-induced coagulopathy may be a driving mechanistic force in heat stroke pathology, especially when exacerbated by prior infection, and that coagulation markers may serve as an accessible biomarker for heat stroke severity and therapeutic strategies.

**Key points:**

- A signature of pro-coagulation markers predicts circadian core body temperature and levels of organ damage in heat stroke
- Changes in coagulopathy-related gene expression are evidenced before histopathological organ damage

## Introduction

Heat stroke is the most severe form of heat illness, resulting in central nervous system dysfunction, organ failure, cardiovascular damage, and death^1^. Heat stroke can be initiated passively (“classic” heat stroke) by high ambient temperatures with the inability to effectively dissipate the accumulated heat load on the body. Even upon the onset of heat illness symptoms, quantitative assessment of progression and severity is complicated, leading to uncertainty in diagnosis and evaluation of danger to the patient. Heat stroke induces similar symptoms to those induced by infection, including hyperthermia, neurological abnormalities, dehydration, and inflammation. Prior viral or bacterial infection is known to increase risk for heat stroke, potentially by pre-inducing a febrile and immune compromised state^2,3^. Elevated levels of pro-inflammatory cytokines such as IL-1β, IL-6, IFN-γ, and TNFα are common to both infection and heat stroke^4–6^, suggesting that infection can prime a harmful immune response that is synergistically promoted by heat challenge and drives severe pathology.

Currently, no definitive clinical test exists to measure severity or predict outcome and recovery time from heat stroke. The most common molecular biomarkers utilized for assessment of organ damage with heat stroke diagnosis are circulating cytokines (a measure of systemic inflammation), blood urea nitrogen and creatine (kidney injury), myoglobin (rhabdomyolysis), and the liver transaminases aspartate aminotransferase (AST) and alanine aminotransferase (ALT), which can be released by several tissues but namely skeletal muscle and liver. As such, these represent general biomarkers of organ dysfunction and damage that can be a sign of alternative, independent conditions, and are thus not exclusive markers for heat stroke^1^. Accordingly, they are not sufficiently specific for use in diagnosis, prognosis, or assignment of therapeutic strategy. To construct a quantitative measure that can be used as a diagnostic and prognostic tool, we hypothesized that shifts in the profile of blood cell populations may provide an opening for construction of a unique “fingerprint” of blood factors indicative of heat stroke and its severity. Using a mouse model of classic heat stroke developed in our laboratory, we examined the impact of prior viral or bacterial infection on hematological aspects of recovery. To avoid the influence of illness-induced fever on our model, mice were exposed to heat either 48 or 72 hours following poly I:C or LPS injection, timepoints when symptoms of illness (fever, lethargy, anorexia) were minimized or completely absent.

Here, we demonstrate the use of a complete blood count (CBC) profile as a multivariate predictor of circadian thermoregulation, with a signature indicating underlying heat stroke-induced coagulopathy (HSIC). A multivariate pattern of pro-coagulation markers further correlates with the amount of liver damage in heat-stroked animals, and genes related to immune response, coagulation, hypoxia, and wound healing are up-regulated even in histologically healthy tissue. These findings suggest a central role for HSIC that mimics disseminated intravascular coagulopathy (DIC)^7^ as a main pathophysiological driver of organ damage and mortality in classic heat stroke. Further, these results demonstrate that the coagulation response can be detected early, before tissue damage occurs, suggesting a potential therapeutic strategy to detect, treat, and halt severe heat stroke cases.

## Methods

### Animals

Male C57BL/6J mice (Jackson Laboratories) weighing 23.4±0.1 g were individually housed in Nalgene polycarbonate cages (11.5 in × 7.5 in × 5 in) fitted with HEPA-filter tops and Shepherds Specialty Blend bedding (ScottPharma) under standard laboratory conditions (25±2 °C; 12:12 hour light-dark cycle, lights on at 0600 h). Rodent laboratory chow (Harlan Teklad 7012) and water were provided *ad libitum*. Each cage was supplied with a Nalgene Mouse House, maple wood product (#W0002; Bio-Serv), and stainless steel ring attached to the wire lid for enrichment. Procedures were approved by the Institutional Animal Care and Use Committee at the United States Army Research Institute of Environmental Medicine, adhering to the *Guide for the Care and Use of Laboratory Animals* in an Association for Assessment and Accreditation of Laboratory Animal Care-accredited facility.

### Radiotelemetry transmitter implantation

Under isoflurane anesthesia, mice were intraperitoneally implanted with radiotelemetry transmitters (1.1 g, G2 Emitter; Starr Life Sciences) to measure body core temperature (T_c_±0.1 °C) and general activity (counts)^6,8^. Surgical analgesia was provided with subcutaneous buprenorphine injection (0.05 mg/kg) just prior to surgery, and again 24 and 48 hours post-procedure. Mice recovered from surgery in 7-10 days as assessed by return to pre-surgical body weight, normal food and water consumption, and stable circadian T_c_ and activity rhythms^9^. T_c_ and activity were continuously monitored at 1-minute intervals throughout recovery and experimentation using the VitalView system.

### Immune stimulant injection

Injections were performed between 0700 and 1000 h, when mice displayed a normal baseline T_c_<36.0°C. Mice were weighed and intraperitoneally (IP) injected with vehicle (equivolume of saline; SAL), polyinosinic:polycytidylic acid (100 μg/kg; poly I:C), or lipopolysaccharide (50 μg/kg; LPS) as viral and bacterial stimulants, respectively. Body weight and food and water consumption were measured daily following injection. Mice otherwise recovered undisturbed for 48 or 72 hours prior to heat stress protocol (Figure 1, Table S1, Table S2).

**Figure 1.**
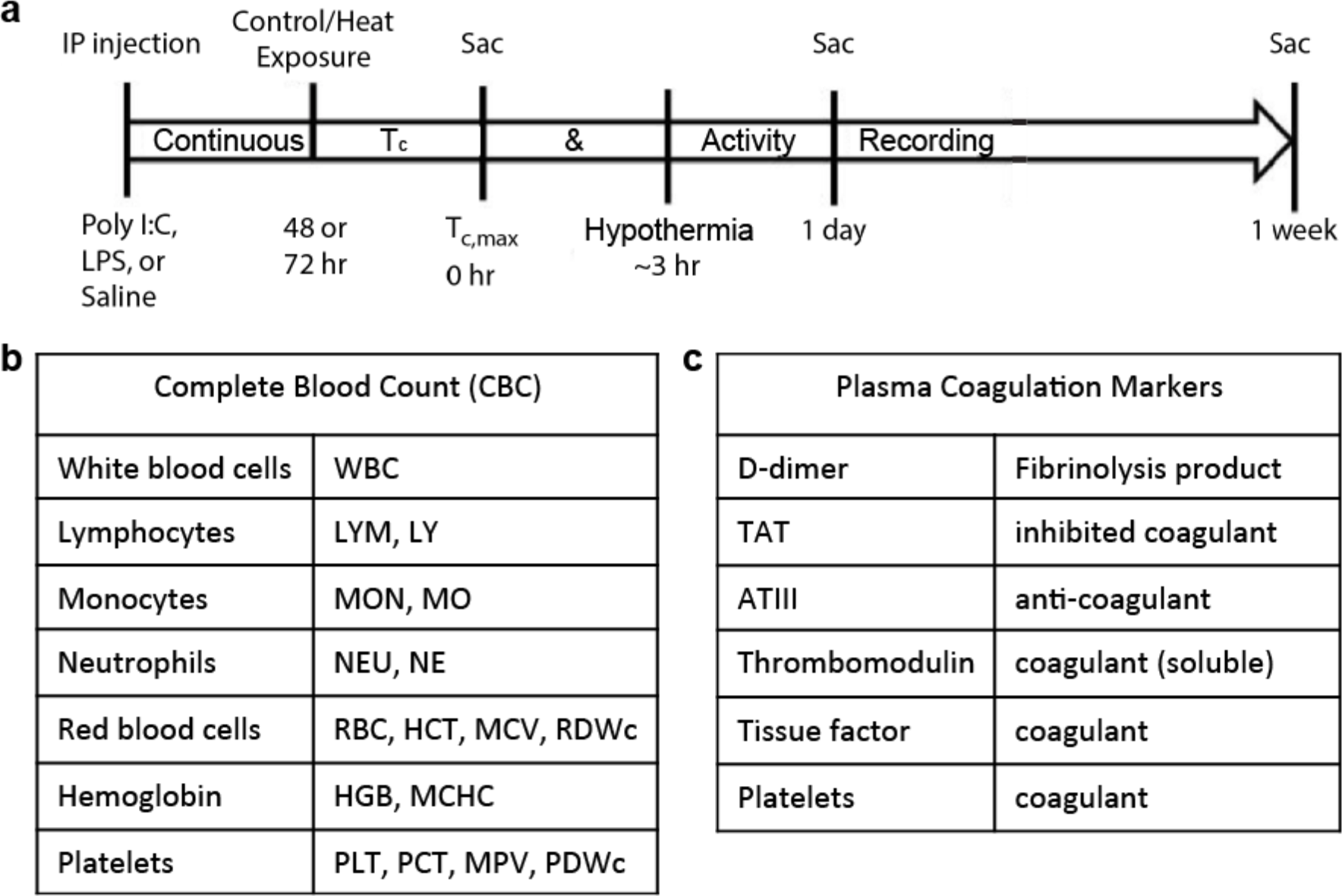
Experimental design. (a) Cohort of mice was divided into 3 groups and received an injection of poly I:C, LPS, or sterile saline. Mice were returned to their home cages for either 48 or 72 hours, after which they were exposed to heat stress at 39.5 °C, or were unheated controls at 25 °C. Mice were sacrificed at Tc,max (42.4 °C), 1 day following Tc,max, or 7 days following Tc,max, for a total of 36 groups (Table S1, Table S2) and N=13 mice per group. Upon sacrifice, organs were harvested and blood was drawn from the heart to perform (b) complete blood count (CBC) and (c) assay of plasma coagulation markers (Methods).

### Heat stress protocol

Mice in their home cages were placed into a floor-standing incubator (model 3950; Thermo Forma) at 25±2 °C the day before heat exposure to acclimate to incubator noises^8^. Cage filter tops were removed to permit air circulation. Between 0700 and 1000 h the next day, mice with baseline T_c_<36.0 °C were selected for heat stress protocol (Figure 1, Table S1, Table S2). Mice were weighed, food, water, and enrichment items were removed from the cage, and incubator ambient temperature (T_a_) was increased to 39.5±0.2 °C. Mice remained in the heated chamber until maximum T_c_ (T_c,max_) of 42.4 °C was reached, upon which mice were removed from heat, weighed, provided food and water *ad libitum*, and recovered undisturbed at 25°C until sample collection. Control mice were exposed to identical experimental conditions at T_a_ of 25±2 °C with time-matched procedures.

### Sample collection

Mice were sacrificed at timepoints for sample collection: 1) baseline (prior to heat/control exposure), 2) T_c,max_, 3) 24 hours following T_c,max_, or 4) 7 days following T_c,max_ (Figure 1, Table S1, Table S2). At sacrifice, mice were anesthetized with isoflurane and blood was collected via intracardiac puncture and immediately transferred to ethylenediaminetetraacetic acid (EDTA) microcentrifuge tubes and placed on ice. A complete blood count (CBC) was performed on blood samples using a VetScan HM5 Hematology Analyzer (Abaxis), including (Figure 1b, Figure S1): white blood cell count (WBC), lymphocyte count (LYM), % lymphocytes (LY), monocyte count (MON), % monocytes (MO), neutrophil count (NEU), % neutrophils (NE), red blood cell count (RBC), hematocrit (HCT), mean corpuscular volume (MCV), width of red blood cell size distribution (RDWc), hemoglobin (HGB), mean corpuscular hemoglobin concentration (MCHC), platelet count (PLT), % platelets (PCT), mean platelet volume (MPV), width of platelet size distribution (PDWc). Concentrations of plasma coagulative factors (Figure 1c, Figure S1) were measured by ELISA according to manufacturer protocol: D-dimer (Mybiosource), thrombin-antithrombin complex, antithrombin III (Abcam), soluble thrombomodulin (R&D Systems), and tissue factor (BosterBio), as well as granzyme B, a marker of liver damage (ThermoFisher).

### Partial least squares modeling

We used partial least squares regression^10^ and discriminant analysis with variable importance of projection (VIP) scores to identify a signature of blood factors correlating with heat stroke pathology, as measured by core body temperature (T_c_) and granzyme B concentration. Modeling was conducted in MATLAB using PLS_Toolbox (Eigenvector Research). Data was normalized along each parameter by Z-score. Cross-validation was performed with one-third of the relevant dataset, or with one-fifth of the relevant dataset if N<15. The number of latent variables (LVs) was chosen to minimize cumulative error over all predictions. Where noted, we orthogonally rotated models to achieve maximal separation across LV1. We calculated model confidence by randomly permuting Y to form a distribution of error for 100 random models, and comparing our model to this distribution with the Mann-Whitney U test. Importance of each parameter to the model prediction was quantified using VIP score, an average of the weights of each parameter over all latent variables normalized by latent variable percent of variance. A VIP score greater than 1 (above average contribution) was considered important for model performance and prediction.

### RNAseq

RNA was purified from 20 mg tissue samples of kidney using the RNeasy Mini Prep kit with DNase (Qiagen) according to manufacturer protocol. Mice were selected from the larger cohort as 6 mice randomly chosen from each of the 8 sub-groups undergoing 48-hour post-injection incubation: saline- or poly I:C-treated, heated or unheated, T_c,max_ or 1 day timepoint. RNAseq was performed by the BioMicroCenter at the Massachusetts Institute of Technology on a HiSeq2000 with read length of 40SE, using Kapa mRNA hyperprep for library preparation.

### Transcriptomic analysis

Gene expression was mapped using the STAR/RSEM pipeline with *Mus musculus* GRCm38.93 as reference (Ensembl). We estimated variance-mean dependence in count data and tested for differential expression based on a model using the negative binomial distribution with the DESeq package in R from the Bioconductor repository. Six outliers were identified by principal component analysis (PCA), and later confirmed by quality control to be contaminated or feature low read counts. These samples were excluded from further analysis. Remaining samples were ranked using the log2-fold change and PCA loadings and submitted for pre-ranked gene set enrichment analysis^11^ (GSEA) of heated poly I:C-treated mice compared with non-heated saline-treated mice at the 1 day timepoint. Enrichment for Hallmark Gene Sets was calculated using the classic scoring scheme. Full GSEA combining T_c,max_ and 1 day timepoints was performed as confirmation, using the Diff_Of_Classes metric with phenotype permutation.

Log2-fold change of gene transcripts was used as input to the *prcomp* function in R, and loadings were extracted using *get_pca_var*. The most predictive principal components for the four given classes (heat challenge, poly I:C injection) were determined by multiclass log regression and leave-one-out cross validation. Principal components 2 and 3 were chosen as the best model, with class prediction accuracy of 90.5%.

### Histopathology

Samples of liver, kidney, spleen, lung, and duodenum from the same 48 mice examined with RNAseq were formalin-fixed and submitted to IDEXX BioResearch for histopathology analysis. The submitted tissues were trimmed, processed, blocked, sectioned, stained with H&E, and examined microscopically. Observed microscopic changes were graded utilizing a standard system: 0=no significant change, 1=minimal, 2=mild, 3=moderate, and 4=severe.

## Results

### Hematology signature predicts difference in circadian T_c_

Although core body temperature (T_c_) is used clinically as a measure of heat stroke severity, individual responses to a given value of T_c_ are highly variable^1^ and not necessarily indicative of internal pathology. Immune involvement in heat stroke pathology provides an opening for potential assessment of severity of patient response, where the ideal biomarkers should be easily and quickly accessible such that they are practical for clinical use. We thus aimed to identify immune biomarkers circulating in blood quantifiable with a standard clinical test and correlating with a known measure of thermoregulatory control. As T_c_ is the most commonly used metric for heat stroke diagnosis, even though imperfect, we used the circadian differential in average T_c_, ΔT_c_ (awake T_c_ minus sleep T_c_), as a measure of thermal response and heat stroke severity. We found that quantification of the distributions of blood cell populations in a complete blood count (CBC) can predict ΔT_c_ (Figure 2). While analysis of individual parameters was inconclusive (Figure S1), our orthogonalized partial least squares regression model (Methods) predicts changes in circadian thermoregulation from covariation among CBC panel measurements, separating individual mice along a spectrum from those demonstrating “flipped” circadian thermoregulation (Figure 2a, blue), with higher T_c_ during sleep, to those with “normal” circadian thermoregulation (Figure 2a, red). The full pattern of factors distinguishing thermoregulatory regime (Figure 2b) indicates depressed immune cell populations with increasing heat stroke severity. We also found increased numbers of red blood cells, with a smaller average size but widened size distribution. This pattern is indicative of dehydration, which is commonly experienced both in heat stroke and infection, and synergistically in the presence of both stimuli. Low platelet levels were also correlated with heat stroke severity, and their large size and broad size distribution indicate younger, more active platelets^12^, introducing the potential for coagulopathy. Variable importance in projection (VIP) analysis (Methods) identified the red blood cell signature and platelet signature as the dominating factors in determining ΔT_c_, and thus the most relevant to heat stroke pathology (Figure 2b, red bars). Importantly, the overall pattern predictive of the circadian thermoregulation proxy for heat stroke severity is a faint but highly statistically significant signal among the biological “noise” of individual variation in response to heat and infection stimulus, making up only 9.8% of the variation in blood cell populations but explaining 31% of the variation in ΔT_c_ (p=0.008).

**Figure 2.**
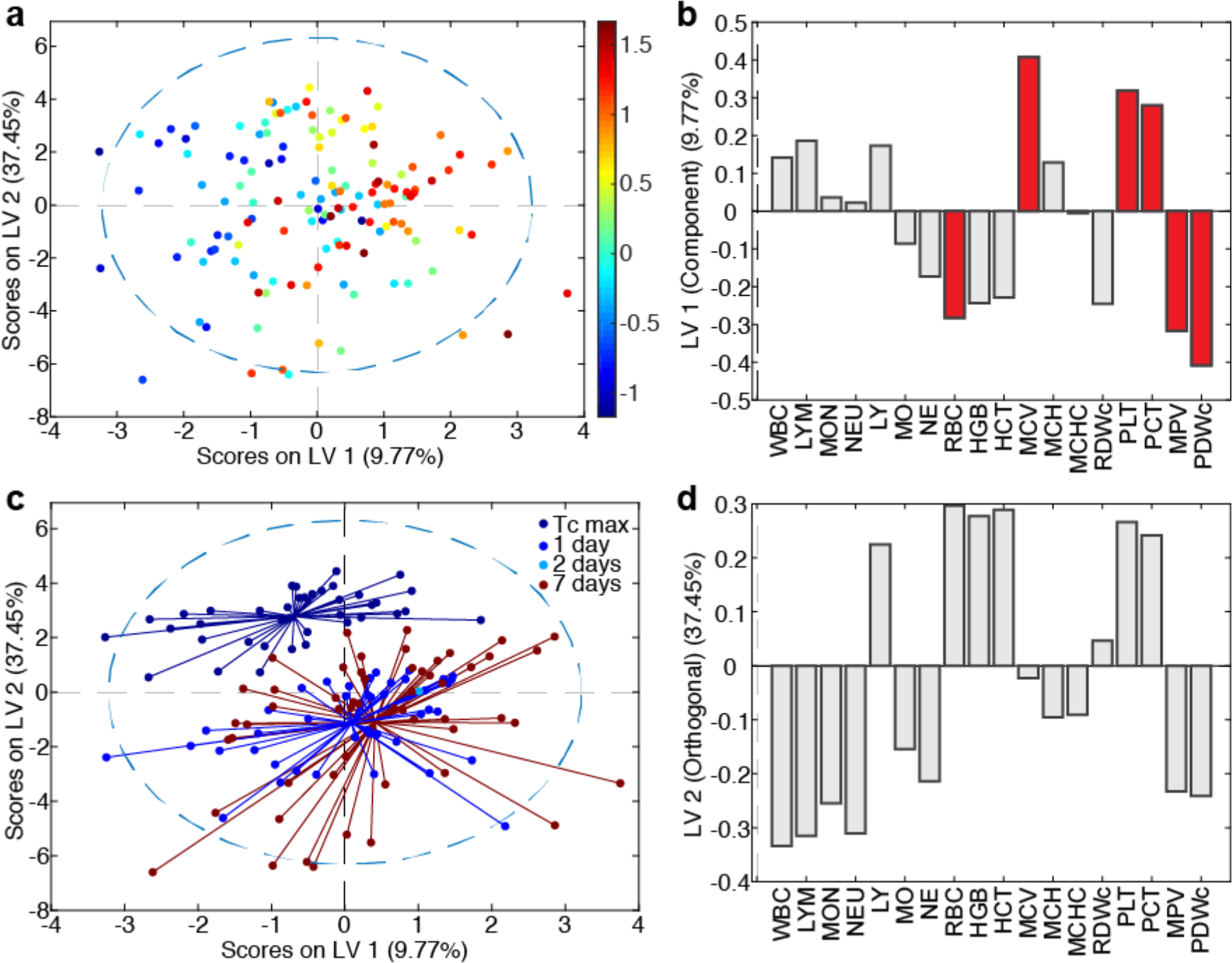
Multivariate hematological signature predicts circadian ΔT_c_, a proxy of heat stroke severity. (a) Orthogonalized partial least squares regression model uses covariation in complete blood count results to separate mice on a spectrum of ΔT_c_ as average T_c,day_ - average T_c,night_ (color bar, °C). N=137, four latent variables, cross-validated using one-third of input data. R^2^ of cross-validation: 0.205, Wilcoxon cross-validation p-value 0.008. (b) Loadings on latent variable 1 (LV1) represent contributions of each factor to ΔT_c_. Red bars indicate VIP≥1, a greater than-average contribution to correlation. (c) Recoloring of existing model according to timepoint of sacrifice. LV2 represents differences between CBC at T_c,max_, 1 day, 2 days, and 7 days. (d) Loadings on LV2 represent contribution of each factor to timeline of pathology.

We assessed the timeline of changes in blood cell populations in heat stroke pathology by mapping the timepoint of each blood sample onto the existing partial least squares regression model (Figure 2c). We found that, without providing any information about timeline in the original model, the time elapsed following removal from heat stress formed a major axis of variation and separation in the data, representing 37.5% of the overall variation in blood cell populations. While dehydration and lowered immune cell counts are ameliorated at later timepoints, the pathological signature of thrombocytopenia strengthens (Figure 2d), suggesting a delayed physiological response that could include platelet-consumptive coagulopathy.

### Plasma coagulative markers predict liver damage

To further investigate potential coagulopathy indicated by the thrombocytopenia signature predictive of ΔT_c_, we measured levels of coagulation and fibrinolysis markers (from here forward collectively called coagulative markers) in the blood by ELISA: D-dimer, thrombin-antithrombin complex (TAT), antithrombin (ATIII), and soluble thrombomodulin (TM). Levels of tissue factor were also assessed, but could not be performed in a sufficient number of mice per group for statistically significant analysis. We constructed an orthogonalized partial least squares regression model correlating these markers with hepatic concentration of granzyme B, a serine protease secreted by cytotoxic lymphocytes that mediates apoptosis and indicates organ damage. In order to decrease the number of variables in our model, we included only mice treated with saline or poly I:C, undergoing a 48-hour post-injection incubation, sacrificed at T_c,max_ or 1 day following. We excluded mice treated with LPS from this analysis, as LPS is known to promote coagulopathy and could therefore confound measurement of the contribution of the coagulation system to heat stroke pathology. We excluded the 7-day timepoint from our model because mouse-to-mouse variability in recovery from heat stroke stimulus was amplified at this later timepoint, resulting in non-relevant noise that interfered with the predictive molecular signature.

The resulting orthogonalized partial least squares regression model predicts levels of hepatic granzyme B using concentrations of 4 circulating coagulative markers (Figure 3), separating mice along a spectrum of no (blue) to high (red) liver damage with 54% of the variation in the coagulative markers explaining 50% of the liver damage phenotype (p=0.004) (Figure 3a). Examination of treatment conditions, which were not considered in the model, reveals a synergistic relationship between previous viral infection and heat challenge in coagulopathy-derived organ damage (Figure 3), with saline injection (Figure 3b, blue) at the low end of the spectrum and poly I:C plus heat (Figure 3b, red) on the extreme high end of the spectrum. The pattern of factors defining this spectrum of increasing organ damage highlights high levels of soluble thrombomodulin and D-dimer as the primary predictors of pathology (Figure 3c), demonstrating upregulated activation of coagulation and fibrinolysis pathways leading to (i) clot formation, as D-dimer is only released after formation of cross-linked fibrin clots; (ii) fibrinolysis, i.e. the breakdown of clots as seen by the release of D-dimer; and (iii) resistance to fibrinolysis, as soluble thrombomodulin greatly augments activation of thrombin-activatable fibrinolysis inhibitor (TAFI)^13^.

**Figure 3.**
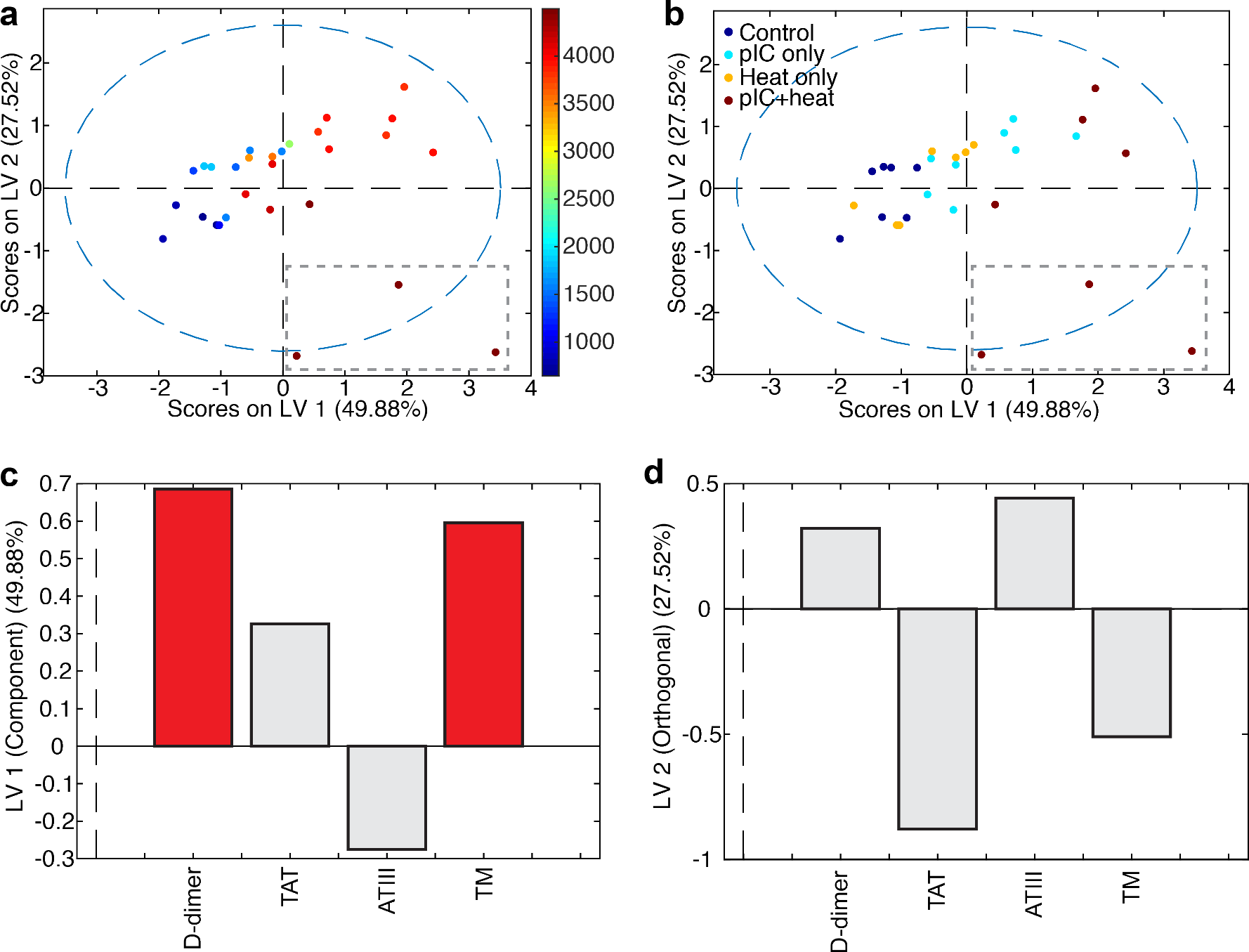
Coagulation perturbations correlate with liver damage in heat- and viral-challenged mice. (a) Orthogonalized partial least squares regression model uses covariation in levels of circulating coagulation markers to separate mice on a spectrum of liver damage from none (blue) to high (red), as measured by levels of granzyme B (color bar, pg/mL). Grey dashed box highlights three outliers in (d). N=30, two latent variables, cross-validated using one-third of input data. R2 of cross-validation: 0.518, Wilcoxon cross-validation p-value 0.004. (b) Recoloring of existing model according to treatment, 48 hours prior to heat challenge. Differences in treatment lie on the same axis (LV1) as the granzyme B spectrum, indicating correlation between treatment and organ damage. Grey dashed box highlights three outliers in (d). (c) Loadings on latent variable 1 (LV1) represent contributions of each factor to concentration of granzyme B. Red bars indicate VIP≥1, a greater than-average contribution to correlation. (d) Three outliers differ sufficiently to create an orthogonal axis of variation, LV2, upon which they separate from other points. Loadings on LV2 represent distinguishing factors of these outliers with extreme pathology, including extremely high values of thrombomodulin but decreased D-dimer, potentially due to fibrinolysis resistance.

Additionally, three animals exhibited a sufficiently different relationship between molecular coagulative markers and liver damage from the rest of the subjects that they defined their own latent variable (Figure 3a,b, grey dashed box), representing 27.5% of the overall variation in the dataset. These three animals presented with pronounced liver damage and extremely high levels of soluble thrombomodulin, but much lower levels of D-dimer than their granzyme B concentration would suggest (Figure 3d). We interpreted this significant deviation from the model fit as evidence of fibrinolysis resistance, which may be reasonably expected with very high levels of soluble thrombomodulin due to TAFI activation^13^, and is often indicative of platelet-rich clots.

### Immune response and clotting factor genes up-regulated in histologically normal tissues

As the above results have demonstrated, the immune and hematologic consequences of heat stroke are evident at the tissue level soon after heat challenge. To further elucidate the molecular mechanisms involved in instigating these tissue-level changes, we performed transcriptomic analysis on livers and kidneys of mice from the most severely affected groups, mice treated with poly I:C and heat-challenged 48 hours following treatment, along with comparable controls (saline-treated, unheated mice). We targeted the T_c,max_ and 1 day timepoints to avoid the biological noise created by the high variability in recovery observed at 7 days. While RNA in livers had already begun to degrade and was unsuitable for analysis, pathology was less advanced in kidneys. Among the most differentially-expressed genes between severe heat stroke and controls were those involved in metabolism, inflammation, apoptosis, and coagulation (Figure 4a). However, genes involved in these processes were both up- and down-regulated, leading to confusion in interpretation. Because the accumulation of many small changes can result in a larger effect than a small number of large changes, we performed pre-ranked Gene Set Enrichment Analysis^11^ (GSEA) to identify overall classes of genes that are differentially-expressed in the poly I:C, heat-challenged mice. While down-regulated gene sets almost exclusively revealed decreases in metabolic processes (Table S3), we identified multiple up-regulated gene sets involved in inflammatory response, hypoxia, wound healing, and coagulation (FDR q-value < 0.25) (Figure 4b, Table S3).

**Figure 4.**
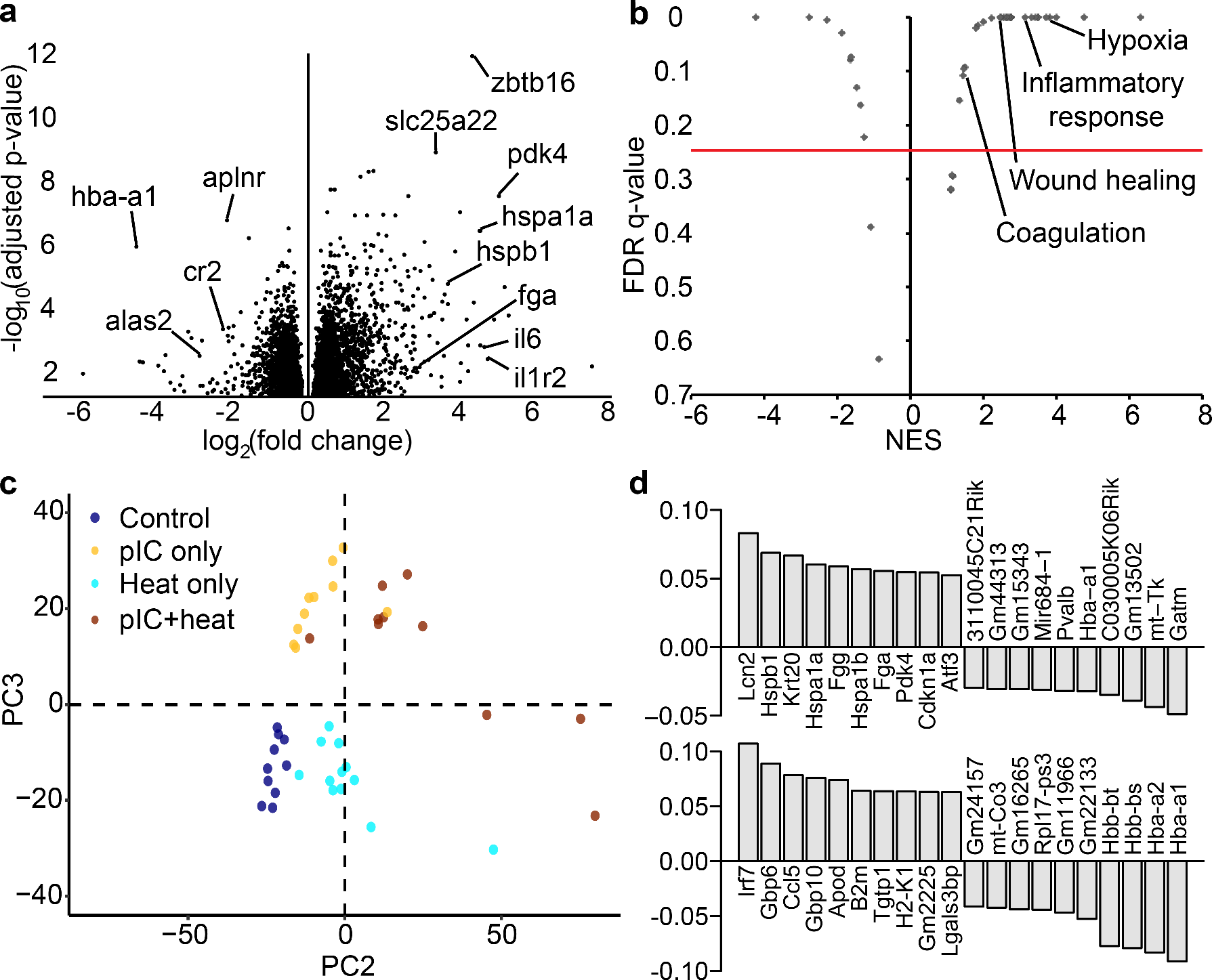
Transcription changes in genes related to vascular injury response in heat stroke. (a) Volcano plot showing differentially expressed genes (log2(fold change)) in kidneys of poly I:C-treated, heat-challenged mice, and their statistical significance as adjusted p-value. (b) Volcano plot showing highly enriched gene sets (NES, normalized enrichment score) and their statistical significance as FDR q-value. (c) Principal component analysis of gene transcripts separates groups by heat application (PC2, 9% of variation in gene transcripts) or poly I:C injection (PC3, 5% of variation in gene transcripts). (d) Top ten positive and top ten negative contributing gene transcripts to heat response (PC2, top) and poly I:C response (PC3, bottom).

To identify patterns of co-varying gene transcripts that together differentiate heat-stroked animals, we performed principal component analysis using all non-zero transcripts (Figure 4c,d). The two principal components separating the treatment groups revealed key players in heat stroke response related to the gene sets identified by GSEA: decreased *hba* and *hbb*, components of hemoglobin that promote fibrinolysis when released from red blood cells^14^; increased *lcn2*, involved in innate immunity and an early biomarker for acute kidney injury; increased *krt20*, a filament protein contributing to structural integrity of epithelial cells; increased *hspa1a* and *hspb1*, protein folding chaperones in response to environmental stress; and increased *fgg* and *fga*, components of fibrinogen, the most abundant protein in blood clots. Notably, we also observe a >2-fold increase in expression of *sdc1*, a marker of vascular activation and injury (adjusted p=0.021). Taken together, these results strongly suggest pre-symptomatic co-occurring vascular damage, reduced oxygen transportation machinery in erythrocytes leading to ischemia, coagulopathy, end-organ damage, and inflammatory response.

## Discussion

Disseminated intravascular coagulopathy (DIC) and similar coagulopathic phenotypes are promoted in infection^15^, and also in heat stroke^16^. The two coagulopathy-promoting stimuli experienced together, or in short succession, may produce a synergistic effect, as we observed in this work. Coagulopathy is thus a prime candidate for the mechanism of the increased risk of heat stroke-related pathology following bacterial or viral infection, and consequently the targeting of early coagulation perturbations before onset of clinical coagulopathy may be beneficial to preventing incipient or future heat stroke^17,18^. The thrombocytopenia we observed in heat-stroked mice is a likely a consequence of DIC-like phenotype^17,19–21^, which we have termed heat-stroke induced coagulopathy (HSIC), although a contribution from immune-mediated destruction of platelets is also possible^22^. Additional support for HSIC as a main mechanism of heat stroke pathological progression to organ dysfunction and failure is the appearance of either or both thrombosis, bleeding, or both in heat stroke patients and animal models^23–25^, a similar phenotype to that observed in DIC^7^.

Taken together, our findings suggest that both pathways may occur simultaneously, with hyperthermia producing a coagulopathy that drives the observed pathology in tissues, instigating an immune reaction to platelet-rich clots and a cytokine storm in response to vascular injury, which together create a feed-forward synergistic reaction to generate incipient multi-organ damage. Our gene expression findings support this proposed feed-forward mechanism: histopathology reports from kidneys assayed with RNAseq indicated no tissue-level pathology (Table S4), while on the molecular level we found highly significant increased expression of genes involved in immune response, wound healing, and hypoxia. We thus are able to distinctly detect molecular-level changes that distinguish the strongest responders to heat challenge before pathology propagates to have tissue-level effects detrimental to organ function. Conversely, while liver samples in these same mice displayed distinct tissue-level pathology including coagulative necrosis (Table S4), RNA in these tissues had already begun to degrade and we were unable to isolate transcripts at high enough quality for sequencing. Overall, the changes in these specific sets of genes suggest that tissue is being deprived of blood due to thrombosis and/or breakdown of vascular structures (e.g., capillaries), and a wound healing response is initiated. These findings are further supported by a report from Hagiwara *et al.*^26^, that administration of recombinant thrombomodulin, a co-factor that enhances the Protein C-mediated anticoagulant response of vascular endothelium, results in reduced organ damage in a rat model of heat stress-induced inflammation.

Because no large clots or hemorrhages were observed in tissue histology, we conclude that the coagulation and corresponding injury to vessel walls occurs at the capillary level, similar to what is observed in DIC. In the case of heat stroke, hyperthermia may be directly damaging the vessel walls of small capillaries, causing the release of tissue factor and initiating the coagulation process. Although we could not test all of the animals studied here for tissue factor levels due to the small volume of blood available in each mouse and the large panel of assays we performed, we did observe heightened levels of tissue factor in heat-stroked animals when all timepoints and incubation times were pooled (Figure S1).

We have presented in this work, to our knowledge, the first RNAseq analysis of tissue from an animal model of heat stroke. While other studies have presented gene expression assayed by quantitative PCR^27–29^, the advantage of RNAseq is the agnostic measure of the full transcriptome, rather than a set of pre-chosen RNAs. This agnostic search allows us to assay a greater extent of perturbation to the system state of the tissue, without narrowing our view with expectations. We can also extrapolate the effect of the measured perturbations using prior knowledge from the STRING protein-protein interaction database^30^. We see that the identified RNAs are interconnected at the protein level, and that transcriptional dysregulation could lead to synergistic dysfunction, disrupting tissue function to a larger degree than evident from individual dysregulated species or processes (Figure 5). We find clusters of genes that regulate immune function, connected via the complement system and coagulation to groups of genes involved in metabolism and stress response. Two tightly-linked clusters of genes involved in cell cycle regulation dominate the center of the network, acting as a hub connecting many other gene sets, with genes involved in tissue growth and remodeling spread throughout.

**Figure 5.**
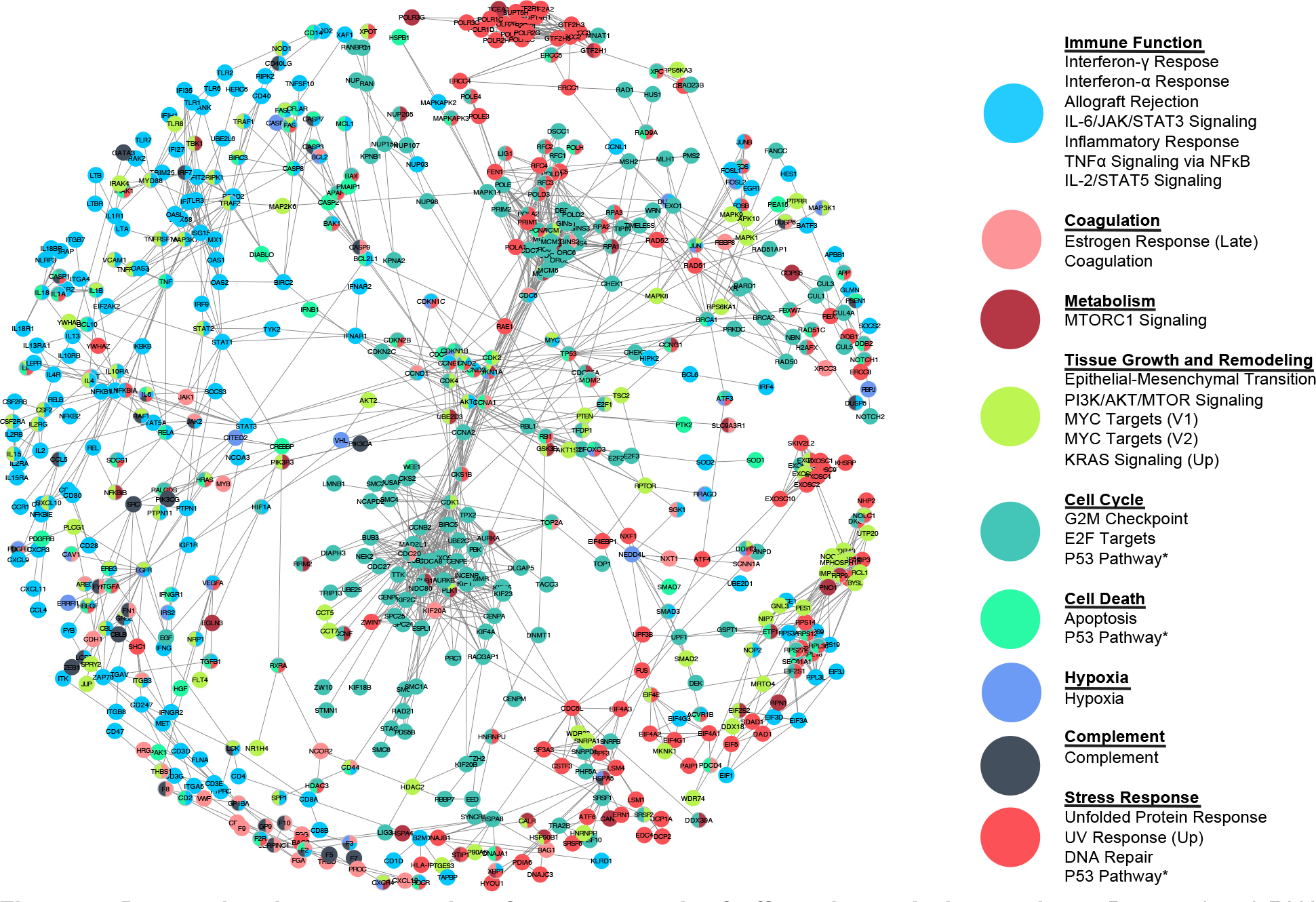
Dysregulated gene transcripts form a network of affected protein interactions. Dysregulated RNA transcripts in heat-stroked mice as identified by combined_score from the STRING database of protein-protein interactions, including experiments, co-expression, and gene fusion evidence of interaction. Nodes (gene transcripts) are colored by HALLMARK gene sets, grouped as listed. Nodes were chosen as leading edge subsets of gene sets with enrichment FDR q-value < 0.25 in poly I:C, heated mice over saline, unheated mice. Edges were chosen as STRING combined_score > 0.99^31^, with shorter length representing higher score (edge-weighted, spring-embedded layout). Nodes classified into more than one enriched gene set are colored as a pie chart^32^.

In summary, we have demonstrated in an animal model of heat stroke that (1) heat stroke severity can be predicted immediately after heat exposure through a simple and widely available clinical test (complete blood count with differential); (2) a coagulopathy (HSIC) with diagnostic features similar to DIC occurs early after heat exposure and before tissue-level histopathologic changes in end-organs can be observed; (3) a complex series of concurrent changes at the gene expression and protein levels highlight increases in immune/inflammatory responses, hypoxia, wound healing, and coagulation; and (4) prior viral infection may increase heat stroke risk and severity through a feed-forward coagulative mechanism. These findings highlight the importance of further investigation into the presentation and mechanisms of HSIC in heat stroke, which could lead to targeted treatment of the early coagulopathic component to halt progression of the condition to organ failure. Such treatments may take the form of focused blood component transfusions or drugs (*e.g.*, tranexamic acid or recombinant factor VIIa) for the bleeding phenotype, or anticoagulation therapy (*e.g.*, heparinization) for the clotting phenotype.

## Acknowledgements

This work was supported by the Institute for Collaborative Biotechnologies through contract W911NF-09-D-0001 from the US Army Research Office, and by a grant from the USAMRAA program W81XWH-13-MOMJPC5-IPPEHA. EAP was partially supported by the MIT CEHS Training Grant in Environmental Toxicology T32-ES007020. CDB and MBY were supported by NIH-DOD Grant UM1-HL120877. The authors thank Brian A. Joughin and Justin R. Pritchard for their valuable input.

The opinions or assertions contained herein are the private views of the author(s) and are not to be construed as official or as reflecting the views of the Army or the Department of Defense.

In conducting the research described in this report, the investigators adhered to the “Guide for the Care and Use of Laboratory Animals” as prepared by the Committee for the Update of the Guide for the Care and Use of Laboratory Animals of the Institute for Laboratory Animal Research, National Research Council.

Citations of commercial organizations and trade names in this report do not constitute an official Department of the Army endorsement or approval of the products or services of these organizations.

## Authorship Contributions

EAP, SMD, DAL, and LRL conceived and planned experiments and analysis; EAP and SMD carried out experiments; EAP, SCVN, MKK, and DKB analyzed data; EAP, CDB, DKB, MBY, DAL, and LRL interpreted results; EAP, CDB, DAL, and LRL wrote the manuscript with input from all authors. All authors provided critical feedback and helped shape the research, analysis, and manuscript.

