## Supplemental Material for "Coagulopathy signature precedes and predicts severity of end-organ heat stroke pathology in a mouse model"

**Table S1. Experimental groups.** Treatment (injected immune stimulus), post-treatment incubation time before heat challenge, heat challenge type, and timepoint of sacrifice for animals in each experimental group.

| Group | Treatment | | | Heat | | Timepoint | | | Incubation | |
|---|---|---|---|---|---|---|---|---|---|---|
| | Saline | LPS | pIC | No | Yes | $T_{c,max}$ | 1 d | 7 d | 48 hr | 72 hr |
| 1 | ■ | | | ■ | | ■ | | | ■ | |
| 2 | ■ | | | ■ | | ■ | | | | ■ |
| 3 | ■ | | | ■ | | | ■ | | ■ | |
| 4 | ■ | | | ■ | | | ■ | | | ■ |
| 5 | ■ | | | ■ | | | | ■ | ■ | |
| 6 | ■ | | | ■ | | | | ■ | | ■ |
| 7 | ■ | | | | ■ | ■ | | | ■ | |
| 8 | ■ | | | | ■ | ■ | | | | ■ |
| 9 | ■ | | | | ■ | | ■ | | ■ | |
| 10 | ■ | | | | ■ | | ■ | | | ■ |
| 11 | ■ | | | | ■ | | | ■ | ■ | |
| 12 | ■ | | | | ■ | | | ■ | | ■ |
| 13 | | ■ | | ■ | | ■ | | | ■ | |
| 14 | | ■ | | ■ | | ■ | | | | ■ |
| 15 | | ■ | | ■ | | | ■ | | ■ | |
| 16 | | ■ | | ■ | | | ■ | | | ■ |
| 17 | | ■ | | ■ | | | | ■ | ■ | |
| 18 | | ■ | | ■ | | | | ■ | | ■ |
| 19 | | ■ | | | ■ | ■ | | | ■ | |
| 20 | | ■ | | | ■ | ■ | | | | ■ |
| 21 | | ■ | | | ■ | | ■ | | ■ | |
| 22 | | ■ | | | ■ | | ■ | | | ■ |
| 23 | | ■ | | | ■ | | | ■ | ■ | |
| 24 | | ■ | | | ■ | | | ■ | | ■ |
| 25 | | | ■ | ■ | | ■ | | | ■ | |
| 26 | | | ■ | ■ | | ■ | | | | ■ |
| 27 | | | ■ | ■ | | | ■ | | ■ | |
| 28 | | | ■ | ■ | | | ■ | | | ■ |
| 29 | | | ■ | ■ | | | | ■ | ■ | |
| 30 | | | ■ | ■ | | | | ■ | | ■ |
| 31 | | | ■ | | ■ | ■ | | | ■ | |
| 32 | | | ■ | | ■ | ■ | | | | ■ |
| 33 | | | ■ | | ■ | | ■ | | ■ | |
| 34 | | | ■ | | ■ | | ■ | | | ■ |
| 35 | | | ■ | | ■ | | | ■ | ■ | |
| 36 | | | ■ | | ■ | | | ■ | | ■ |

**Table S2. Experimental biological replicates.** Number of replicates (N) for each measurement in each experimental group, as defined in Table S1 and illustrated in Figure S1.

| Group | N | | | | | | | |
|---|---|---|---|---|---|---|---|---|
| | $T_{c,ave}$ (day) | $T_{c,ave}$ (night) | Corr BW | Dehydration | WBC | LYM | MON | NEU |
| **1** | 10 | 10 | 10 | 10 | 9 | 9 | 9 | 9 |
| **2** | 10 | 10 | 10 | 10 | 11 | 11 | 11 | 11 |
| **3** | 9 | 9 | 10 | 10 | 9 | 9 | 9 | 9 |
| **4** | 9 | 9 | 10 | 10 | 9 | 9 | 9 | 9 |
| **5** | 10 | 10 | 10 | 10 | 10 | 10 | 10 | 10 |
| **6** | 10 | 10 | 10 | 10 | 9 | 9 | 8 | 8 |
| **7** | 10 | 10 | 10 | 10 | 8 | 5 | 5 | 5 |
| **8** | 10 | 10 | 10 | 10 | 9 | 2 | 2 | 2 |
| **9** | 10 | 10 | 10 | 10 | 9 | 9 | 9 | 9 |
| **10** | 10 | 10 | 10 | 10 | 9 | 9 | 9 | 9 |
| **11** | 10 | 10 | 10 | 10 | 10 | 10 | 10 | 10 |
| **12** | 10 | 10 | 10 | 10 | 10 | 10 | 10 | 10 |
| **13** | 10 | 10 | 10 | 10 | 10 | 10 | 10 | 10 |
| **14** | 9 | 9 | 9 | 9 | 9 | 9 | 9 | 9 |
| **15** | 10 | 10 | 10 | 10 | 9 | 9 | 9 | 9 |
| **16** | 10 | 10 | 10 | 10 | 9 | 9 | 9 | 9 |
| **17** | 10 | 10 | 10 | 10 | 10 | 10 | 10 | 10 |
| **18** | 10 | 10 | 10 | 10 | 10 | 10 | 10 | 10 |
| **19** | 10 | 10 | 10 | 10 | 10 | 9 | 9 | 9 |
| **20** | 10 | 10 | 10 | 10 | 10 | 10 | 10 | 10 |
| **21** | 10 | 10 | 10 | 10 | 8 | 8 | 8 | 8 |
| **22** | 10 | 10 | 10 | 10 | 9 | 9 | 9 | 9 |
| **23** | 10 | 10 | 10 | 10 | 10 | 10 | 9 | 9 |
| **24** | 10 | 10 | 10 | 10 | 10 | 10 | 10 | 10 |
| **25** | 10 | 10 | 10 | 10 | 10 | 10 | 10 | 10 |
| **26** | 10 | 10 | 10 | 10 | 10 | 10 | 10 | 10 |
| **27** | 10 | 10 | 10 | 10 | 7 | 7 | 7 | 7 |
| **28** | 10 | 10 | 8 | 7 | 10 | 10 | 10 | 10 |
| **29** | 10 | 10 | 10 | 10 | 10 | 10 | 10 | 10 |
| **30** | 10 | 10 | 10 | 10 | 10 | 10 | 10 | 10 |
| **31** | 10 | 10 | 10 | 10 | 9 | 6 | 6 | 6 |
| **32** | 10 | 10 | 10 | 10 | 8 | 7 | 7 | 7 |
| **33** | 10 | 10 | 10 | 10 | 4 | 2 | 2 | 2 |
| **34** | 10 | 10 | 8 | 8 | 8 | 8 | 8 | 8 |
| **35** | 10 | 10 | 10 | 10 | 6 | 6 | 6 | 6 |
| **36** | 10 | 10 | 10 | 10 | 9 | 9 | 9 | 9 |

| Group | N | | | | | | | | | | | |
|---|---|---|---|---|---|---|---|---|---|---|---|---|
| | LY | MO | NE | RBC | HGB | HCT | MCV | MCH | MCHC | RDWc | PLT | PCT |
| **1** | 9 | 9 | 9 | 9 | 9 | 9 | 9 | 9 | 9 | 9 | 9 | 9 |
| **2** | 11 | 11 | 11 | 11 | 11 | 11 | 11 | 11 | 11 | 11 | 11 | 11 |
| **3** | 9 | 9 | 9 | 9 | 9 | 9 | 9 | 9 | 9 | 9 | 9 | 9 |
| **4** | 9 | 9 | 9 | 9 | 9 | 9 | 9 | 9 | 9 | 9 | 9 | 9 |
| **5** | 10 | 10 | 10 | 10 | 10 | 10 | 10 | 10 | 10 | 10 | 10 | 10 |
| **6** | 8 | 9 | 9 | 9 | 9 | 9 | 9 | 9 | 9 | 9 | 9 | 9 |
| **7** | 5 | 5 | 5 | 8 | 8 | 8 | 8 | 8 | 8 | 8 | 8 | 8 |
| **8** | 2 | 2 | 2 | 9 | 9 | 9 | 9 | 9 | 9 | 9 | 9 | 9 |

| | | | | | | | | | | | | |
|---|---|---|---|---|---|---|---|---|---|---|---|---|
| **9** | 9 | 9 | 9 | 9 | 9 | 9 | 9 | 9 | 9 | 9 | 9 | 9 |
| **10** | 9 | 9 | 9 | 9 | 9 | 9 | 9 | 9 | 9 | 9 | 9 | 9 |
| **11** | 10 | 10 | 10 | 10 | 10 | 10 | 10 | 10 | 10 | 10 | 10 | 10 |
| **12** | 10 | 10 | 10 | 10 | 10 | 10 | 10 | 10 | 10 | 10 | 10 | 10 |
| **13** | 10 | 10 | 10 | 10 | 10 | 10 | 10 | 10 | 10 | 10 | 10 | 10 |
| **14** | 9 | 9 | 9 | 9 | 9 | 9 | 9 | 9 | 9 | 9 | 9 | 9 |
| **15** | 9 | 9 | 9 | 9 | 9 | 9 | 9 | 9 | 9 | 9 | 9 | 9 |
| **16** | 9 | 9 | 9 | 9 | 9 | 9 | 9 | 9 | 9 | 9 | 9 | 9 |
| **17** | 10 | 10 | 10 | 10 | 10 | 10 | 10 | 10 | 10 | 10 | 10 | 10 |
| **18** | 10 | 10 | 10 | 10 | 10 | 10 | 10 | 10 | 10 | 10 | 10 | 10 |
| **19** | 9 | 9 | 9 | 10 | 10 | 10 | 10 | 10 | 10 | 10 | 10 | 10 |
| **20** | 10 | 10 | 10 | 10 | 10 | 10 | 10 | 10 | 10 | 10 | 10 | 10 |
| **21** | 8 | 8 | 8 | 8 | 8 | 8 | 8 | 8 | 8 | 8 | 8 | 8 |
| **22** | 9 | 9 | 9 | 9 | 9 | 9 | 9 | 9 | 9 | 9 | 9 | 9 |
| **23** | 10 | 9 | 9 | 10 | 10 | 10 | 10 | 10 | 10 | 10 | 10 | 10 |
| **24** | 10 | 10 | 10 | 10 | 10 | 10 | 10 | 10 | 10 | 10 | 10 | 10 |
| **25** | 10 | 10 | 10 | 10 | 10 | 10 | 10 | 10 | 10 | 10 | 10 | 10 |
| **26** | 10 | 10 | 10 | 10 | 10 | 10 | 10 | 10 | 10 | 10 | 10 | 10 |
| **27** | 7 | 7 | 7 | 7 | 7 | 7 | 7 | 7 | 7 | 7 | 7 | 7 |
| **28** | 10 | 10 | 10 | 10 | 10 | 10 | 10 | 10 | 10 | 10 | 10 | 10 |
| **29** | 10 | 10 | 10 | 10 | 10 | 10 | 10 | 10 | 10 | 10 | 10 | 10 |
| **30** | 10 | 10 | 10 | 10 | 10 | 10 | 10 | 10 | 10 | 10 | 10 | 10 |
| **31** | 6 | 6 | 6 | 9 | 9 | 9 | 9 | 9 | 9 | 9 | 9 | 9 |
| **32** | 7 | 7 | 7 | 8 | 8 | 8 | 8 | 8 | 8 | 8 | 8 | 8 |
| **33** | 2 | 2 | 2 | 4 | 4 | 4 | 4 | 4 | 4 | 4 | 4 | 4 |
| **34** | 8 | 8 | 8 | 8 | 8 | 8 | 8 | 8 | 8 | 8 | 8 | 8 |
| **35** | 6 | 6 | 6 | 6 | 6 | 6 | 6 | 6 | 6 | 6 | 6 | 6 |
| **36** | 9 | 9 | 9 | 9 | 9 | 9 | 9 | 9 | 9 | 9 | 9 | 9 |

| **Group** | **N** | | | | | | | |
|---|---|---|---|---|---|---|---|---|
| | **MPV** | **PDWc** | **Granzyme B** | **D-dimer** | **TAT** | **ATIII** | **Thrombomodulin** | **Tissue Factor** |
| **1** | 9 | 9 | 7 | 7 | 5 | 5 | 5 | 8 |
| **2** | 11 | 11 | 0 | 7 | 5 | 5 | 5 | 4 |
| **3** | 9 | 9 | 6 | 5 | 5 | 5 | 5 | 0 |
| **4** | 9 | 9 | 7 | 5 | 5 | 5 | 5 | 0 |
| **5** | 10 | 10 | 6 | 5 | 5 | 5 | 5 | 0 |
| **6** | 9 | 9 | 8 | 5 | 5 | 5 | 5 | 0 |
| **7** | 8 | 8 | 7 | 7 | 5 | 5 | 5 | 7 |
| **8** | 9 | 9 | 0 | 9 | 5 | 5 | 5 | 4 |
| **9** | 9 | 9 | 7 | 5 | 5 | 5 | 5 | 0 |
| **10** | 9 | 9 | 0 | 5 | 5 | 5 | 5 | 0 |
| **11** | 10 | 10 | 7 | 5 | 5 | 5 | 5 | 0 |
| **12** | 10 | 10 | 0 | 5 | 5 | 5 | 5 | 0 |
| **13** | 10 | 10 | 6 | 0 | 0 | 0 | 0 | 0 |
| **14** | 9 | 9 | 0 | 0 | 0 | 0 | 0 | 0 |
| **15** | 9 | 9 | 6 | 0 | 0 | 0 | 0 | 0 |
| **16** | 9 | 9 | 7 | 0 | 0 | 0 | 0 | 0 |
| **17** | 10 | 10 | 6 | 0 | 0 | 0 | 0 | 0 |
| **18** | 10 | 10 | 6 | 0 | 0 | 0 | 0 | 0 |
| **19** | 10 | 10 | 7 | 0 | 0 | 0 | 0 | 0 |
| **20** | 10 | 10 | 0 | 0 | 0 | 0 | 0 | 0 |
| **21** | 8 | 8 | 7 | 0 | 0 | 0 | 0 | 0 |

| | | | | | | | | |
|---|---|---|---|---|---|---|---|---|
| **22** | 9 | 9 | 0 | 0 | 0 | 0 | 0 | 0 |
| **23** | 10 | 10 | 7 | 0 | 0 | 0 | 0 | 0 |
| **24** | 10 | 10 | 0 | 0 | 0 | 0 | 0 | 0 |
| **25** | 10 | 10 | 6 | 9 | 5 | 5 | 5 | 5 |
| **26** | 10 | 10 | 0 | 8 | 5 | 5 | 5 | 8 |
| **27** | 7 | 7 | 7 | 5 | 5 | 5 | 5 | 0 |
| **28** | 10 | 10 | 5 | 5 | 5 | 5 | 5 | 0 |
| **29** | 10 | 10 | 6 | 5 | 5 | 5 | 5 | 0 |
| **30** | 10 | 10 | 4 | 5 | 5 | 5 | 5 | 0 |
| **31** | 9 | 9 | 6 | 8 | 5 | 5 | 5 | 5 |
| **32** | 8 | 8 | 0 | 10 | 5 | 5 | 5 | 8 |
| **33** | 4 | 4 | 5 | 5 | 5 | 5 | 5 | 0 |
| **34** | 8 | 8 | 0 | 5 | 5 | 5 | 5 | 0 |
| **35** | 6 | 6 | 7 | 5 | 5 | 5 | 5 | 0 |
| **36** | 9 | 9 | 0 | 5 | 5 | 5 | 5 | 0 |

**Figure S1. Levels of individual parameters by experimental group and by treatment group over time.** Values displayed for each experimental group, either numbered for ease of reading according to Table S1 or color-coded as in figure legend. Assays are terminal, not longitudinal, with each time point representing different animals. Biological replicates (N) for each measurement in each group are listed in Table S2. Bar or data point indicates mean, error bars represent standard error.

**A. Average day-time core body temperature** is an average over 1-minute intervals over the 12-hour light (sleep) period. Time points are measured as time after animals are removed from heat. Groups are only statistically distinguishable by treatment, with the exception of pIC/no heat/72 hour recovery group having less severe changes in core body temperature. Healthy animals commonly have lower core body temperature during sleep.

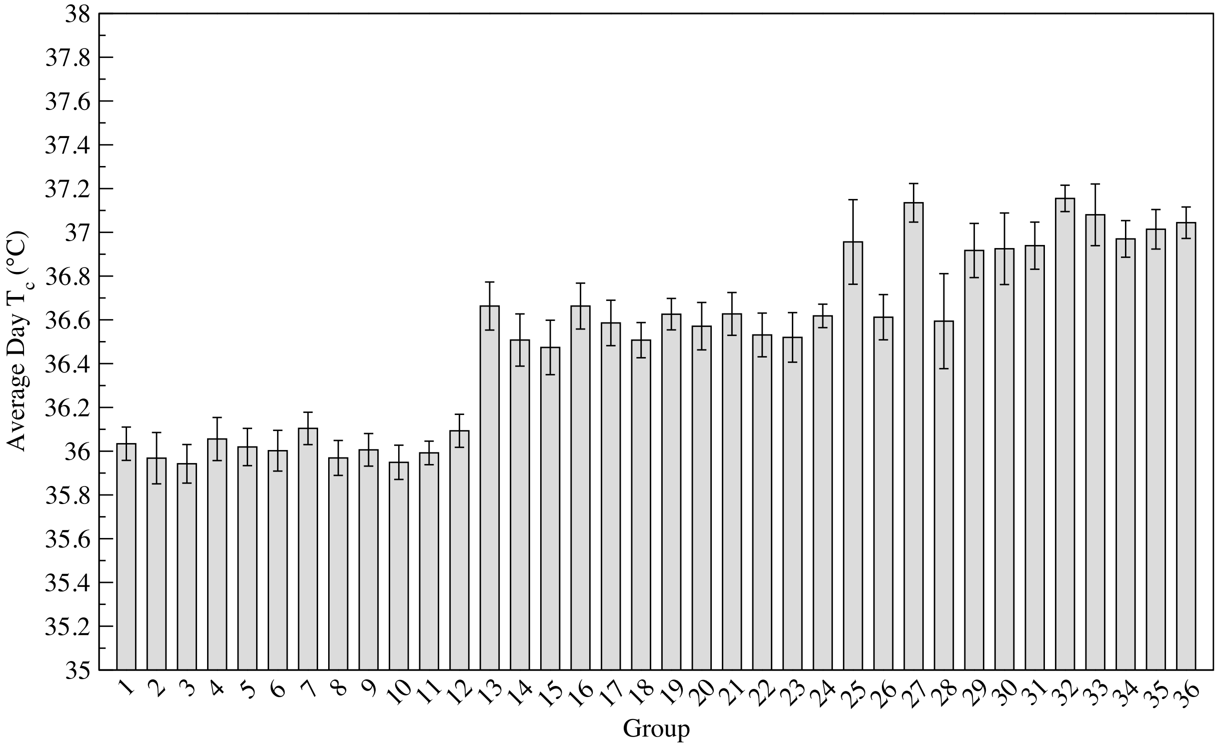


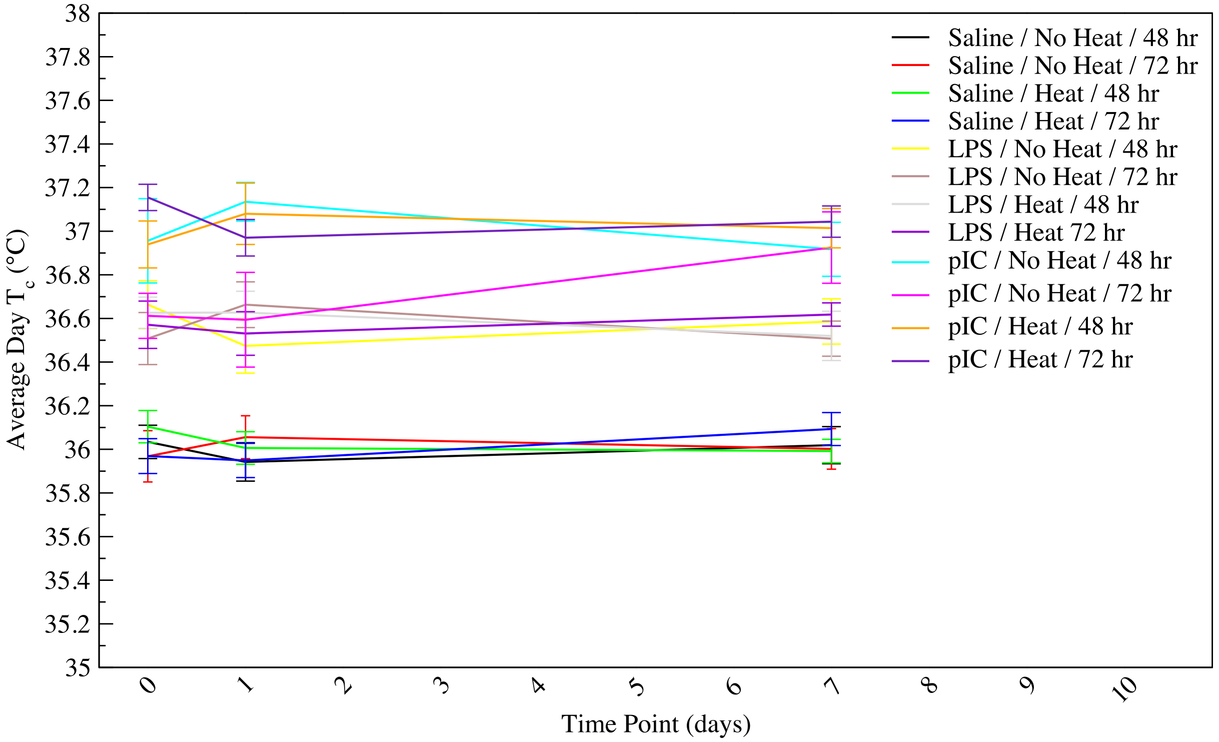

**B. Average night-time core body temperature** is an average over 1-minute intervals over the 12-hour dark (awake) period. Time points are measured as time after animals are removed from heat. Groups are only statistically distinguishable by treatment, with some overlap between LPS and pIC treated animals. Healthy animals commonly have higher core body temperature while awake.

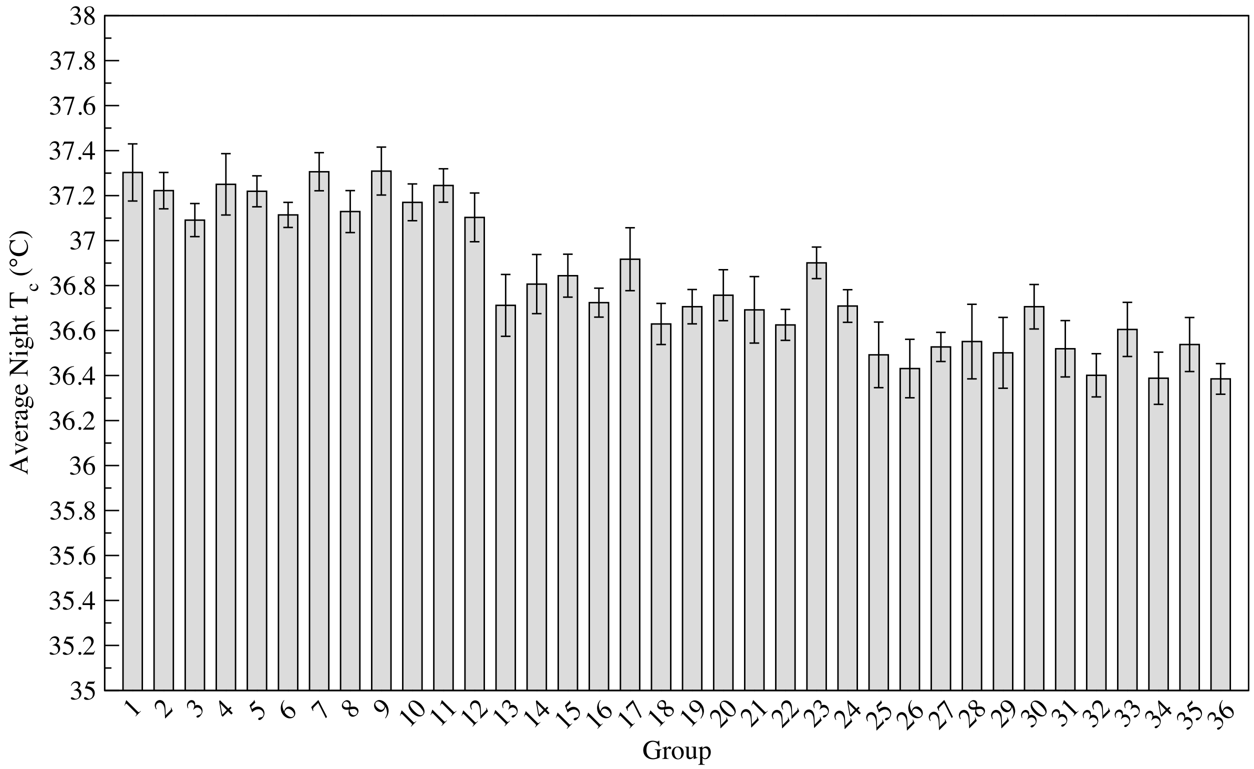


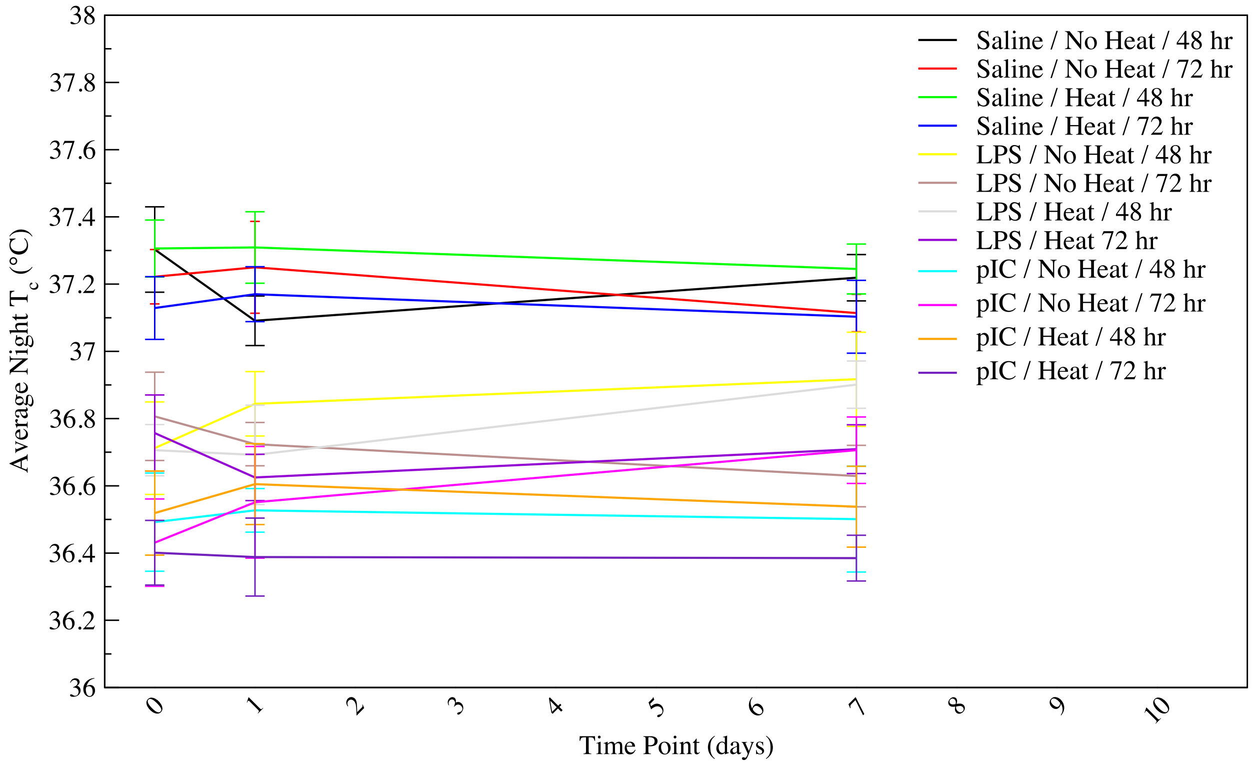

**C. Corrected body weight** accounts for water weight lost to dehydration. Variability is high, with a slight tendency for pIC treated mice to have lower initial body weight but no discernible statistically significant trends.

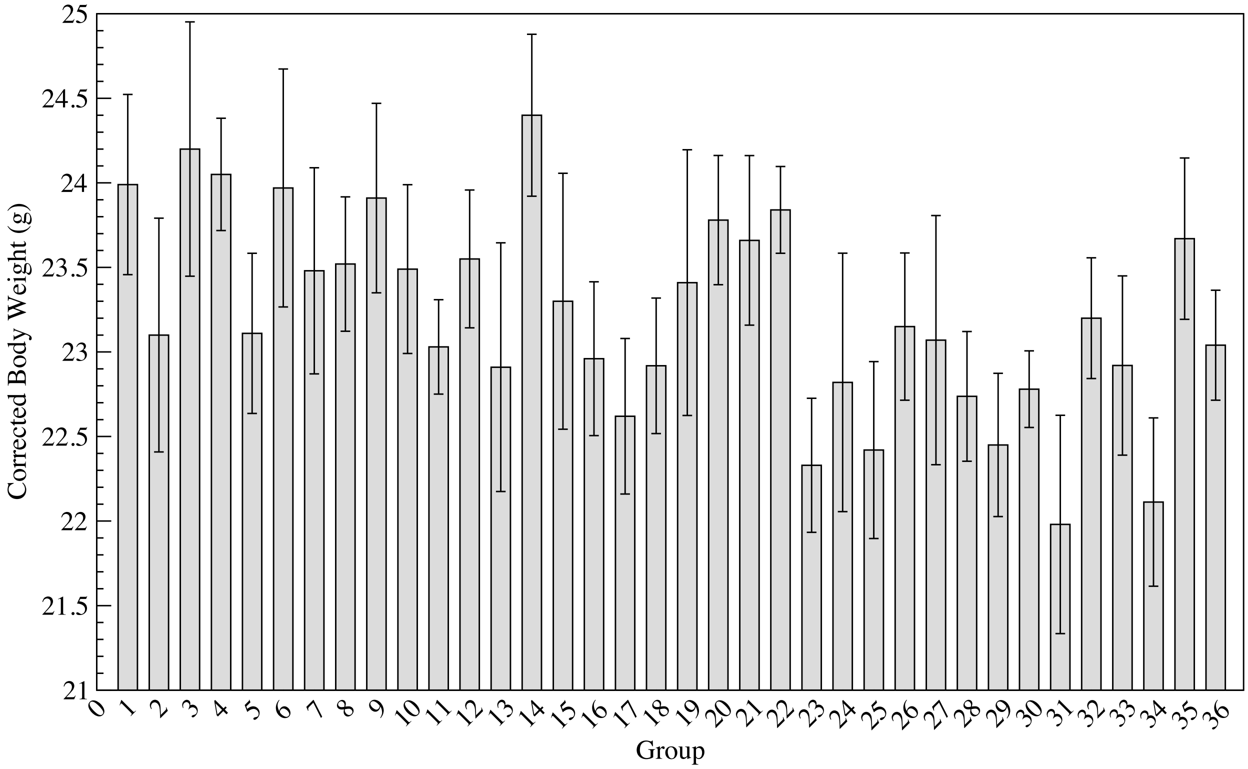


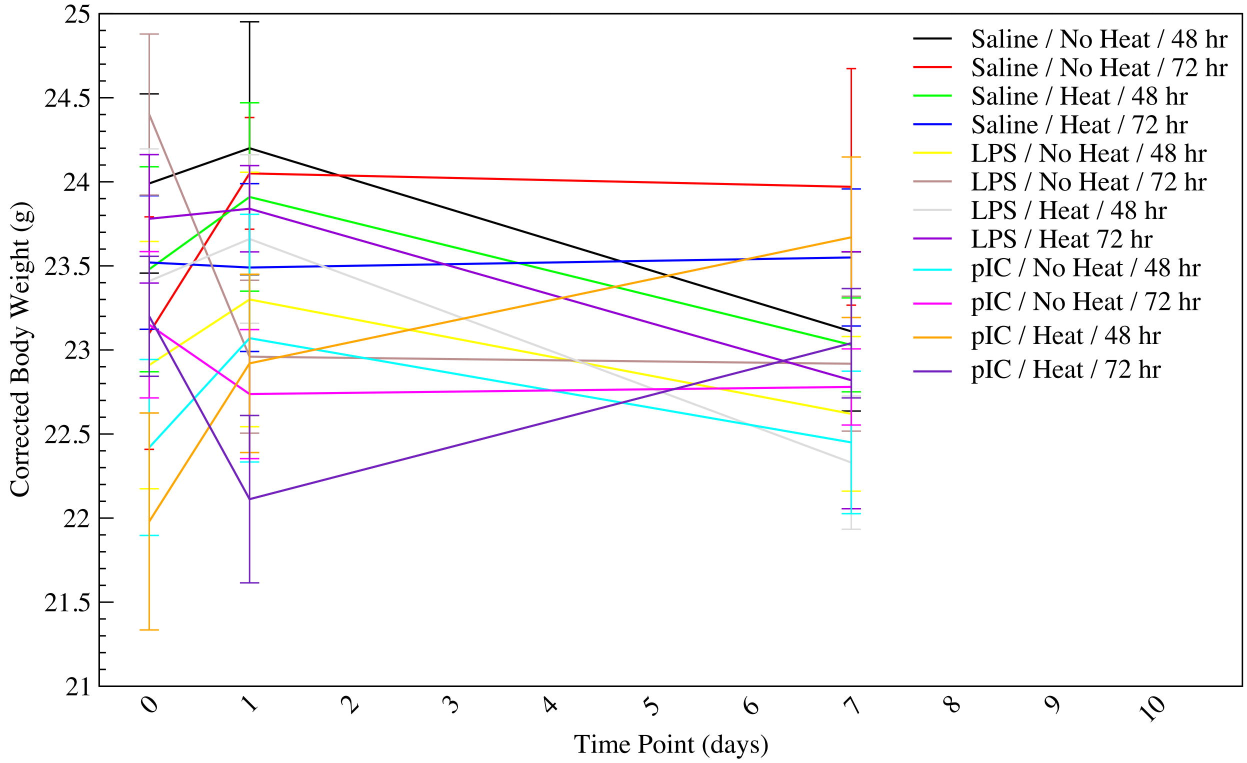

**D. Dehydration** is calculated as the percent change in body weight between pre-heat and $T_{c,max}$ time points. Heat challenged animals at all time points and treatments were significantly more dehydrated than non-heated animals.

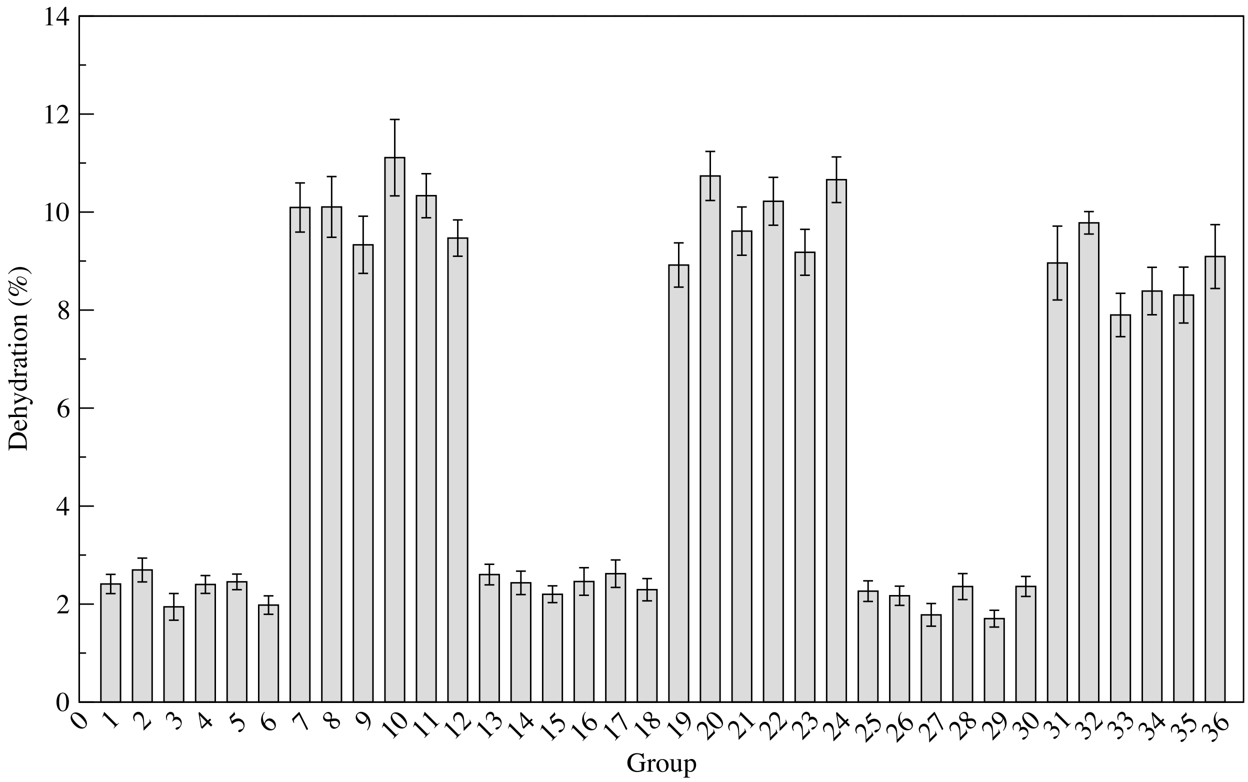


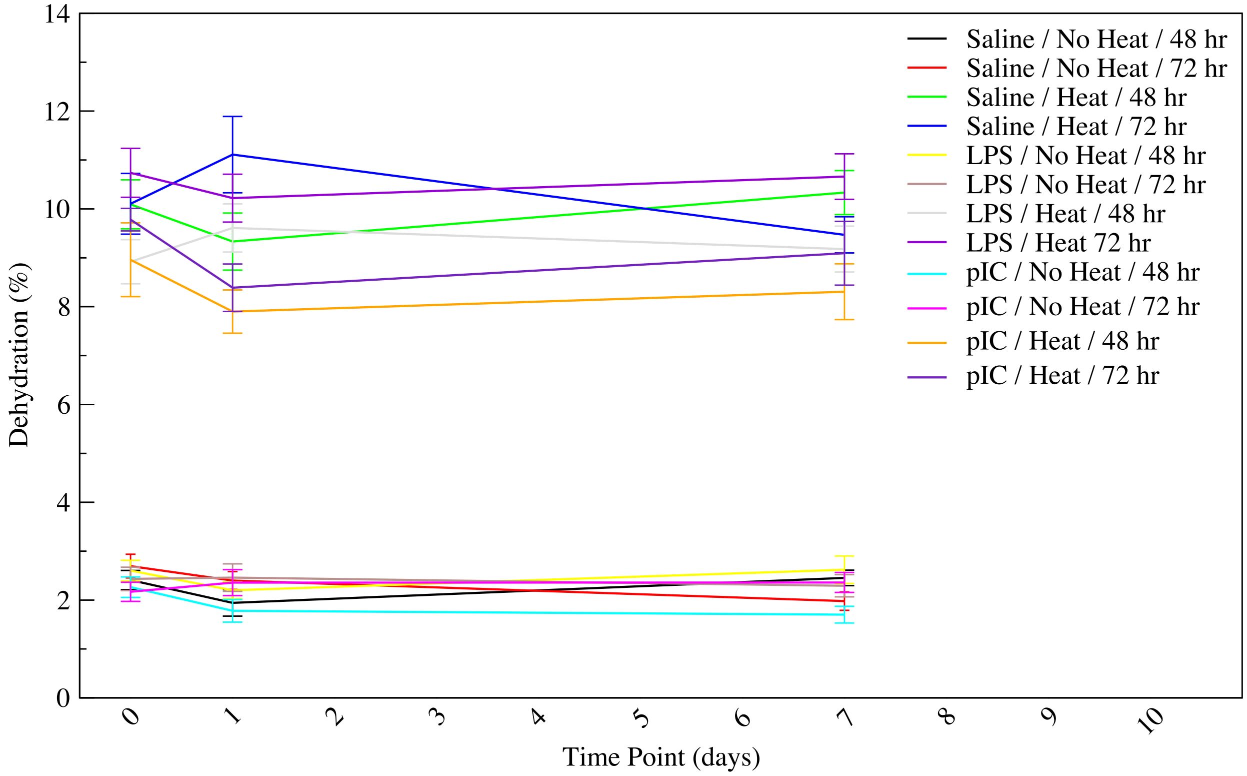

**E. White blood cell concentration** is measured in millions of cells per milliliter, and hence is influenced by blood volume and dehydration. Heated animals, regardless of treatment, exhibit significantly lower white blood cell concentration, indicating severe decreases that more than overcome the artificial increase that would be seen from dehydration. Animals mostly recover white blood cell concentration by the 7 day time point. Recovery is slower in animals treated with pIC, in agreement with the more severe heat stroke phenotype.

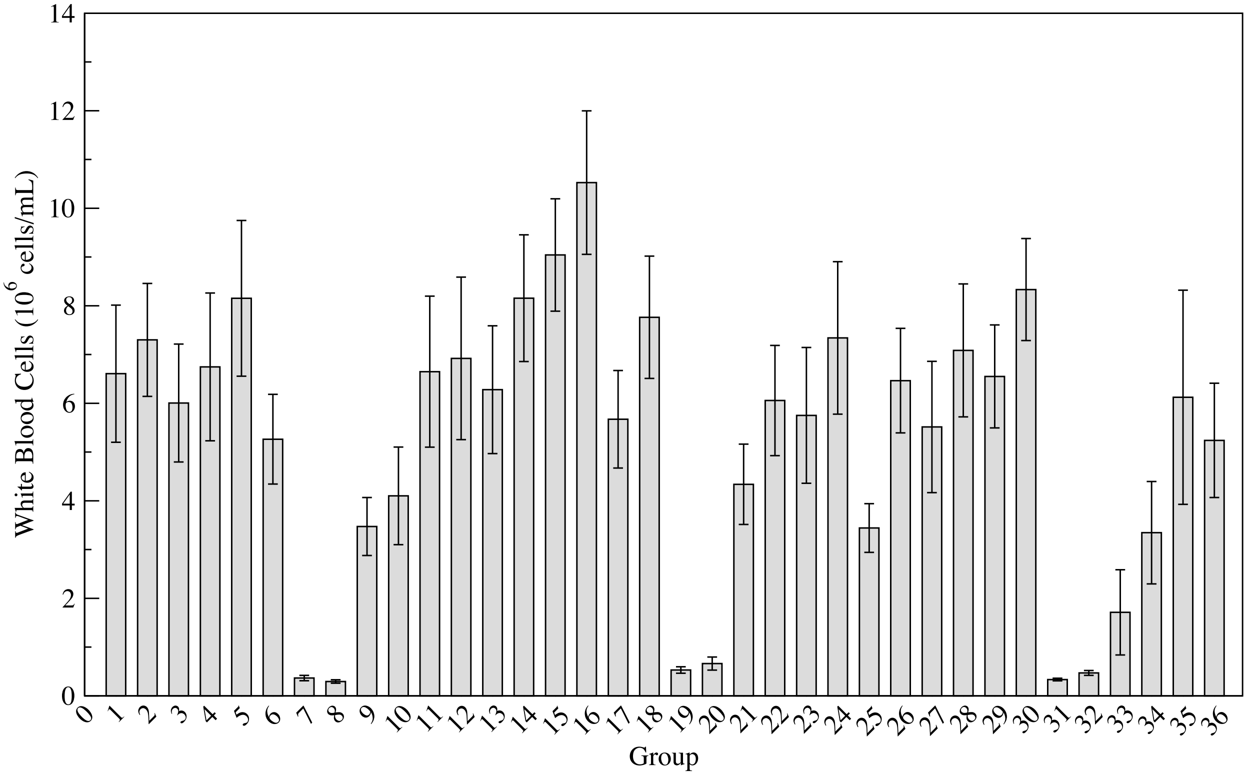


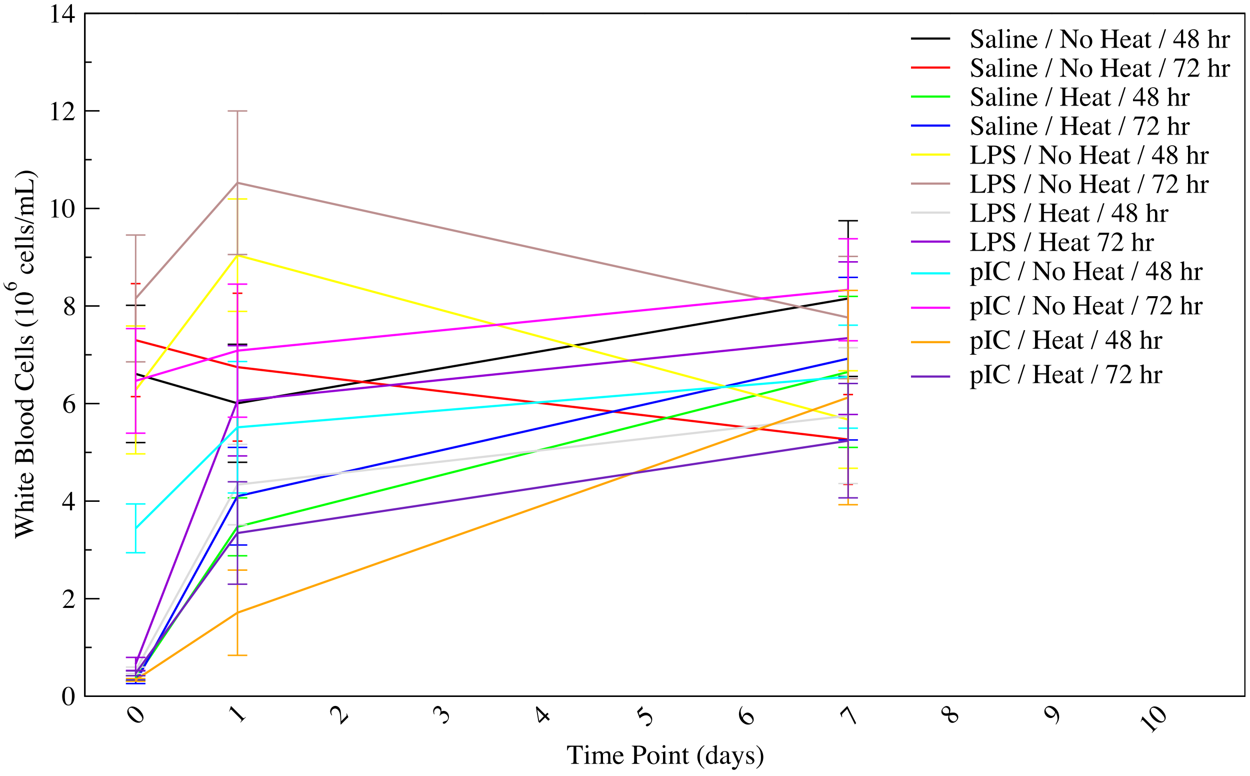

**F. Lymphocyte concentration** is measured in millions of cells per milliliter, and hence is influenced by blood volume and dehydration. Heated animals, regardless of treatment, exhibit significantly lower lymphocyte concentration, indicating severe decreases that more than overcome the artificial increase that would be seen from dehydration. Animals mostly recover lymphocyte concentration by the 7 day time point. Recovery is slower in animals treated with pIC, in agreement with the more severe heat stroke phenotype. Unheated LPS and pIC treated animals also experience an initial decrease in lymphocyte concentration, although this decrease is less statistically significant.

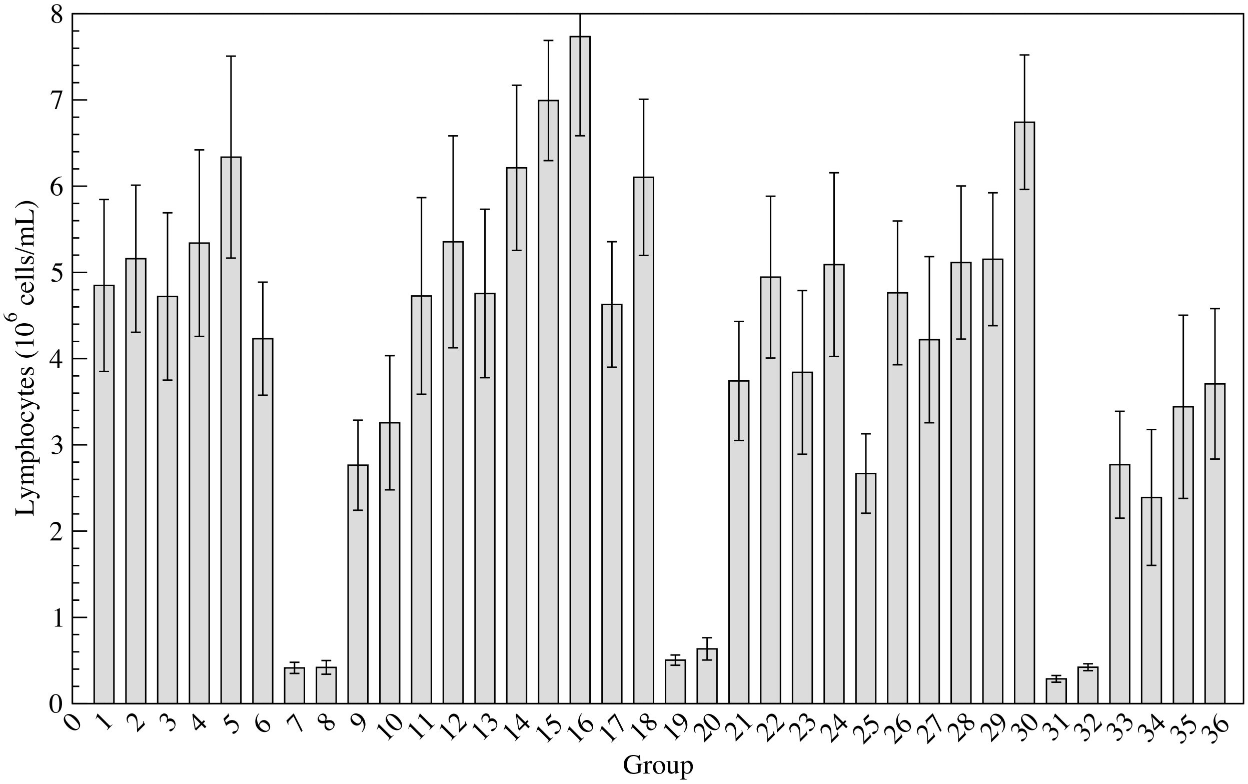


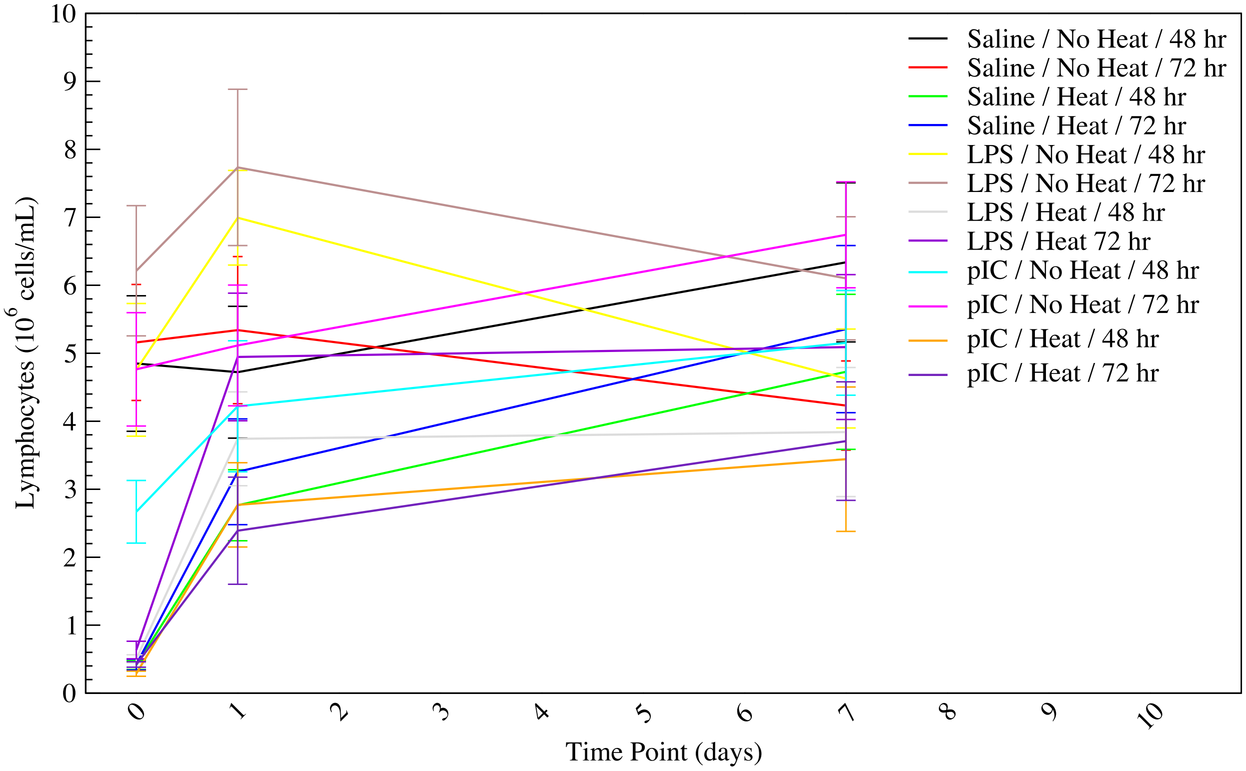

**G. Monocyte concentration** is measured in millions of cells per milliliter, and hence is influenced by blood volume and dehydration. Heated animals, regardless of treatment, exhibit significantly lower monocyte concentration, indicating severe decreases that more than overcome the artificial increase that would be seen from dehydration. Animals mostly recover monocyte concentration by the 7 day time point. Recovery is slower in animals treated with pIC, in agreement with the more severe heat stroke phenotype.

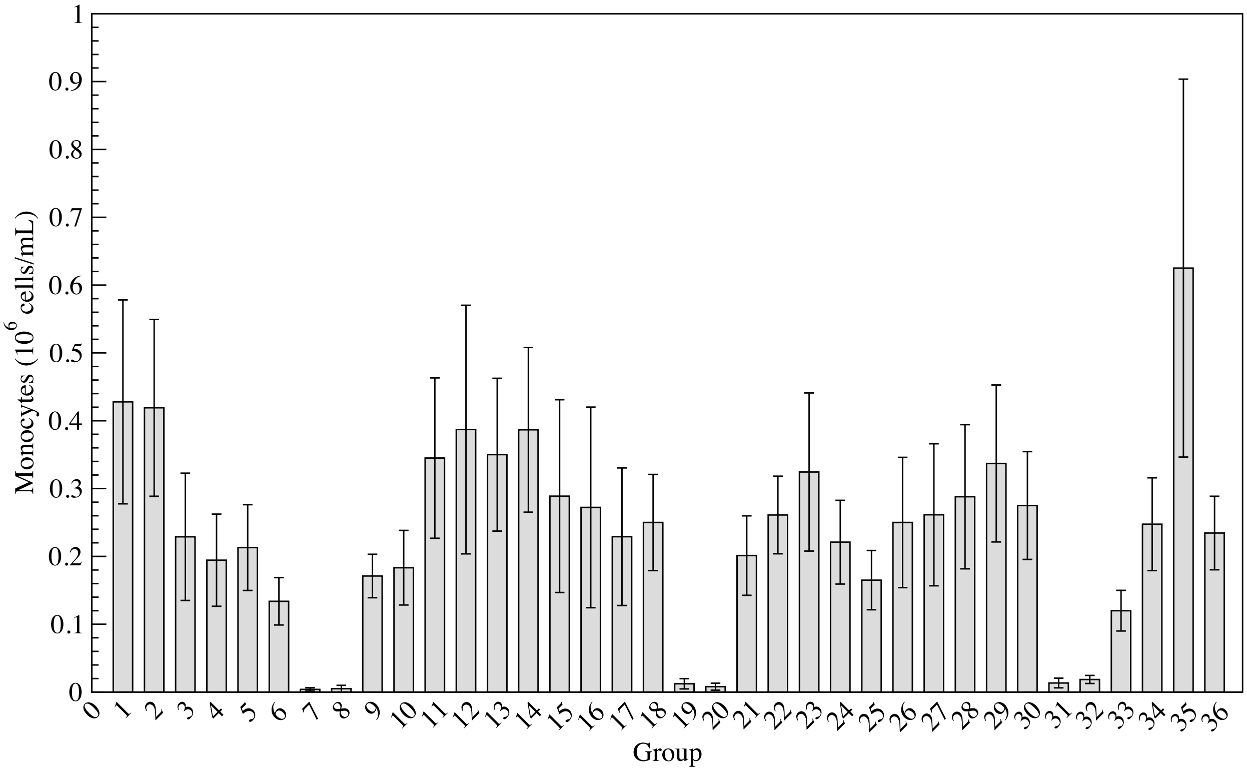


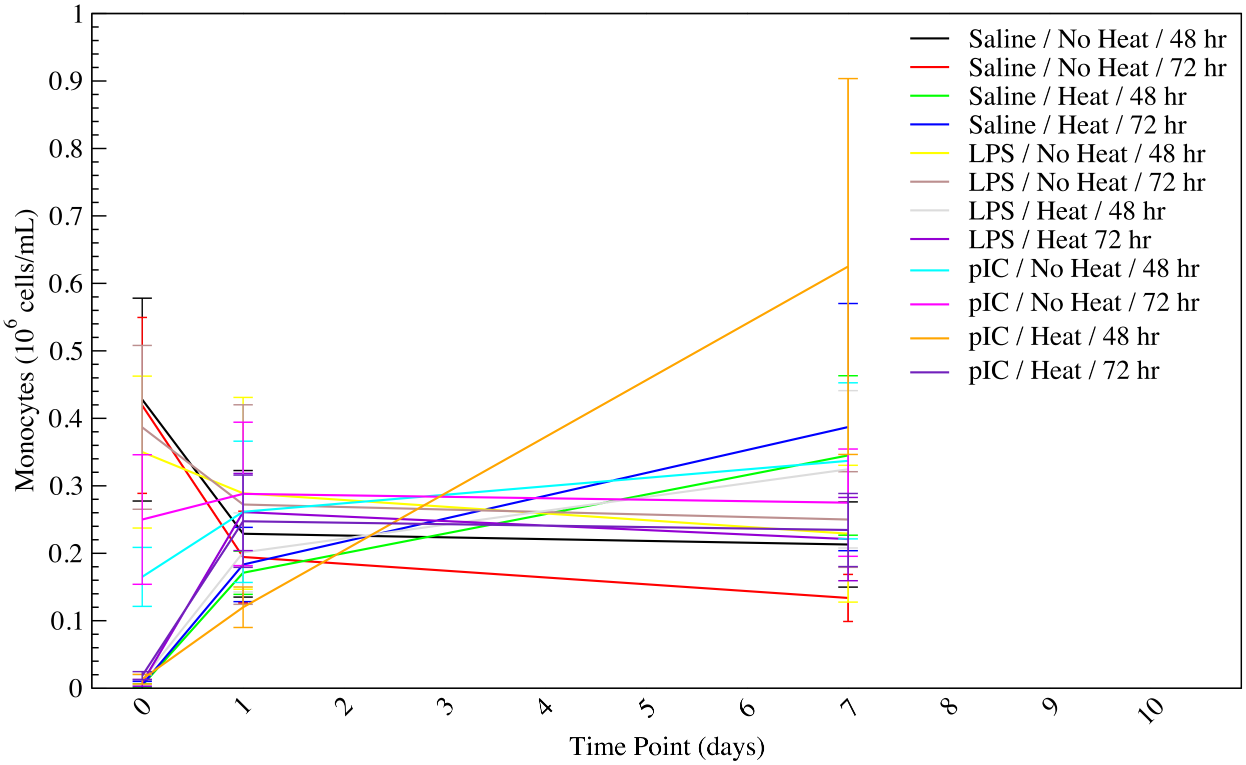

**H. Neutrophil concentration** is measured in millions of cells per milliliter, and hence is influenced by blood volume and dehydration. Heated animals, regardless of treatment, exhibit significantly lower neutrophil concentration, indicating severe decreases that more than overcome the artificial increase that would be seen from dehydration. Animals mostly recover neutrophil concentration by the 7 day time point.

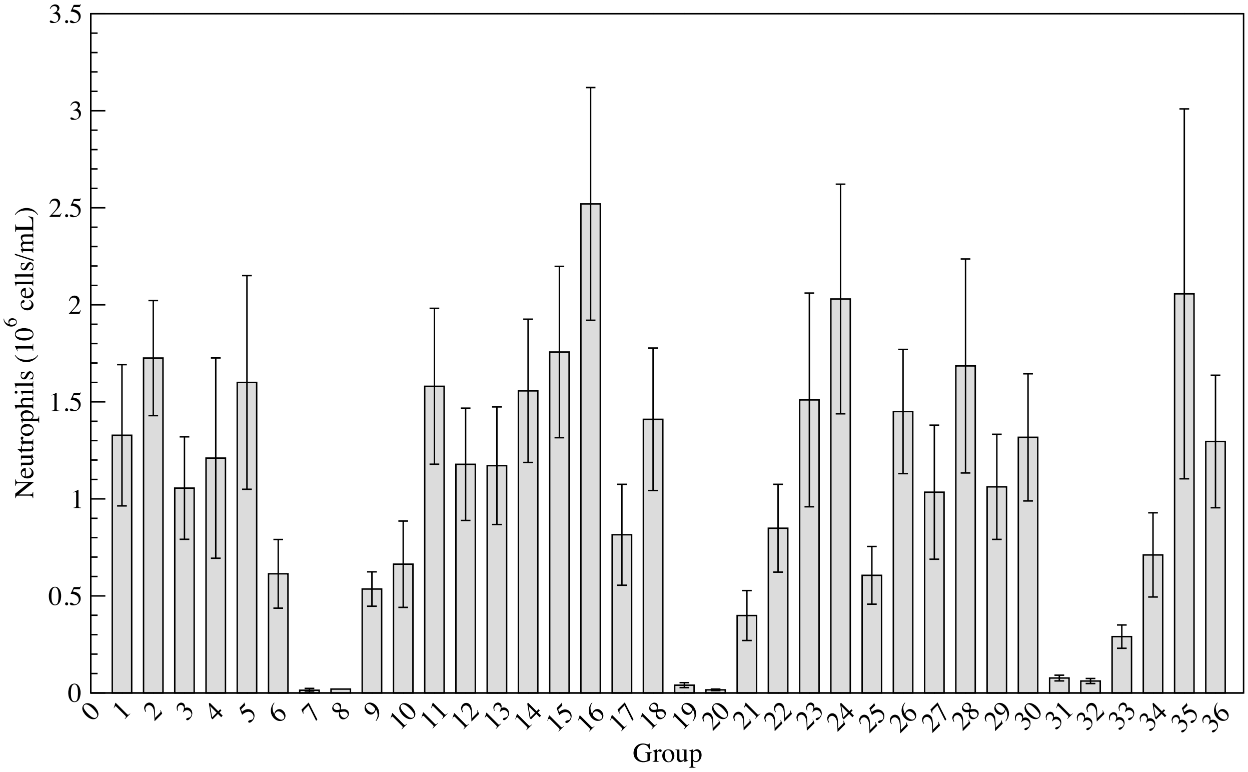


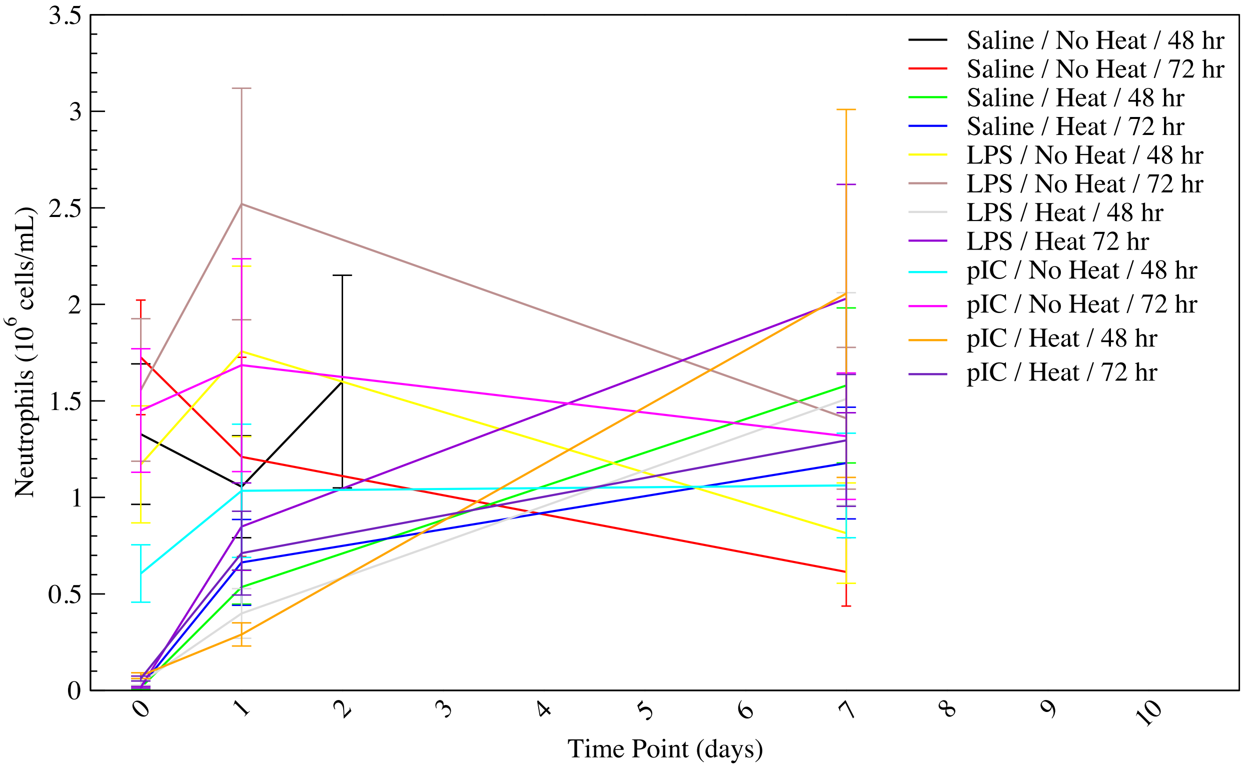

**I. Lymphocyte fraction** is measured as a percentage of the white blood cell population, and therefore (unlike concentration) is not influenced by blood volume or dehydration but is influenced by relative abundance of other white blood cell populations. In the saline and LPS treatment groups, lymphocyte fraction is elevated at $T_{c,max}$ in the heated animals, while pIC treated animals experienced a drop in lymphocyte fraction at later timepoints.

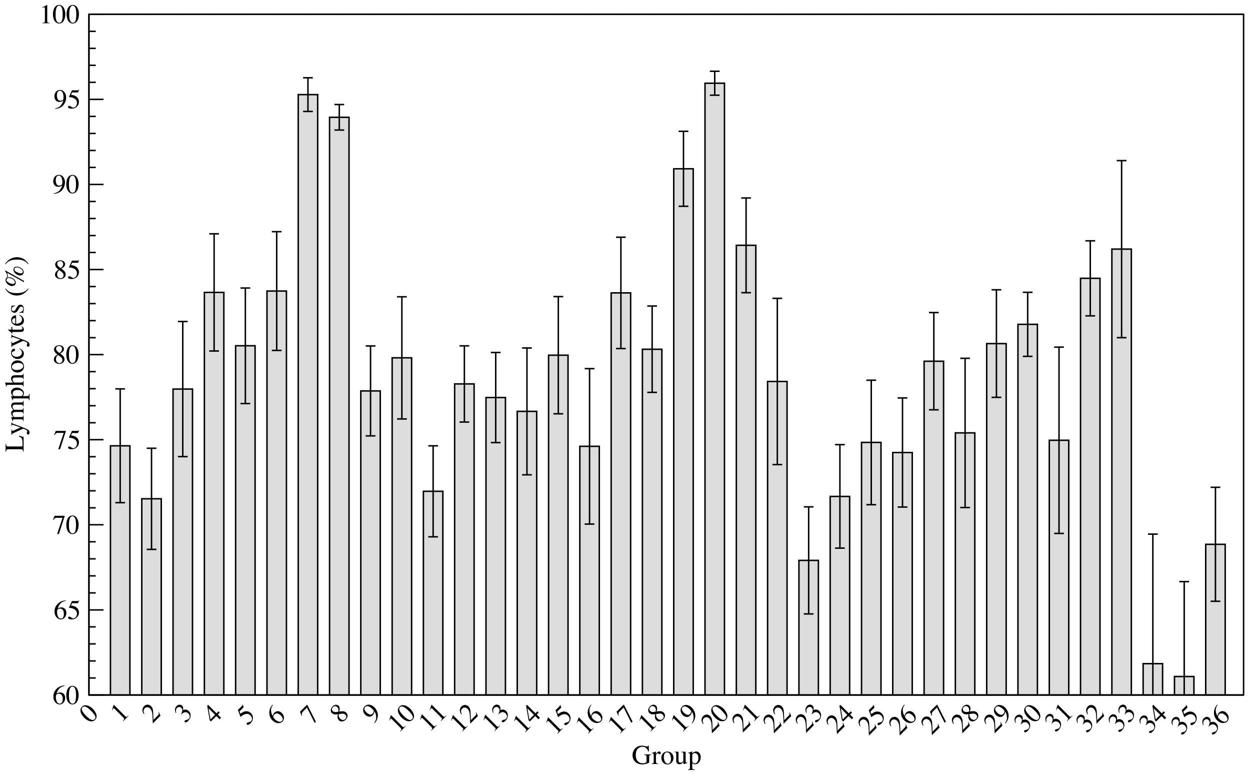


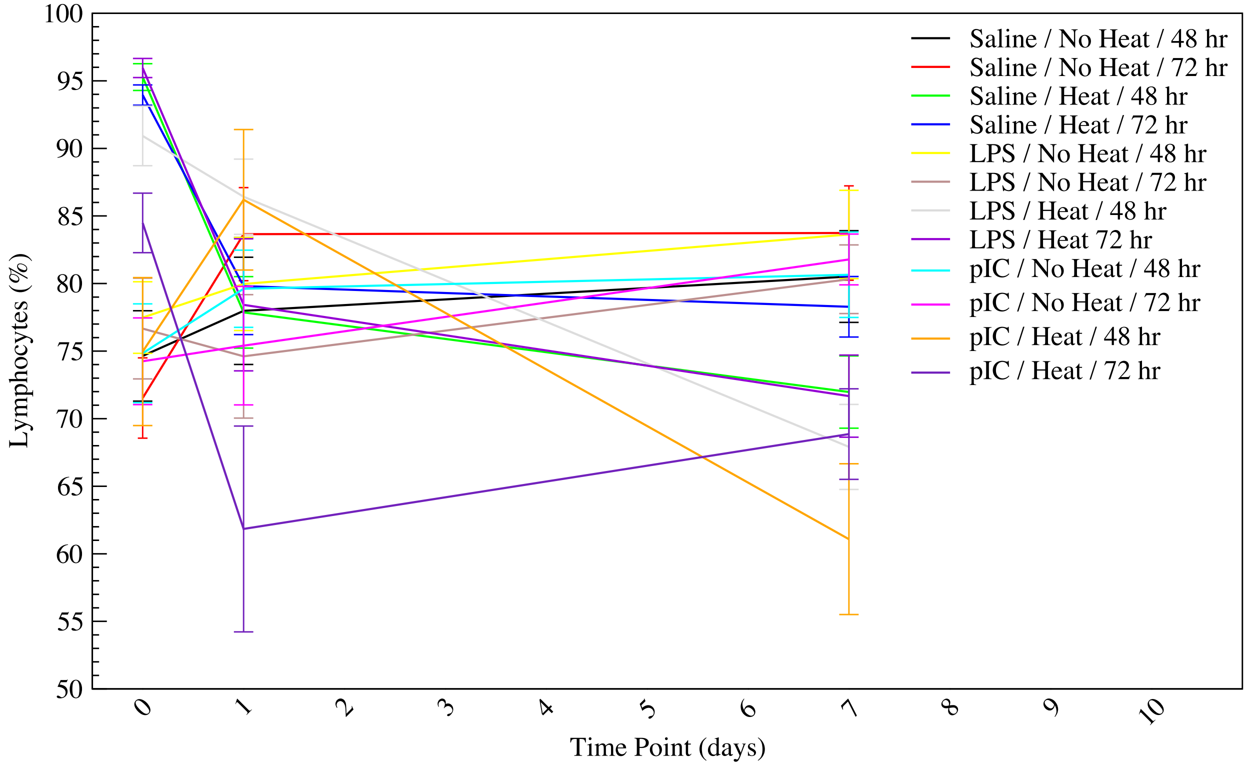

**J. Monocyte fraction** is measured as a percentage of the white blood cell population, and therefore (unlike concentration) is not influenced by blood volume or dehydration but is influenced by relative abundance of other white blood cell populations. In the saline and LPS treatment groups, monocyte fraction is decreased at $T_{c,max}$ in the heated animals, while pIC treated animals experienced a rise in monocyte fraction at later timepoints. These trends are opposite the lymphocyte fraction trends noted above.

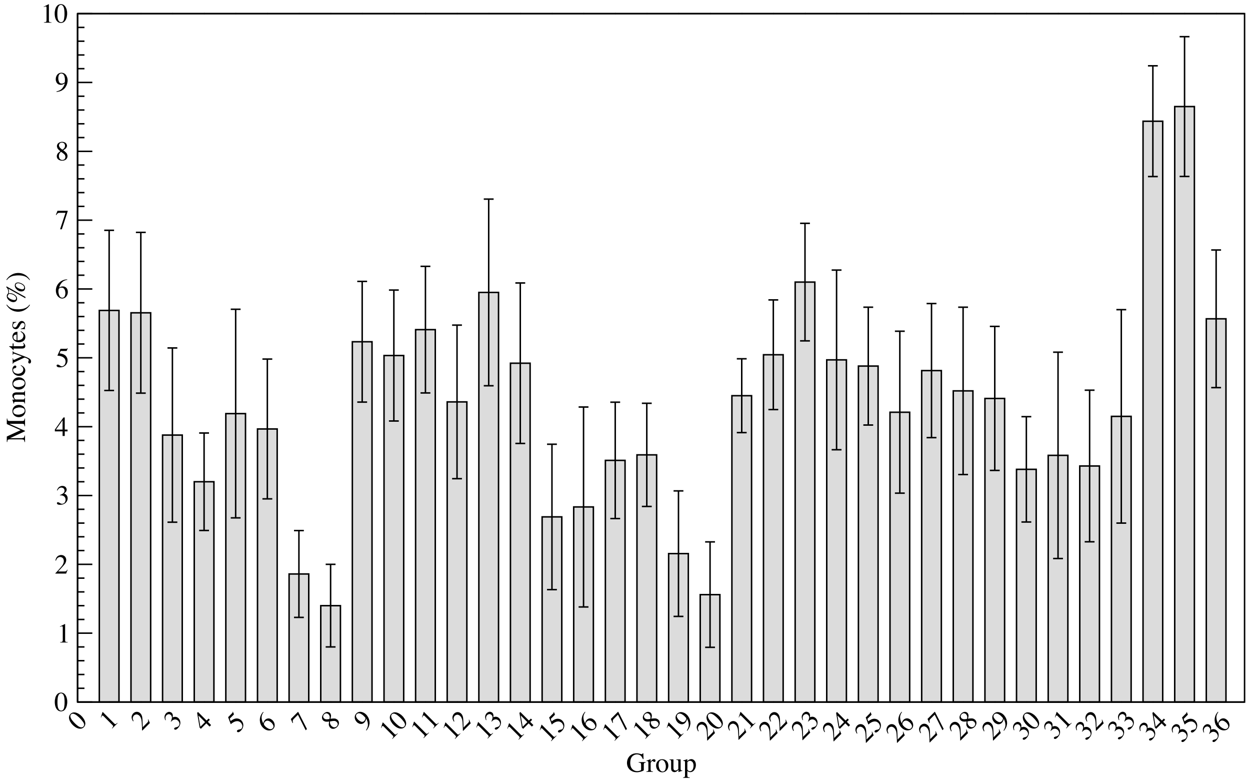


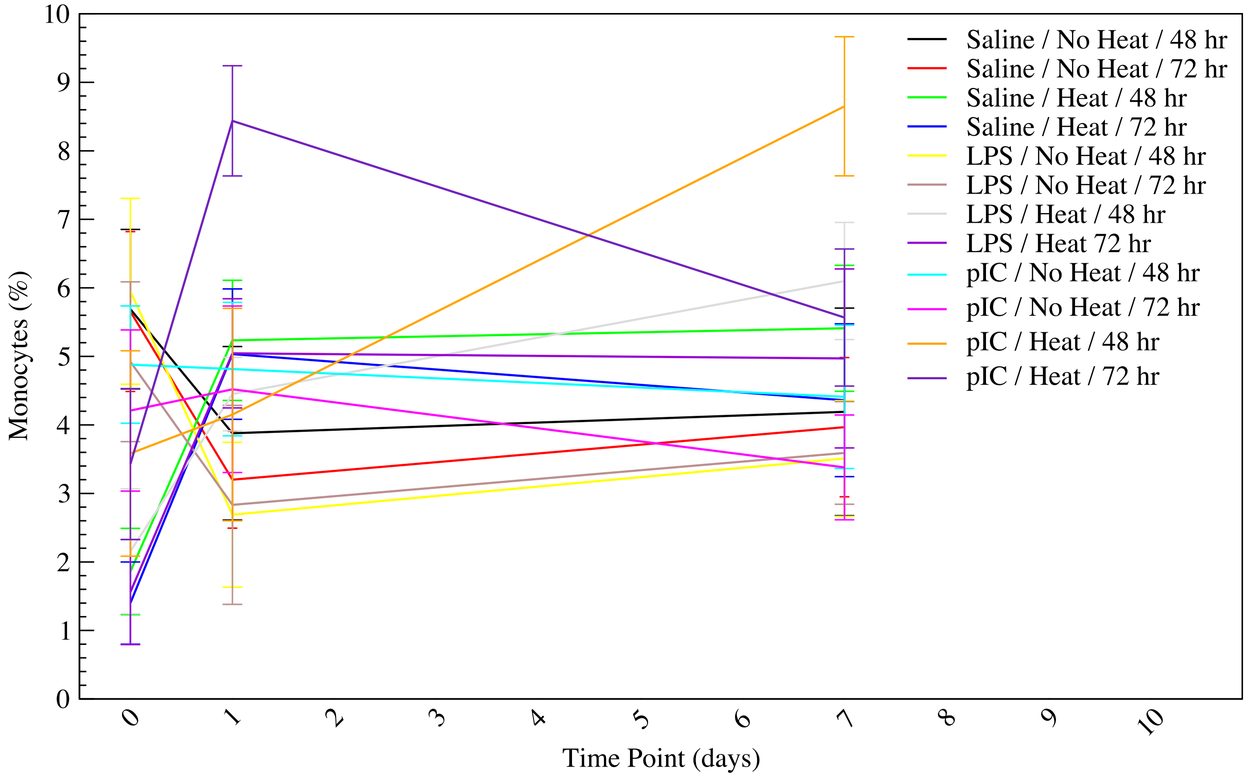

**K. Neutrophil fraction** is measured as a percentage of the white blood cell population, and therefore (unlike concentration) is not influenced by blood volume or dehydration but is influenced by relative abundance of other white blood cell populations. In the saline and LPS treatment groups, neutrophil fraction is decreased at $T_{c,max}$ in the heated animals, while pIC treated animals experienced a rise in neutrophil fraction at later timepoints. These trends match the monocyte trends and are opposite the lymphocyte fraction trends noted above.

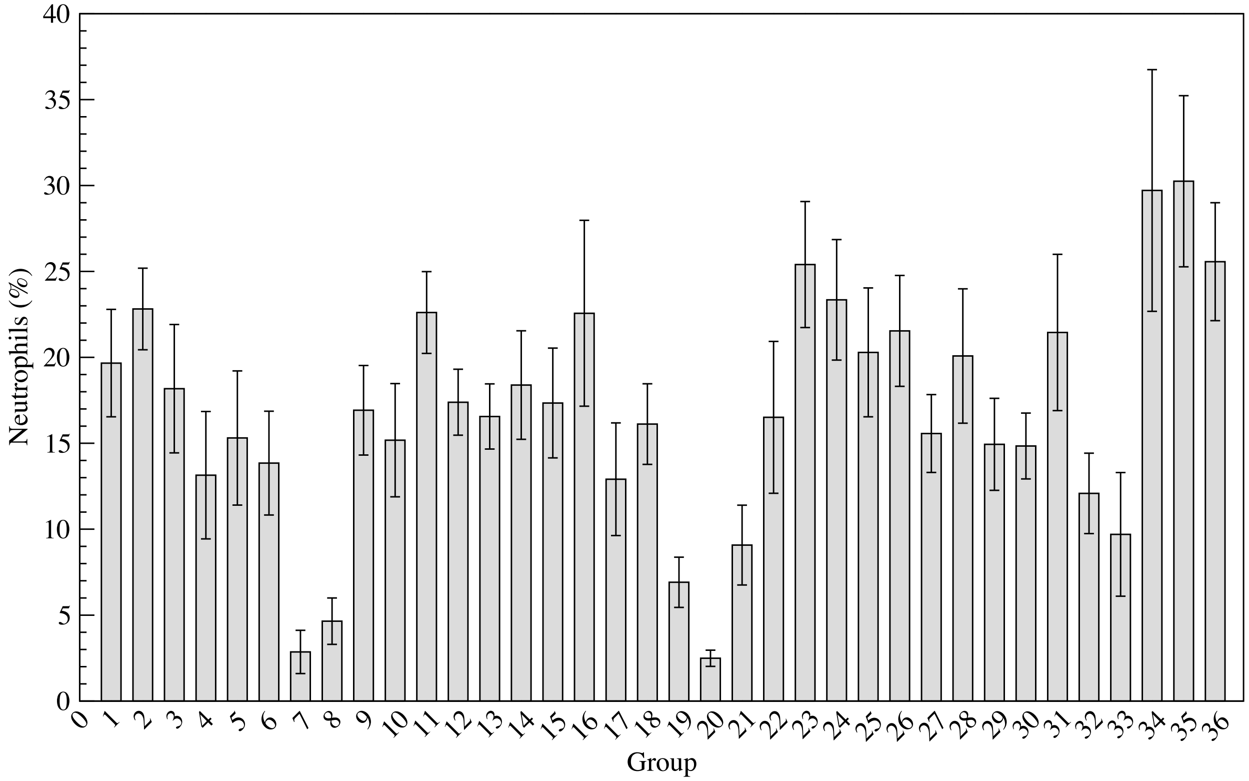


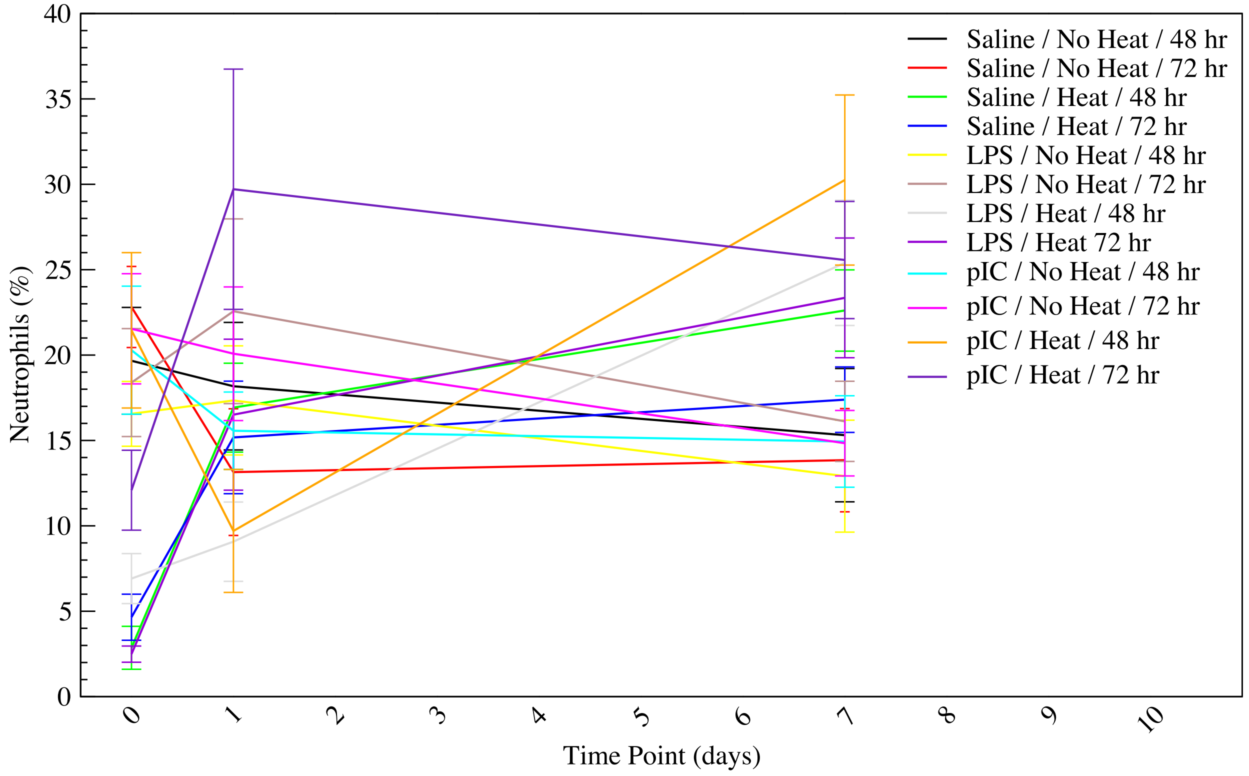

**L. Red blood cell concentration** is measured as billions of cells per milliliter, and hence is influenced by blood volume and dehydration. Heated animals, regardless of treatment, exhibit a spike in red blood cell concentration at $T_{c,max}$. In saline and LPS treated groups, this spike is followed by a severe drop in the 72 hour incubation at 1 day, while the 48 hour incubation group returns to baseline. In pIC treated animals, both incubation groups return to baseline by the 1 day time point.

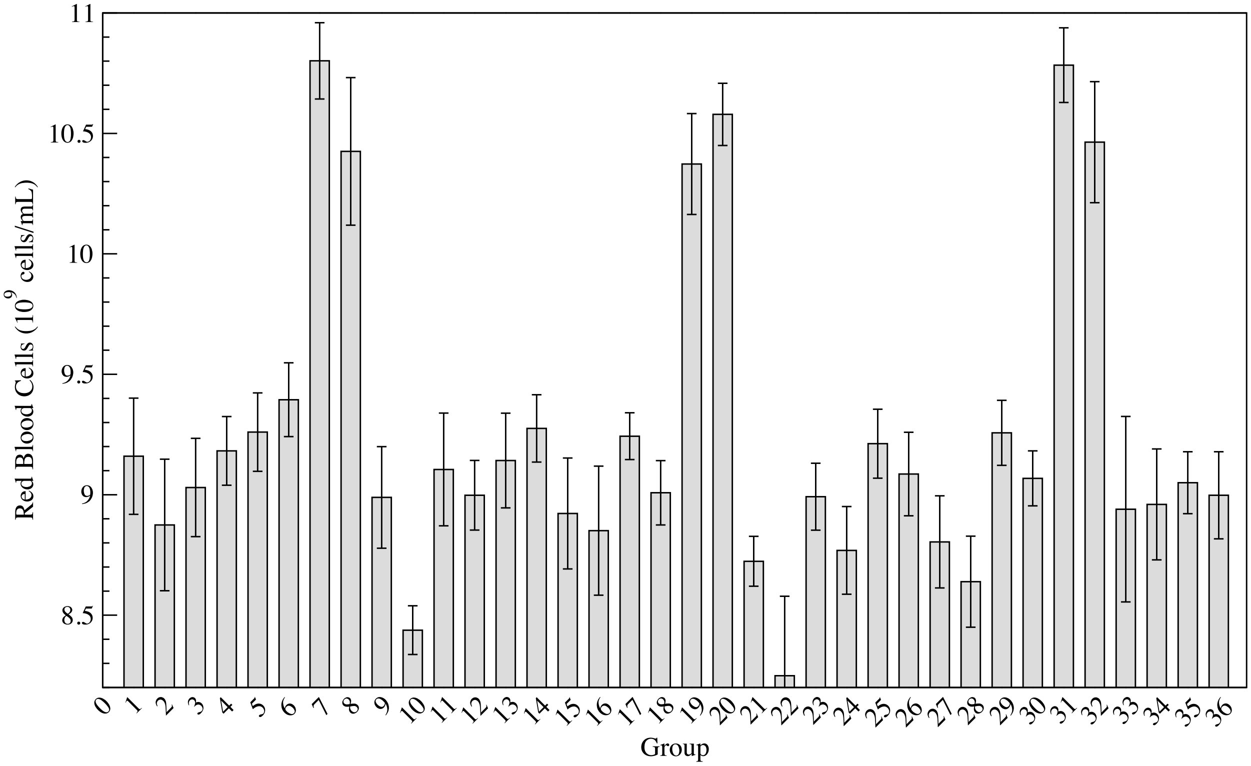


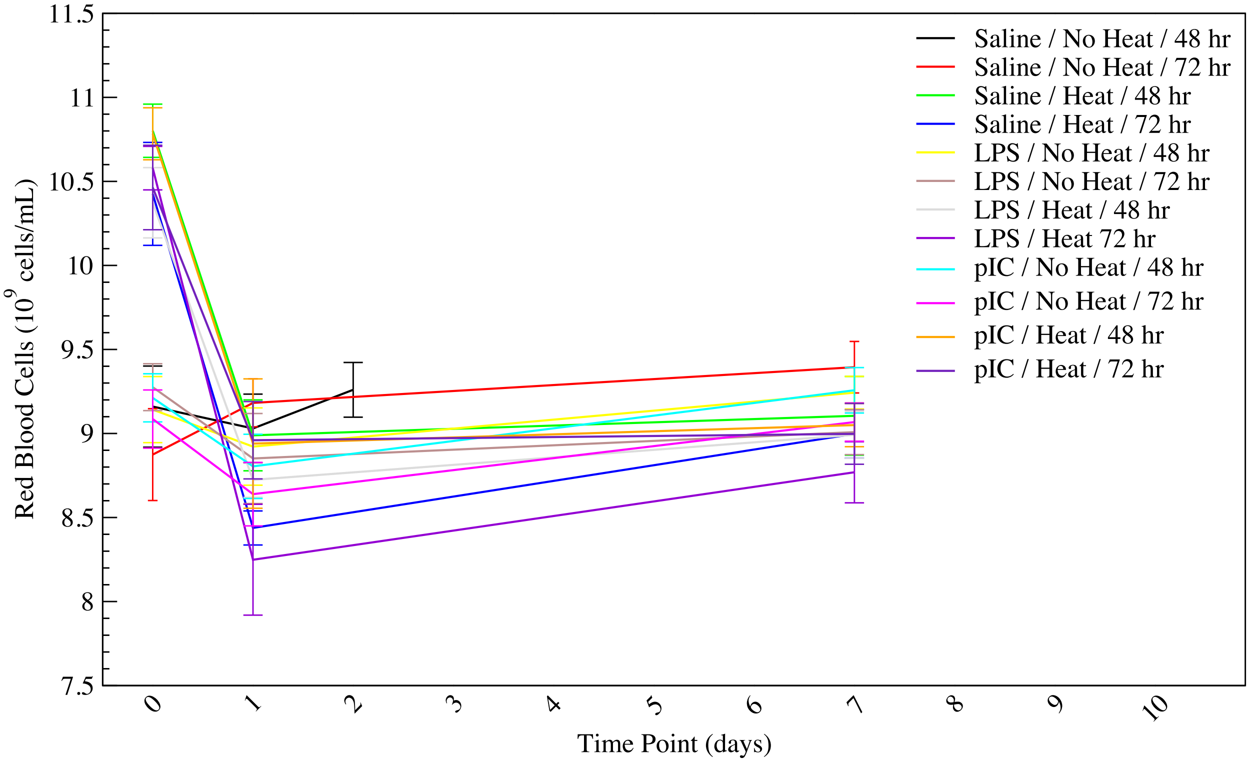

**M. Hemoglobin concentration** is measured as grams per deciliter, and hence is influenced by blood volume and dehydration. Hemoglobin concentration follows the same trends as red blood cell concentration.

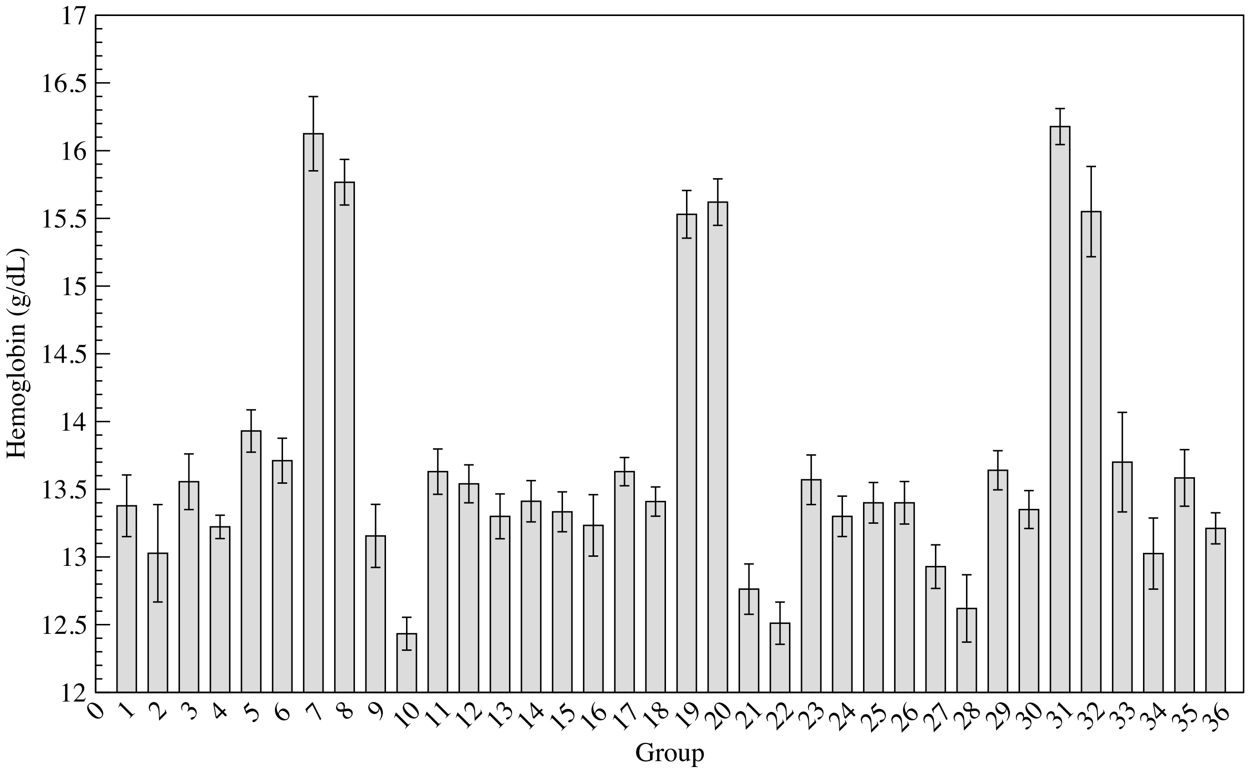


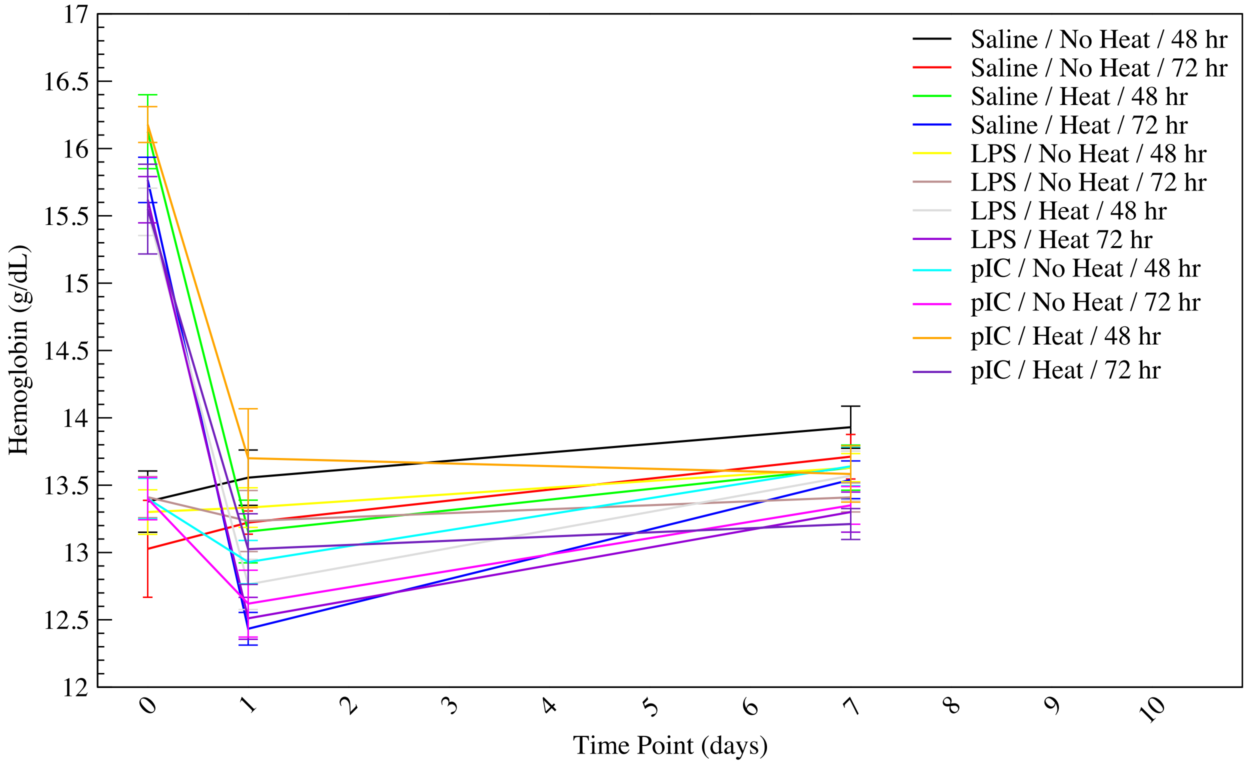

**N. Hematocrit fraction** is measured as the volume percentage of red blood cells in blood, and hence is influenced by blood volume and dehydration. Hematocrit follows the same trends as red blood cell concentration.

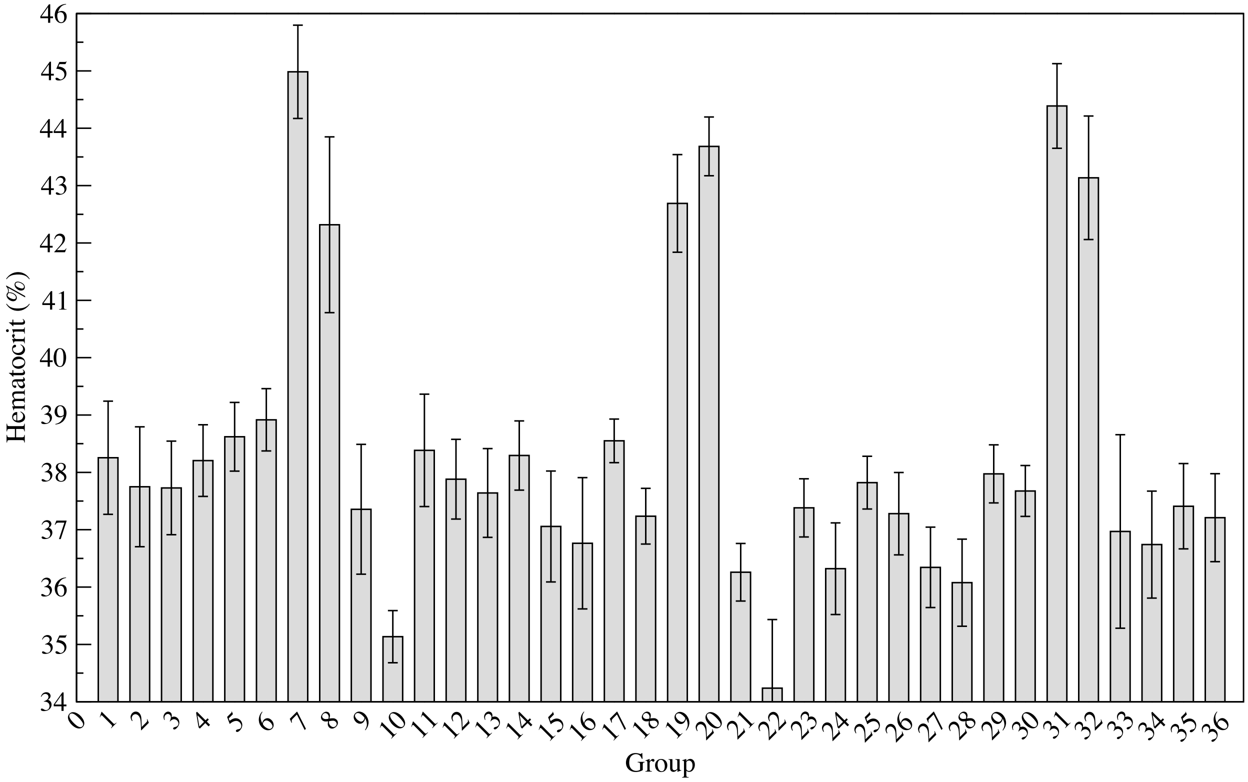


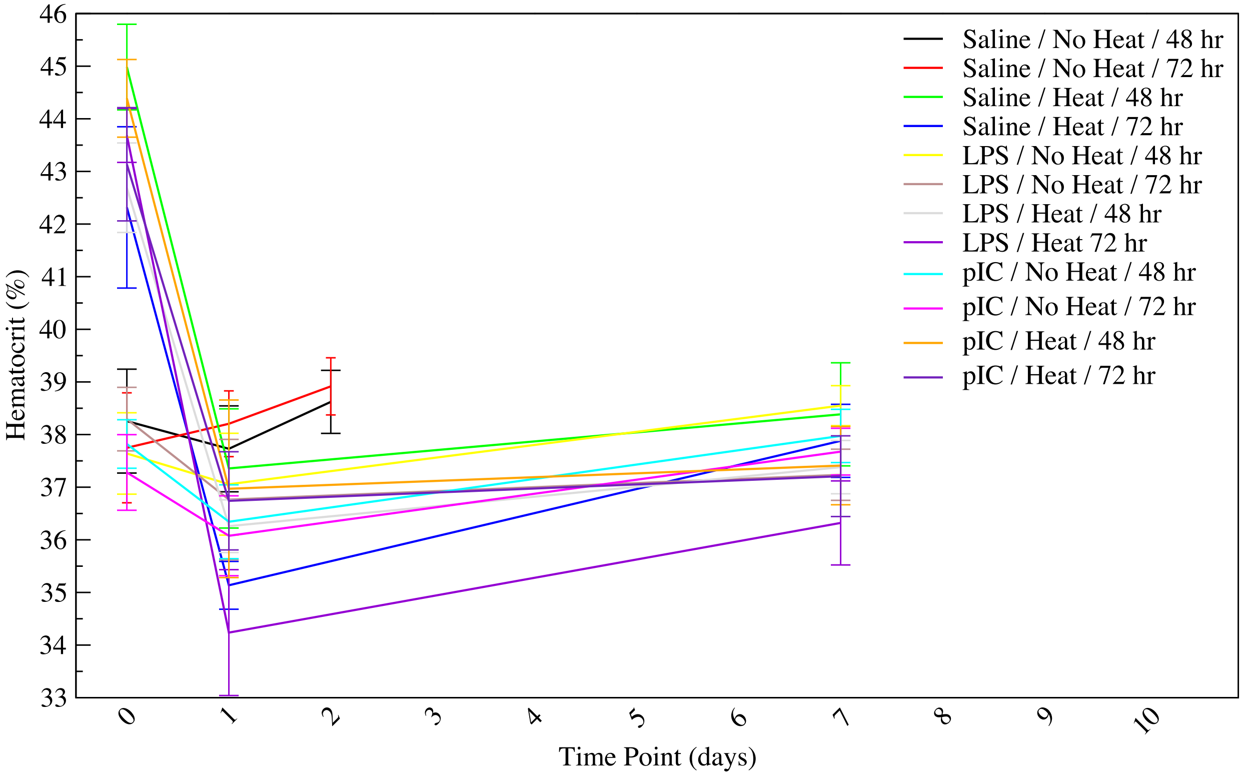

**O. Mean corpuscular volume** is the average volume of a red blood cell, measured in femtoliters, and is the quotient of the hematocrit and the red blood cell concentration. Mean corpuscular volume slightly differs over treatment groups, as saline > LPS > pIC, but measurements were highly variable and differences not statistically significant.

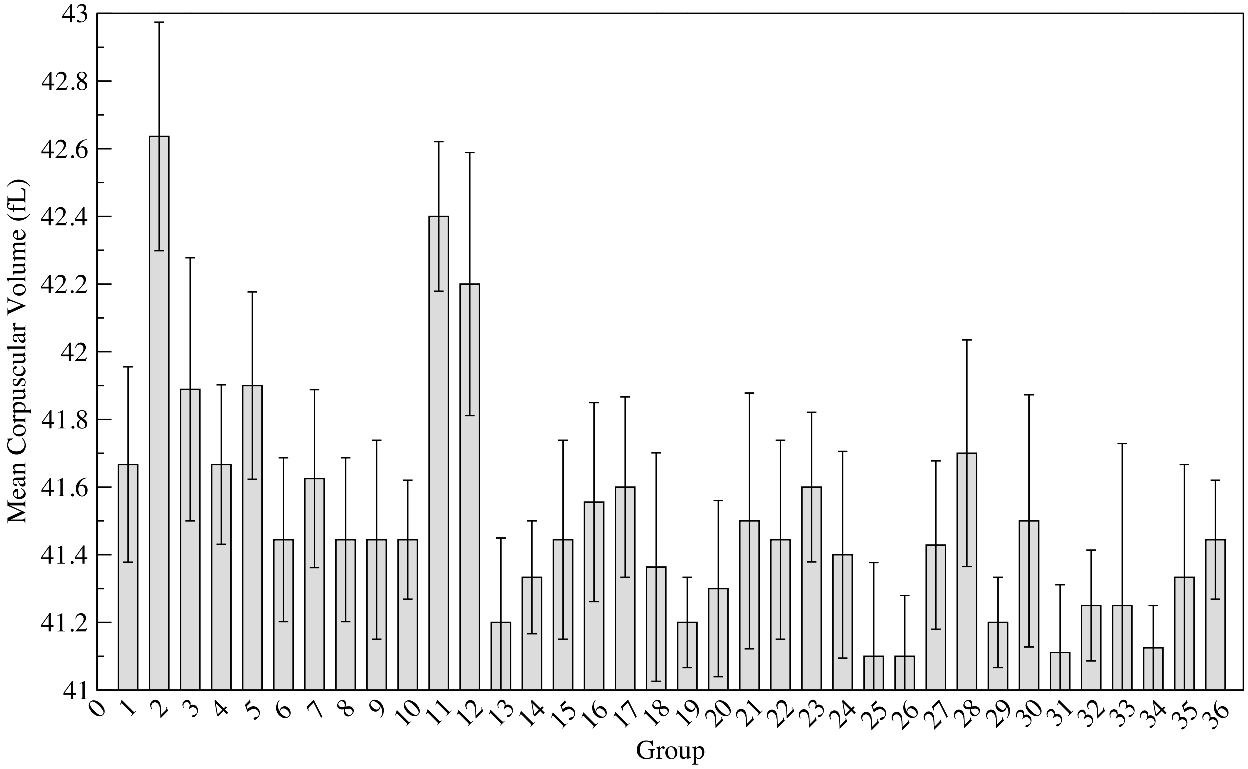


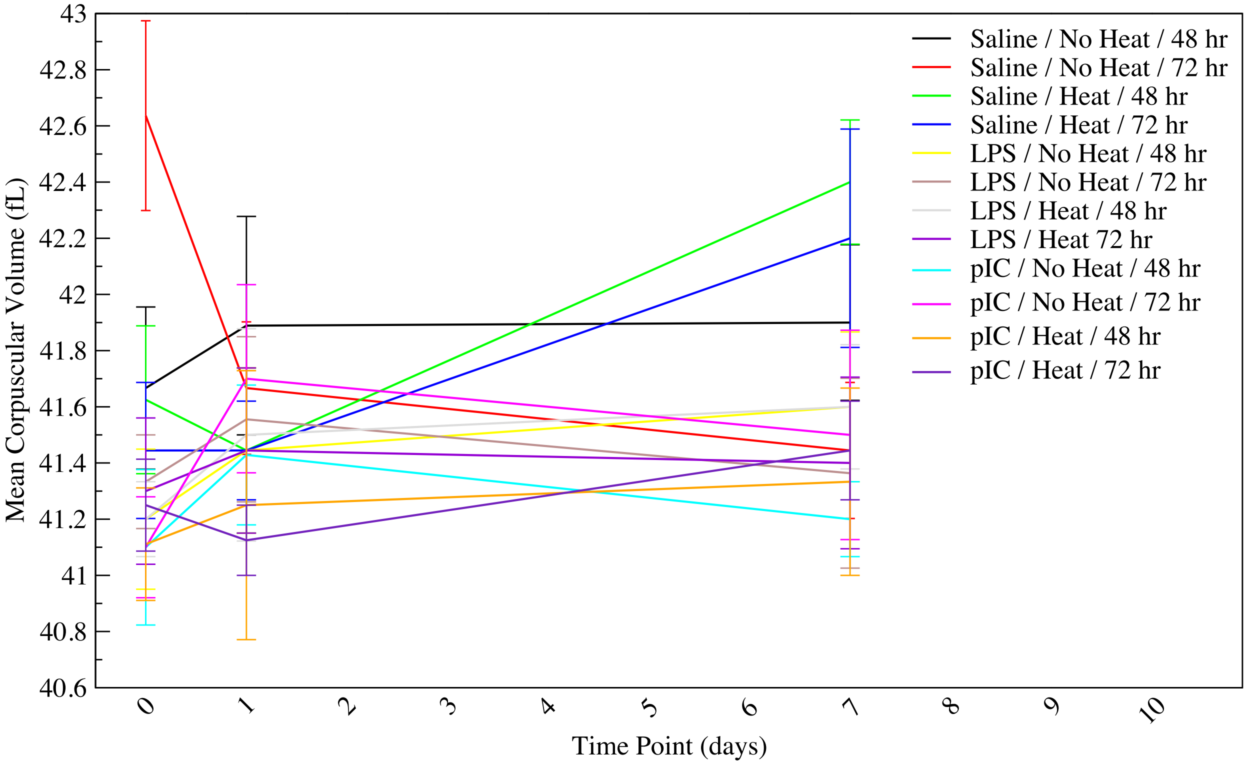

**P. Mean corpuscular hemoglobin mass** is measured as the average mass in picograms of hemoglobin per red blood cell, and is proportional to the quotient of hemoglobin concentration and red blood cell concentration. No discernible patterns emerge.

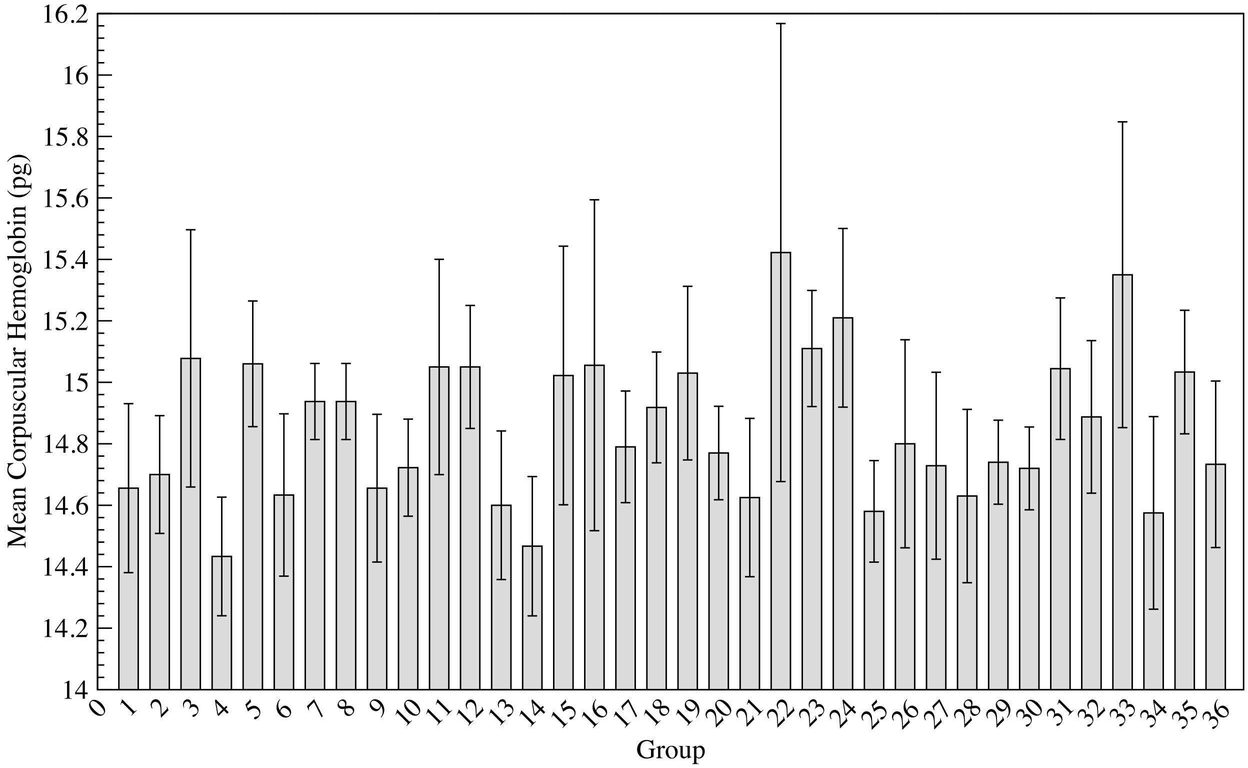


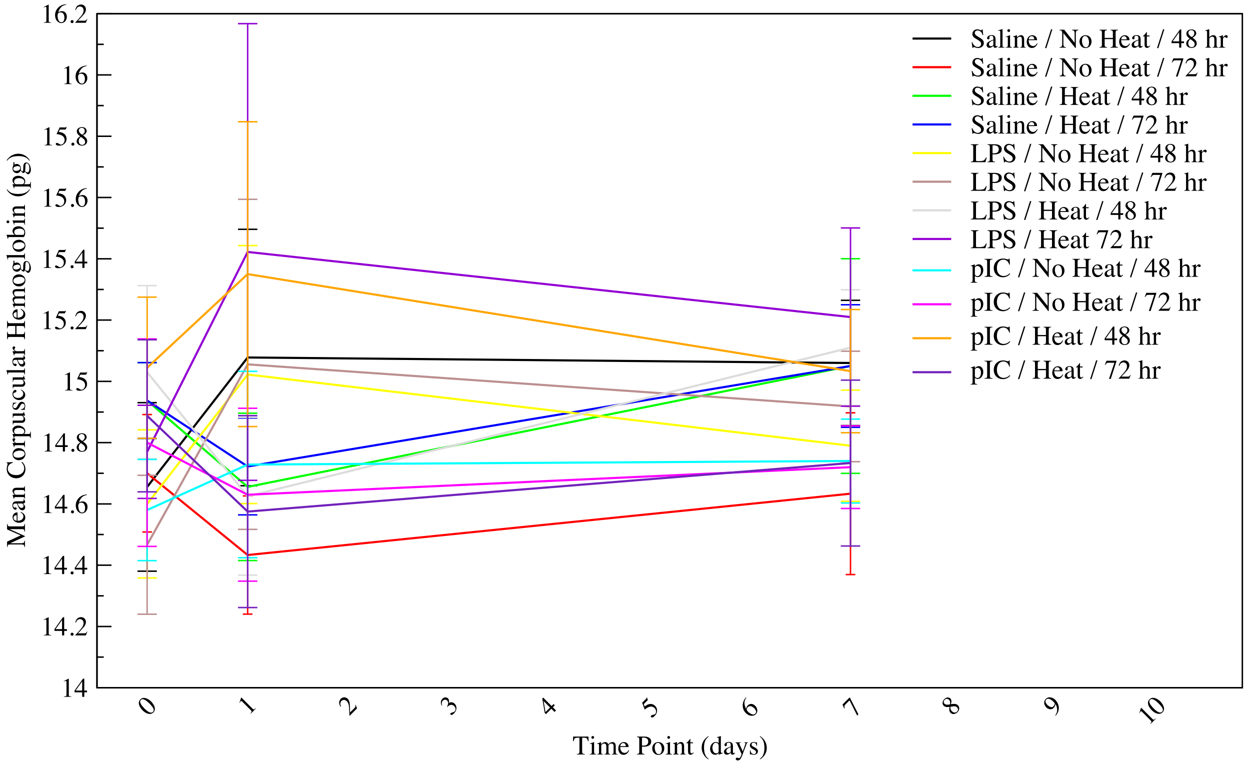

**Q. Mean corpuscular hemoglobin concentration** is measured as the average mass of hemoglobin per volume of red blood cell, as grams per deciliter, and is the quotient of the hemoglobin concentration and the hematocrit fraction. No discernible patterns emerge.

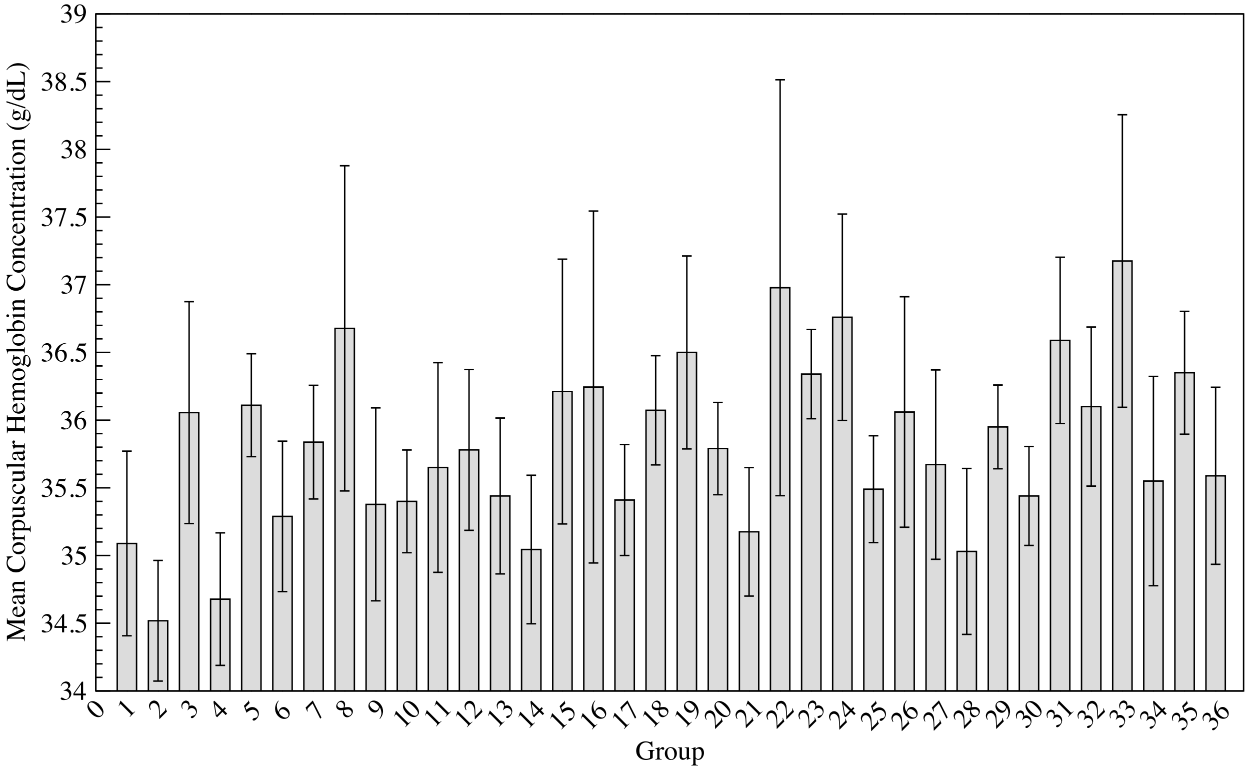


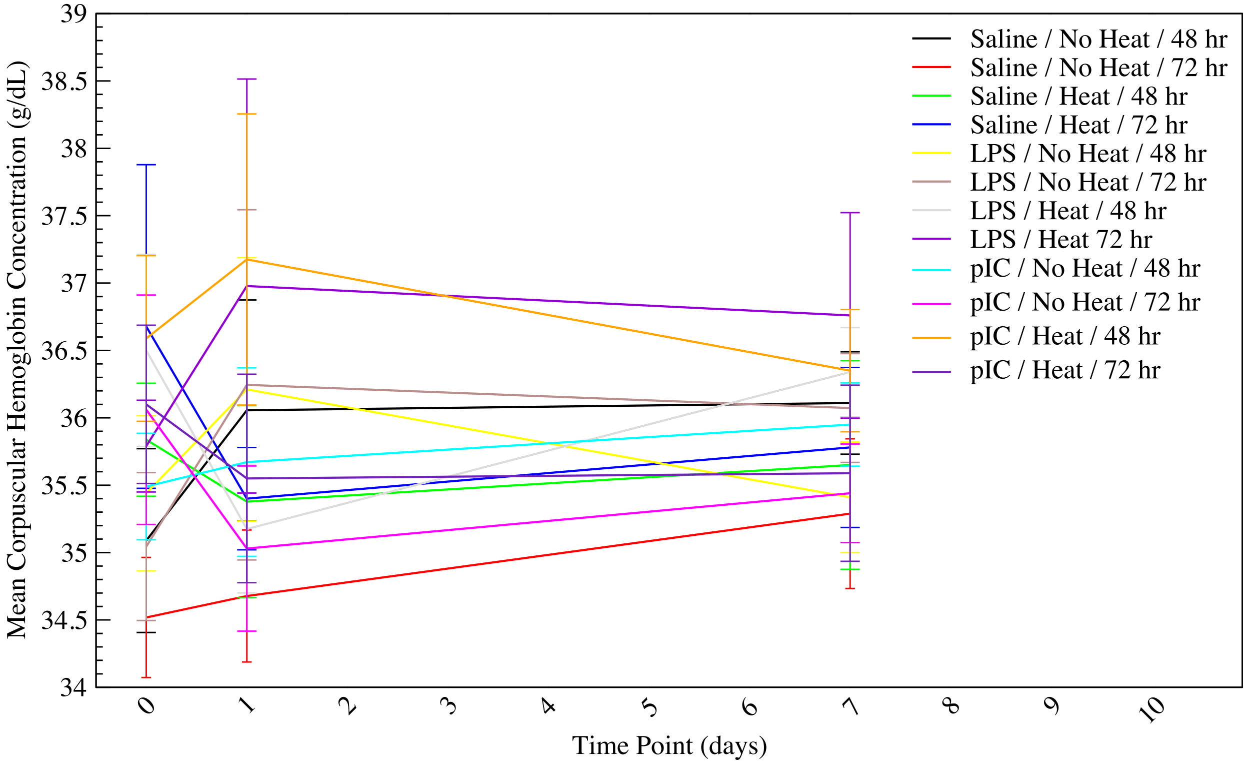

**R. Red blood cell size distribution coefficient of variation** is a measure of the range of variation of red blood cell volume. Heated animals in all treatment groups exhibited a narrowing of the red blood cell size distribution at the 1 day time point, followed by recovery by 7 days. In both heated and unheated pIC treated animals, this narrowing was already present at $T_{c,max}$. Combined with mean corpuscular volume measurements, above, the change in size distribution was not accompanied by a change in mean size.

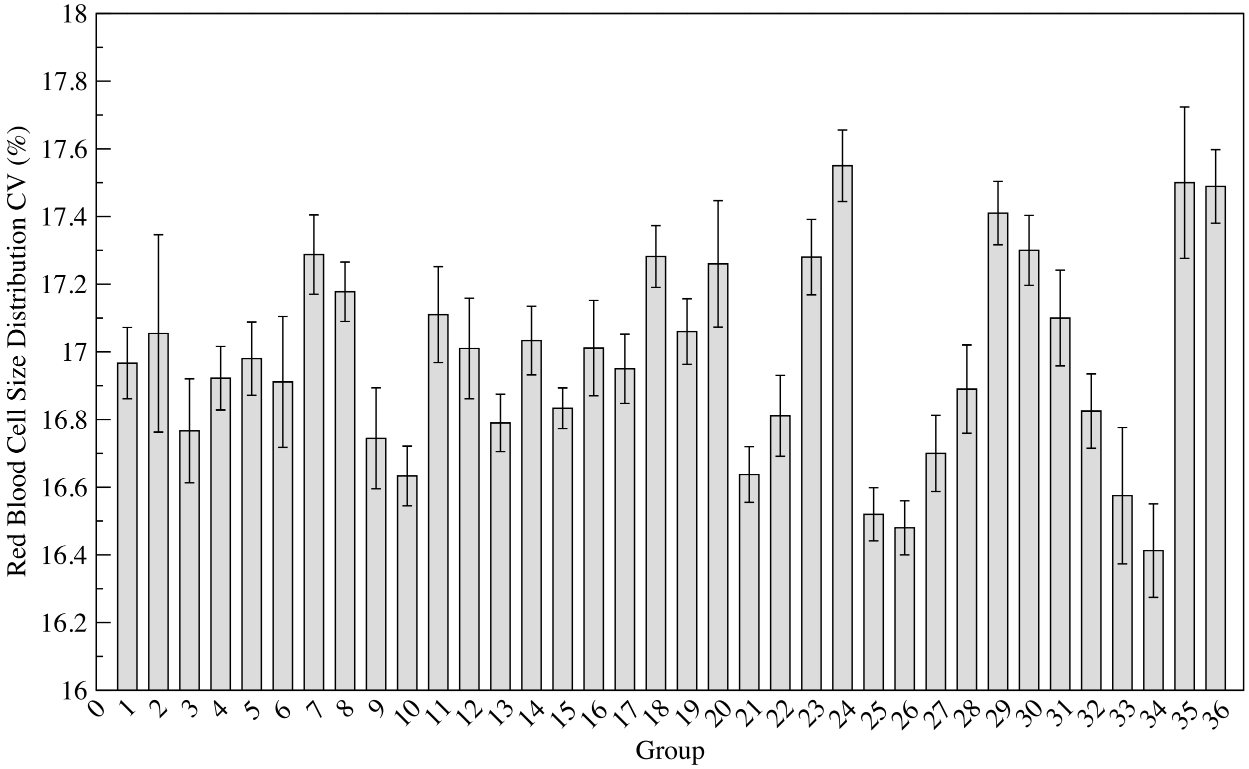


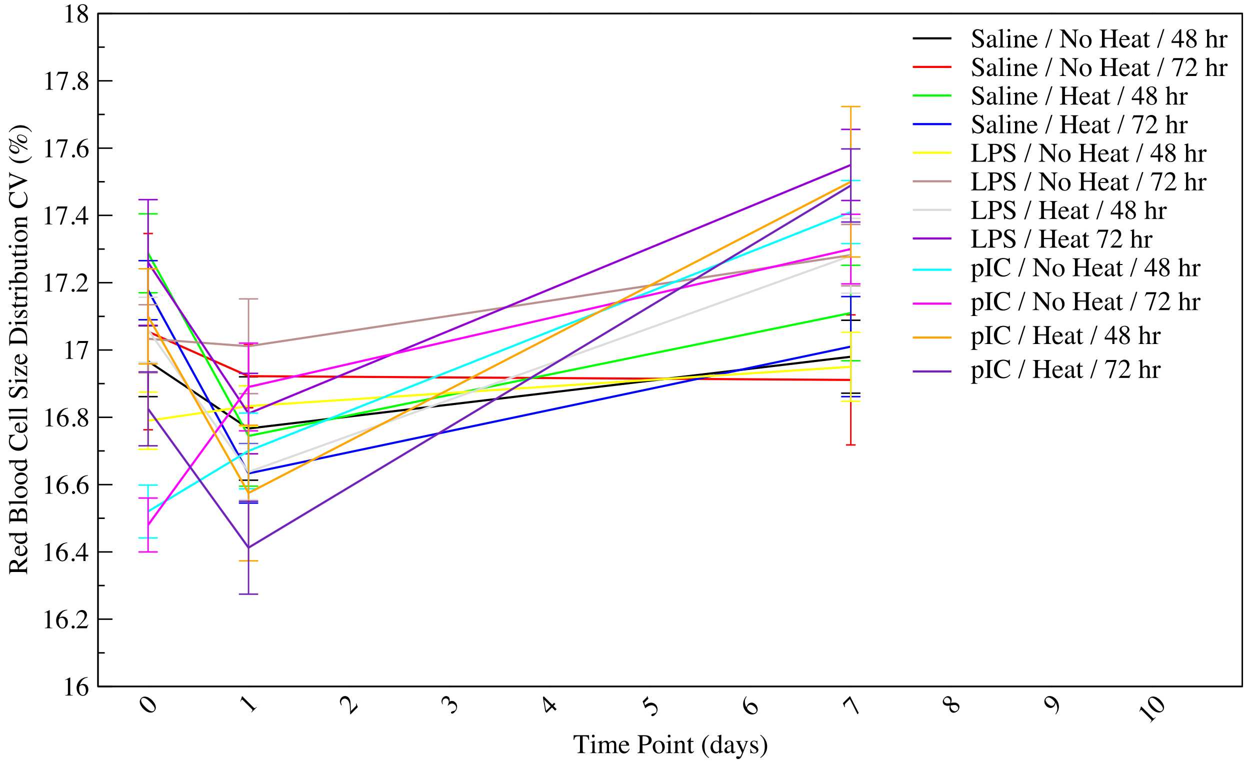

**S. Platelet concentration** is measured in millions of cells per milliliter, and hence is influenced by blood volume and dehydration. Heated animals experience an increase in platelet concentration at $T_{c,max}$, followed by a drop to baseline at 1 day and an increase to nearly $T_{c,max}$ levels at 7 days. Unheated pIC treated animals have decreased platelet concentrations.

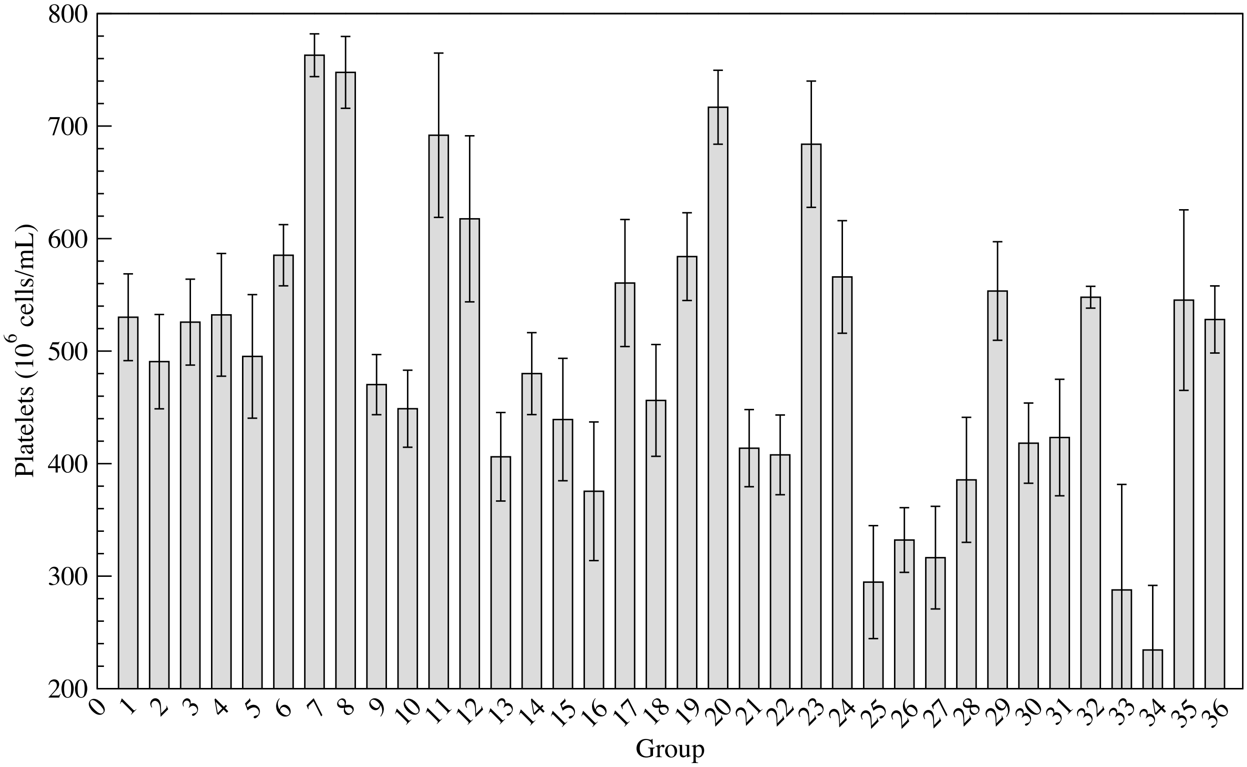


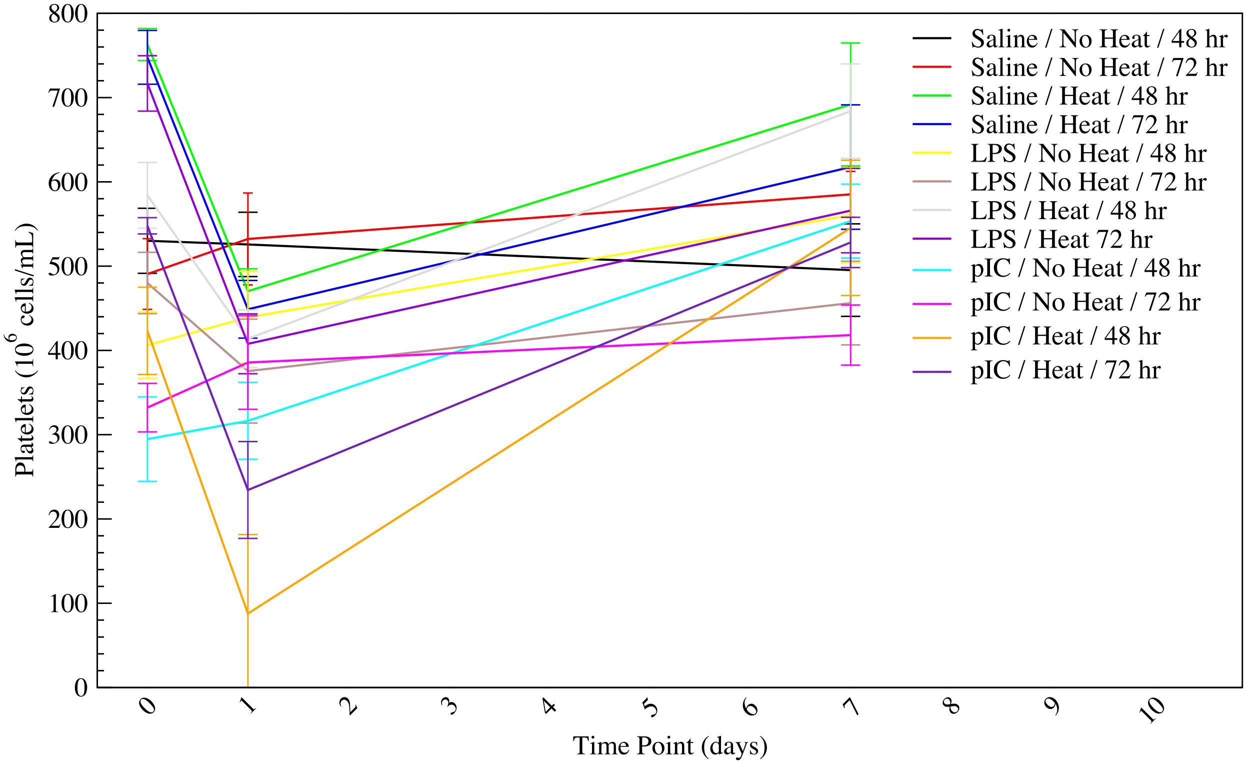

**T. Platelet percentage**, or plateletcrit, is the volume occupied by platelets in the blood. Platelet percentage follows the same trends as platelet concentration.

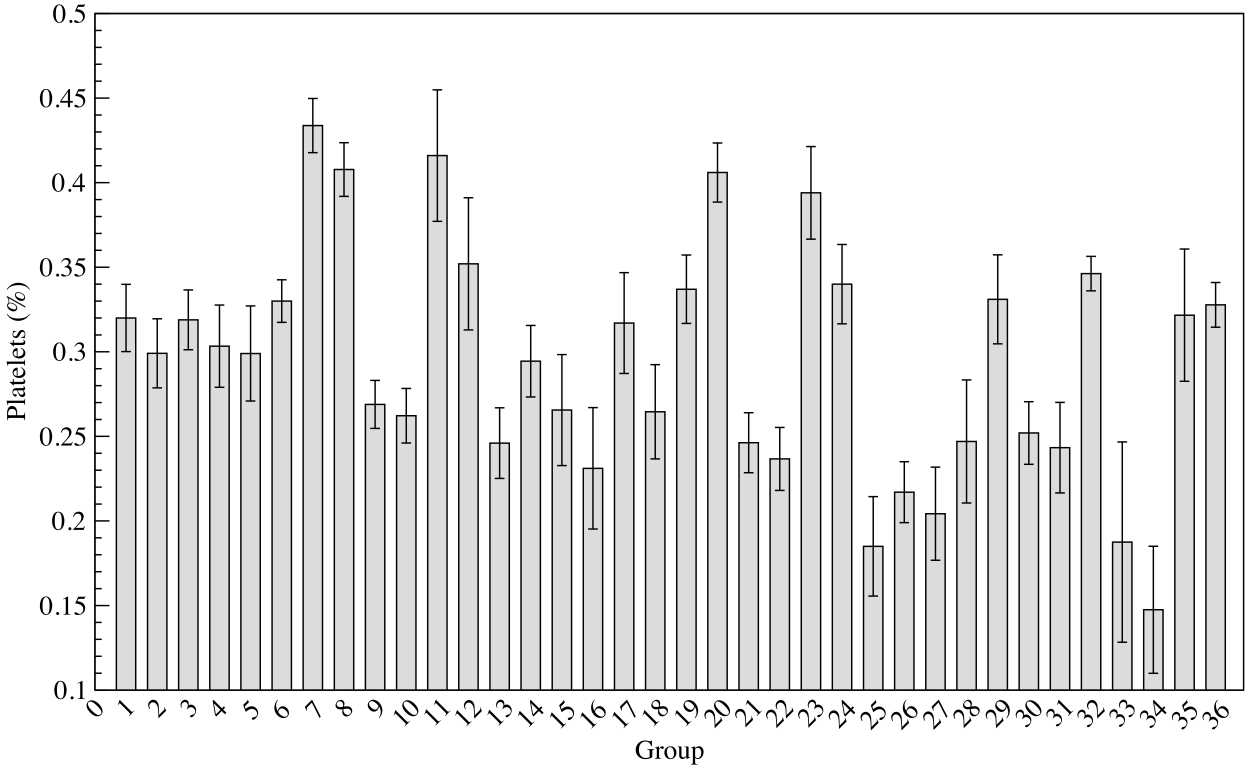


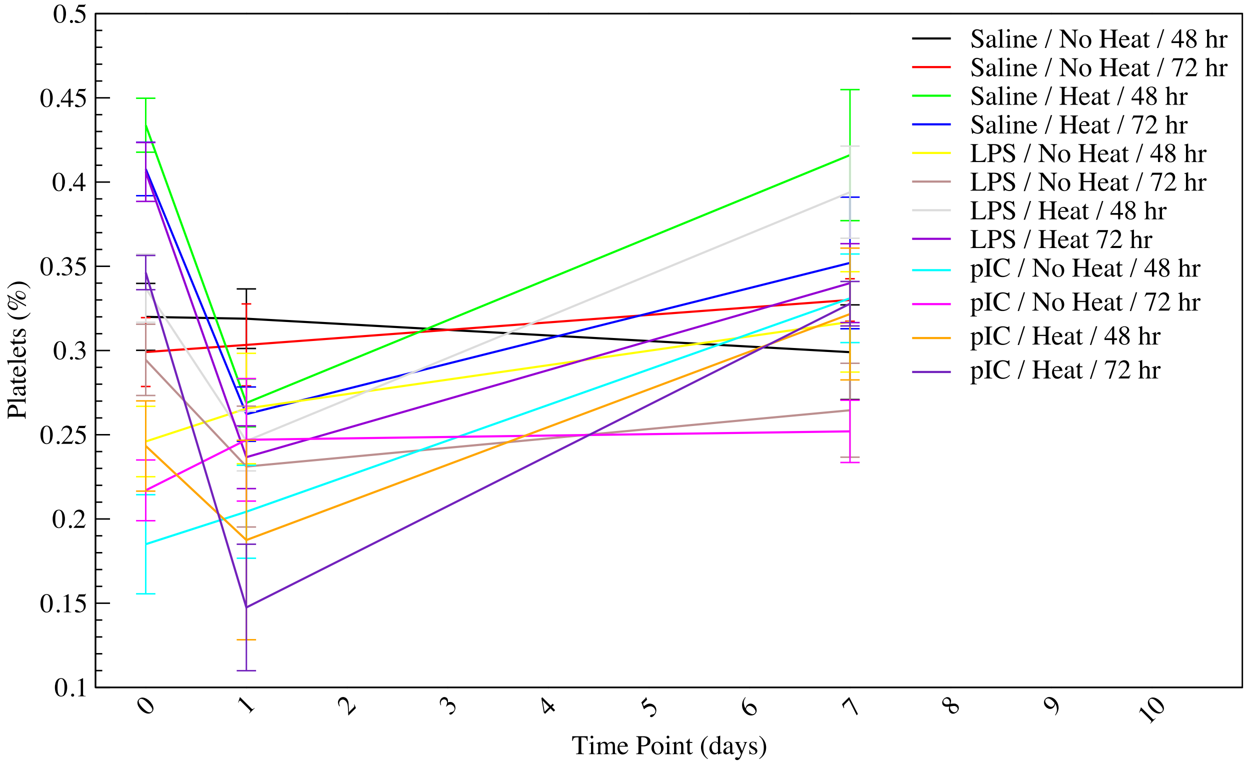

**U. Mean platelet volume** is the average volume of a platelet, measured in femtoliters. Measurements were highly variable, including in saline unheated controls. Heated animals trend toward lower platelet volume at $T_{c,max}$, indicating impaired platelet production, with a recovery to baseline at later time points. pIC treated animals tend to have slightly higher platelet volume across time points, indicating destruction of platelets.

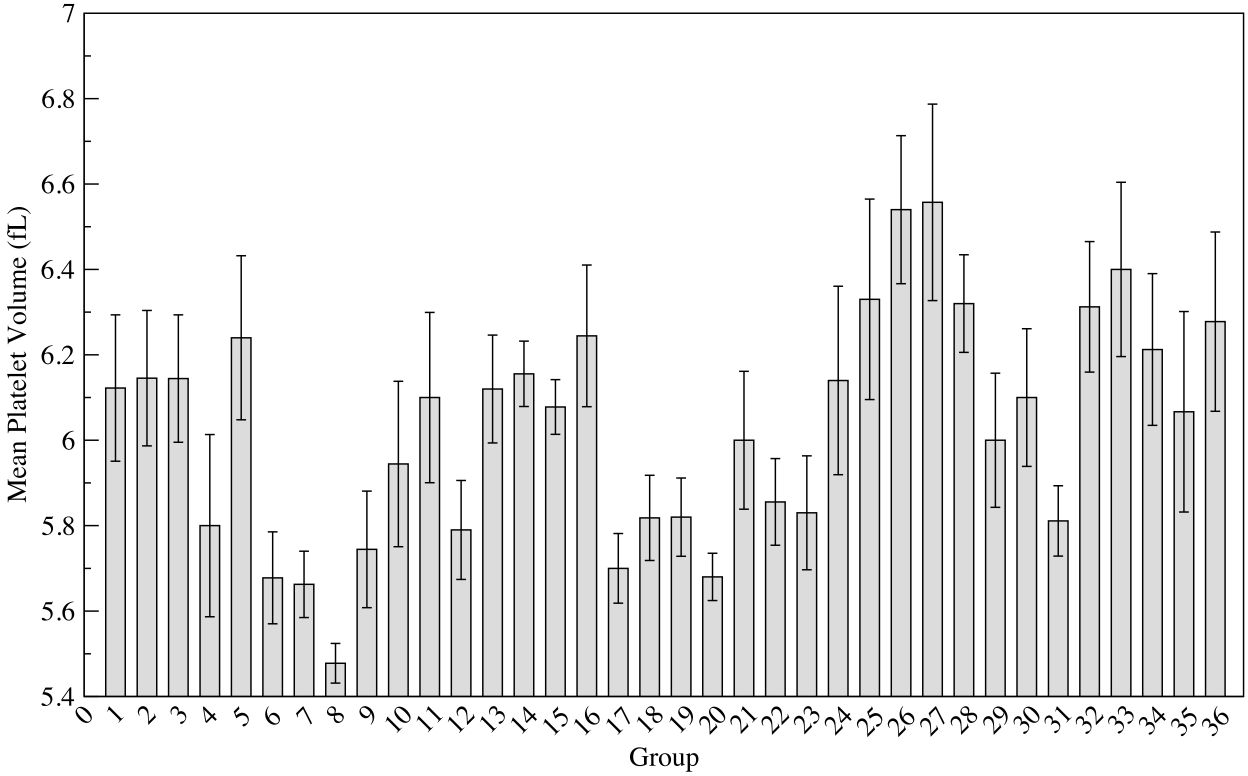


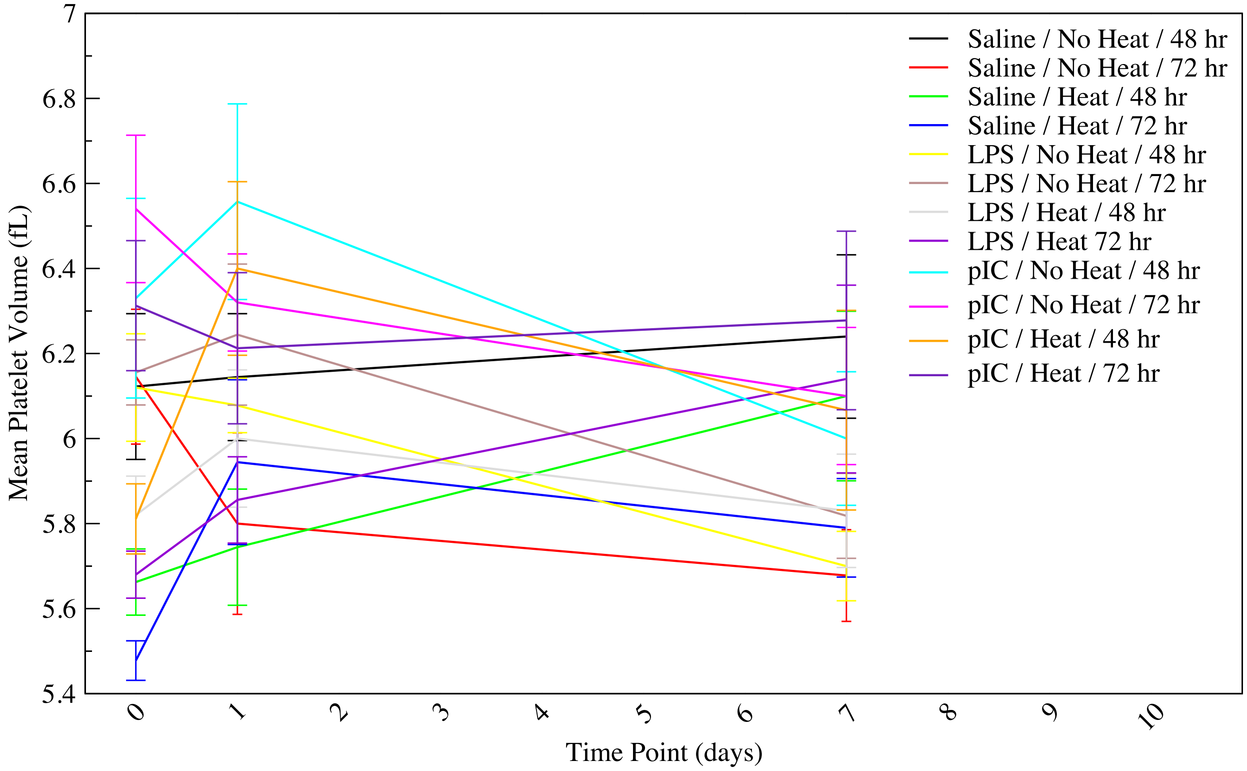

**V. Platelet size distribution coefficient of variation** is a measure of the range of variation of platelet volume. Measurements were highly variable, including in saline unheated controls. Immune challenge tends to elevate the variation in platelet size, while heat challenge tends to decrease the variation. Combined with mean platelet volume, above, shifts in size distribution also shift the mean of the distribution, with heated animals featuring a tighter distribution of smaller platelets and immune challenged animals featuring a broader distribution of slightly larger platelets. The former implies a population of older platelets with no new generation, while the latter implies increased platelet shedding.

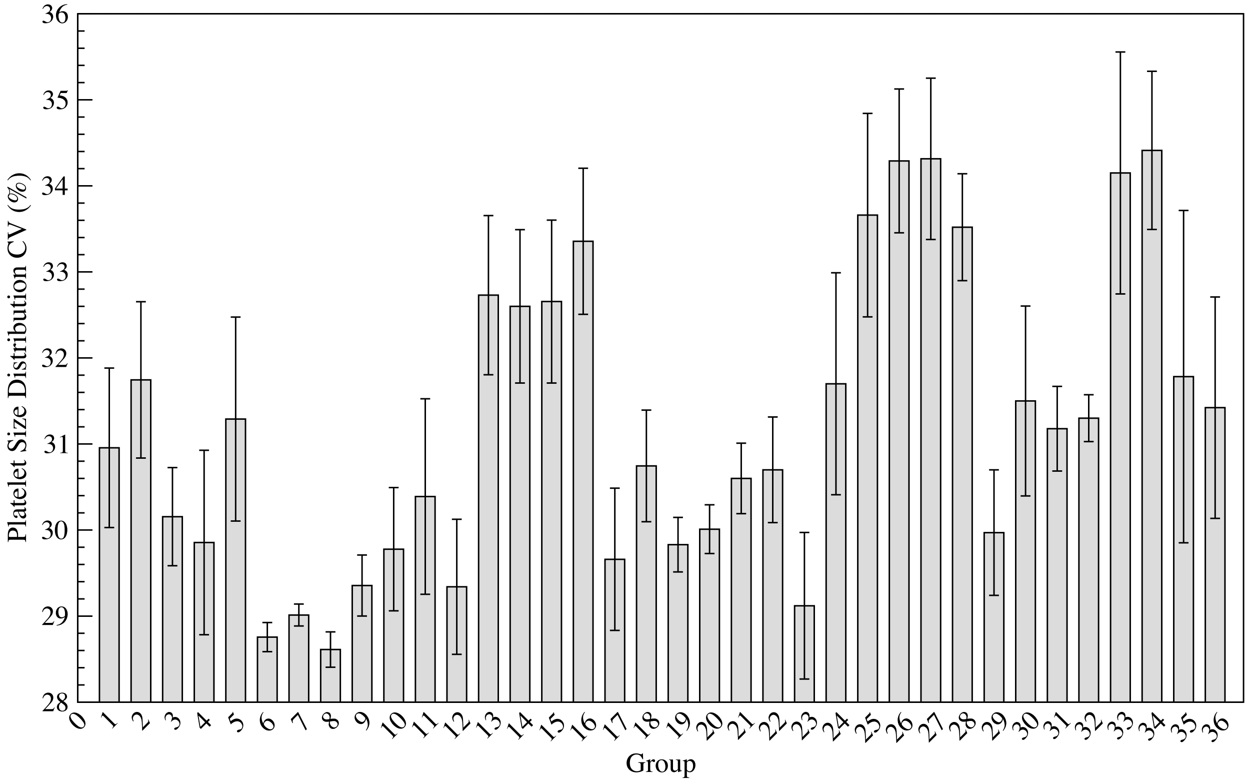


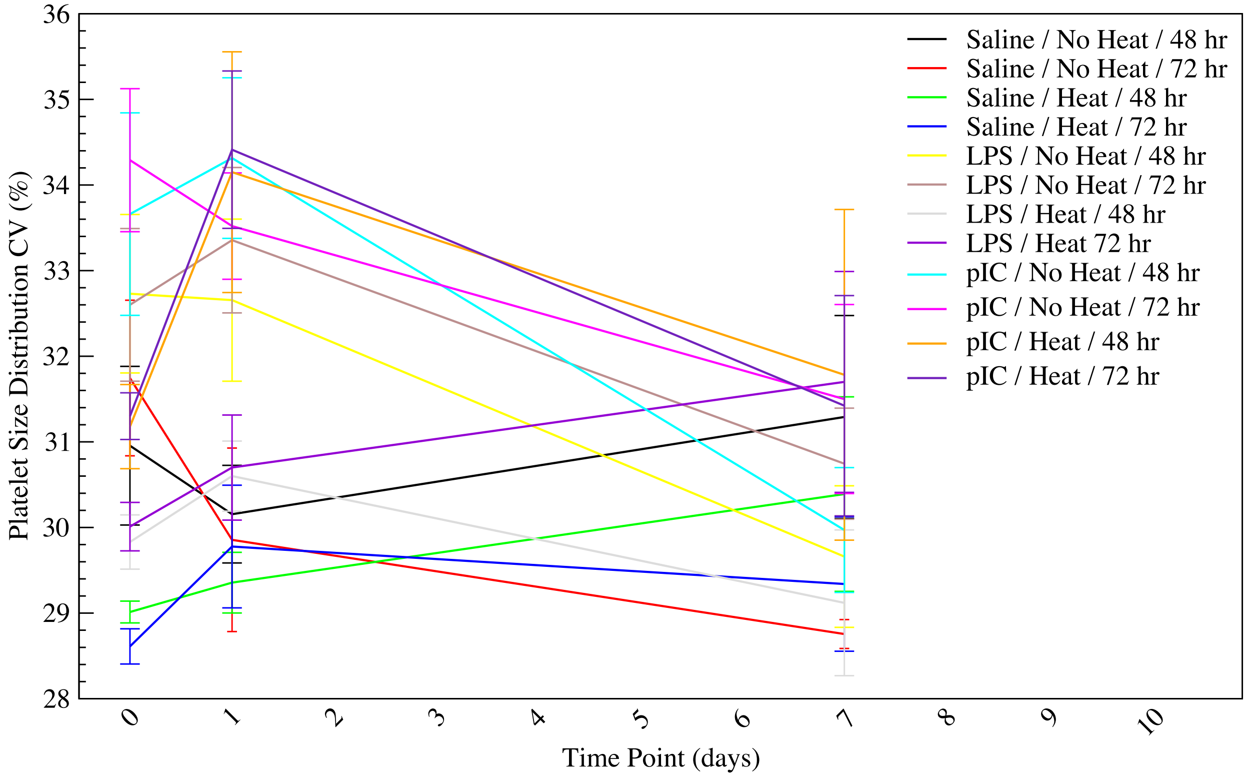

**W. Granzyme B concentration** was measured in the liver and is a marker for apoptosis, hence commonly denoting the level of organ damage. Not all groups were represented in this analysis, but treatment was the largest differentiator, with granzyme B concentratin as saline > LPS > pIC. Heated animals display a slight trend toward higher granzyme B concentration at each time point.

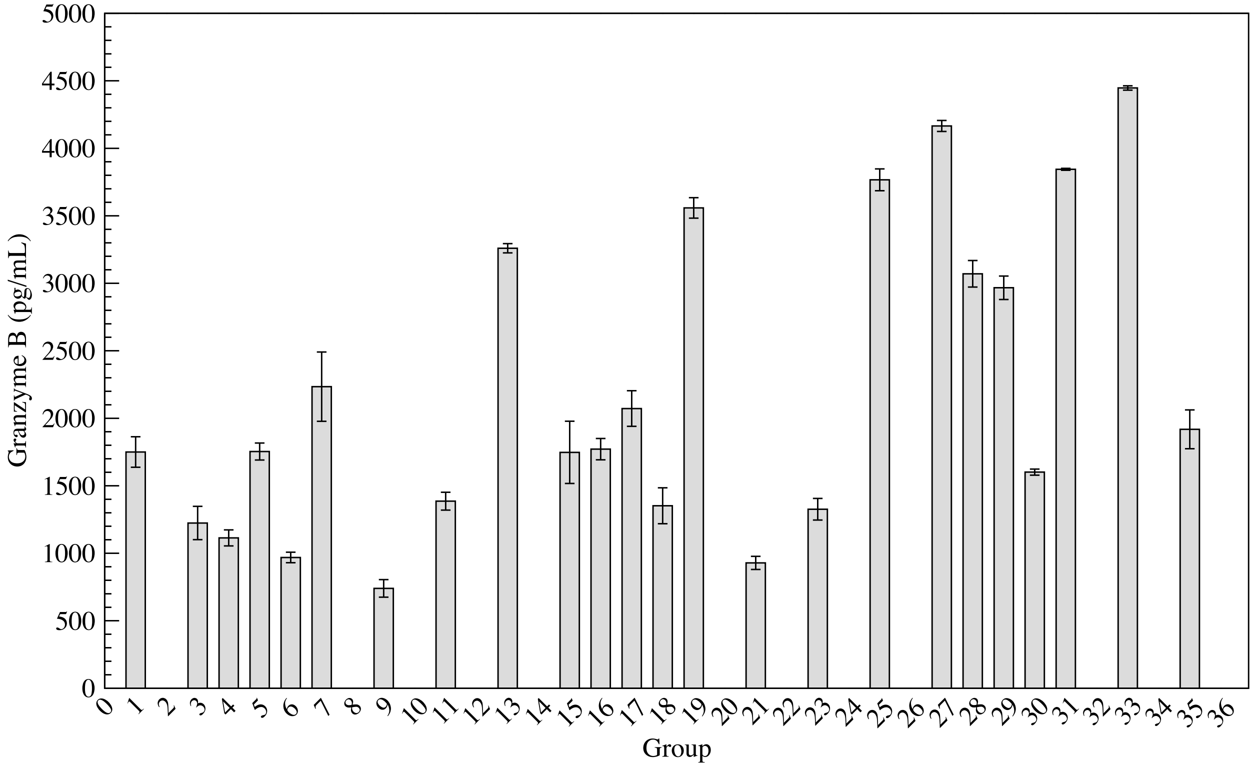


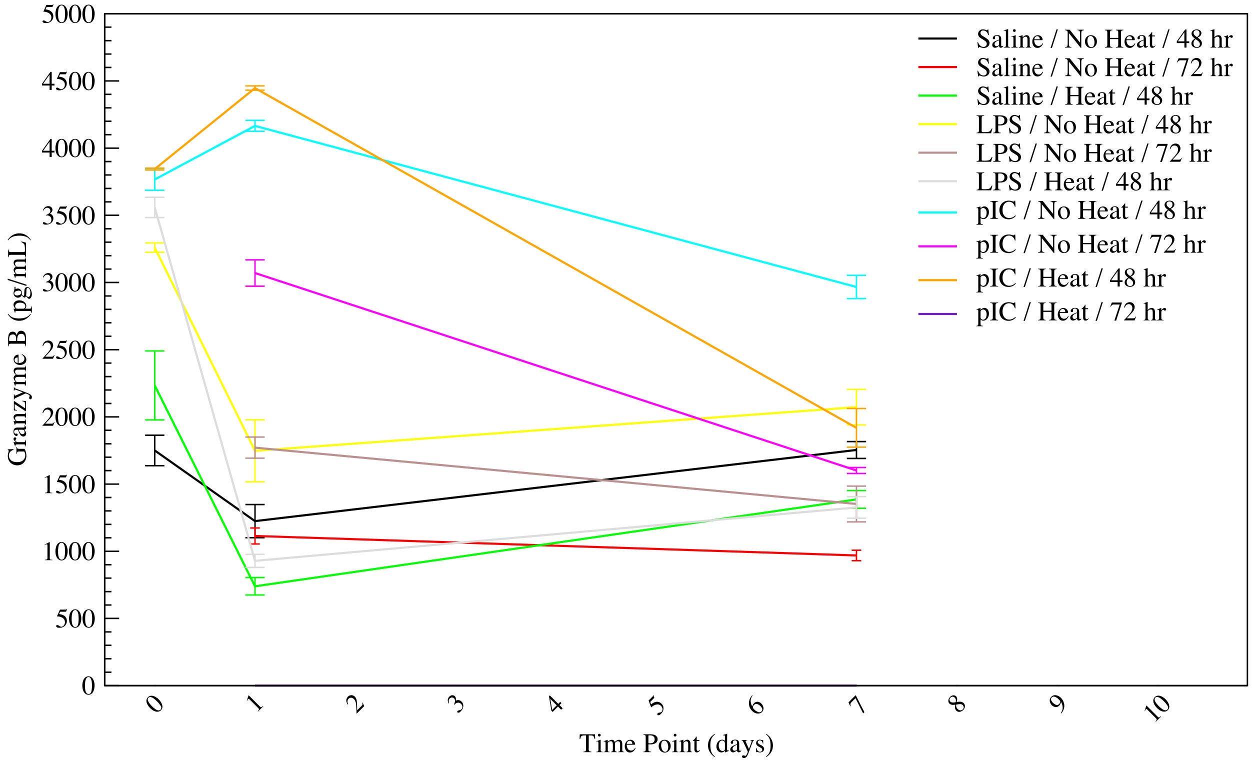

**Y. D-dimer concentration** is the concentration in nanograms per milliliter of a protein degradation product present in the blood following the breakdown of a blood clot (fibrinolysis). This measurement was not performed on LPS treated groups, as LPS is known to interfere with coagulation-related processes. Both heat challenge and pIC treatment increased the evidence of blood clot breakdown, implying an increase in coagulation.

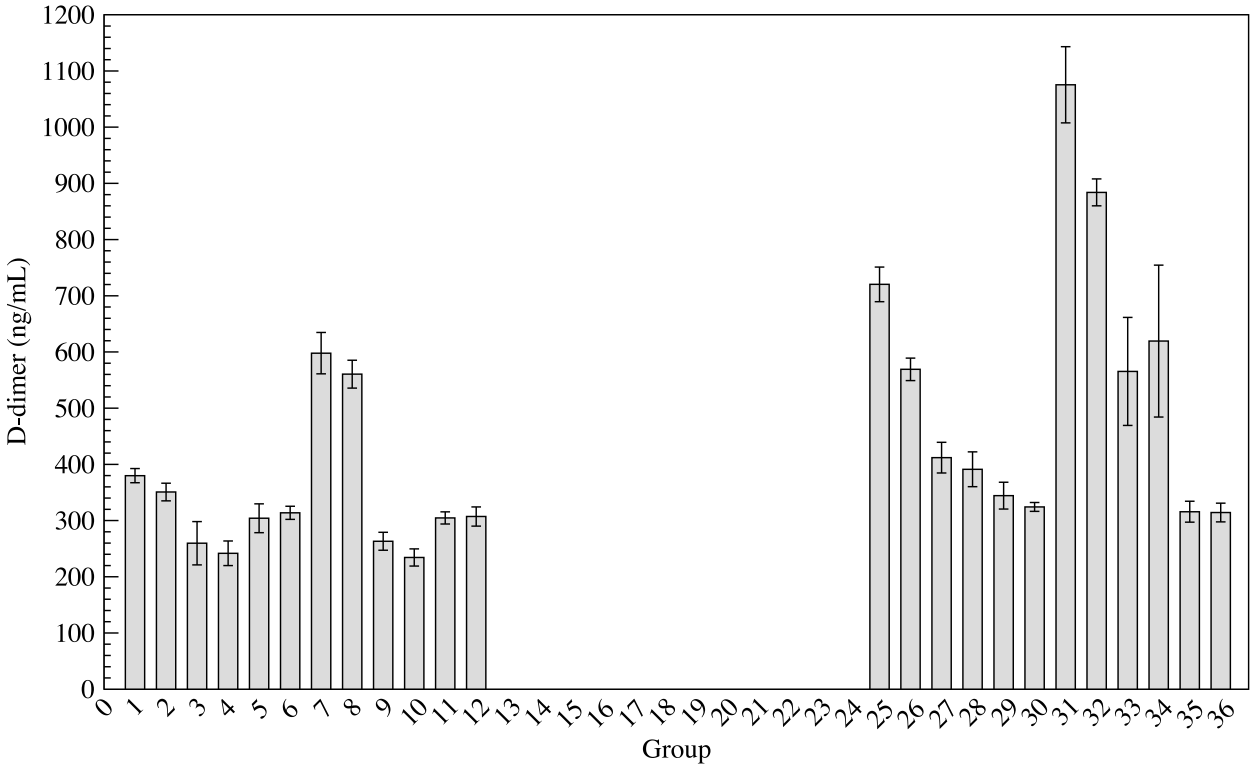


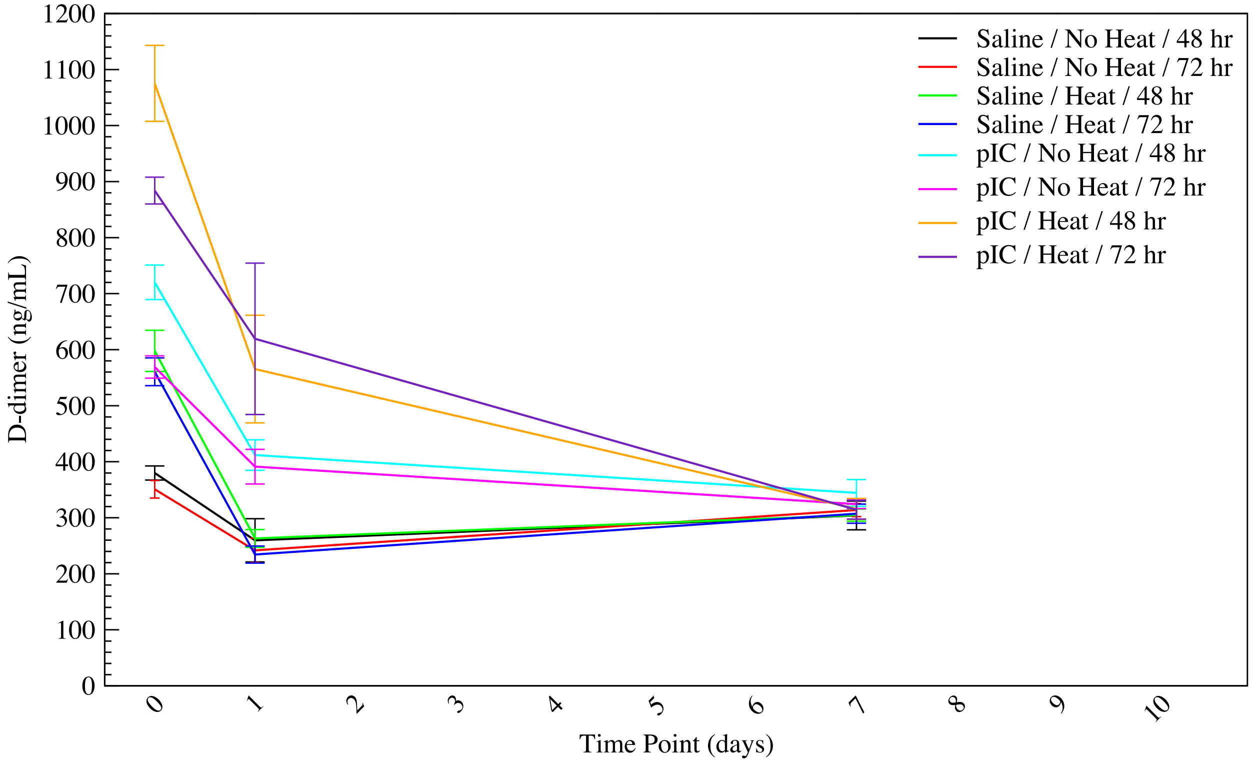

**Z. Thrombin-antithrombin complex concentration** is the concentration in nanograms per milliliter of the complex formed between thrombin and antithrombin in response to high thrombin levels caused by coagulation following a ruptured blood vessel. Because antithrombin is present at much higher levels than thrombin, the concentration of the complex is mostly equivalent to the concentration of thrombin. This measurement was not performed on LPS treated groups, as LPS is known to interfere with coagulation-related processes. pIC treatment elevated levels of thrombin-antithrombin complex at the 1 day time point.

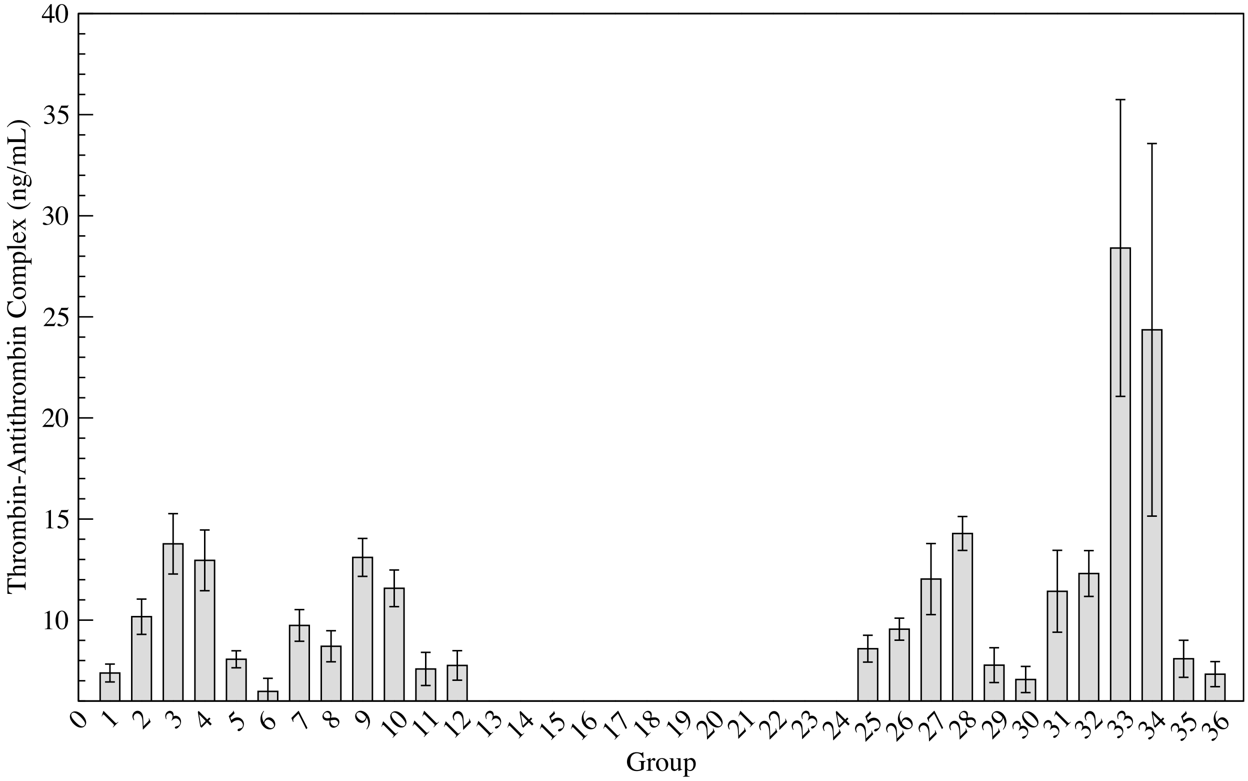


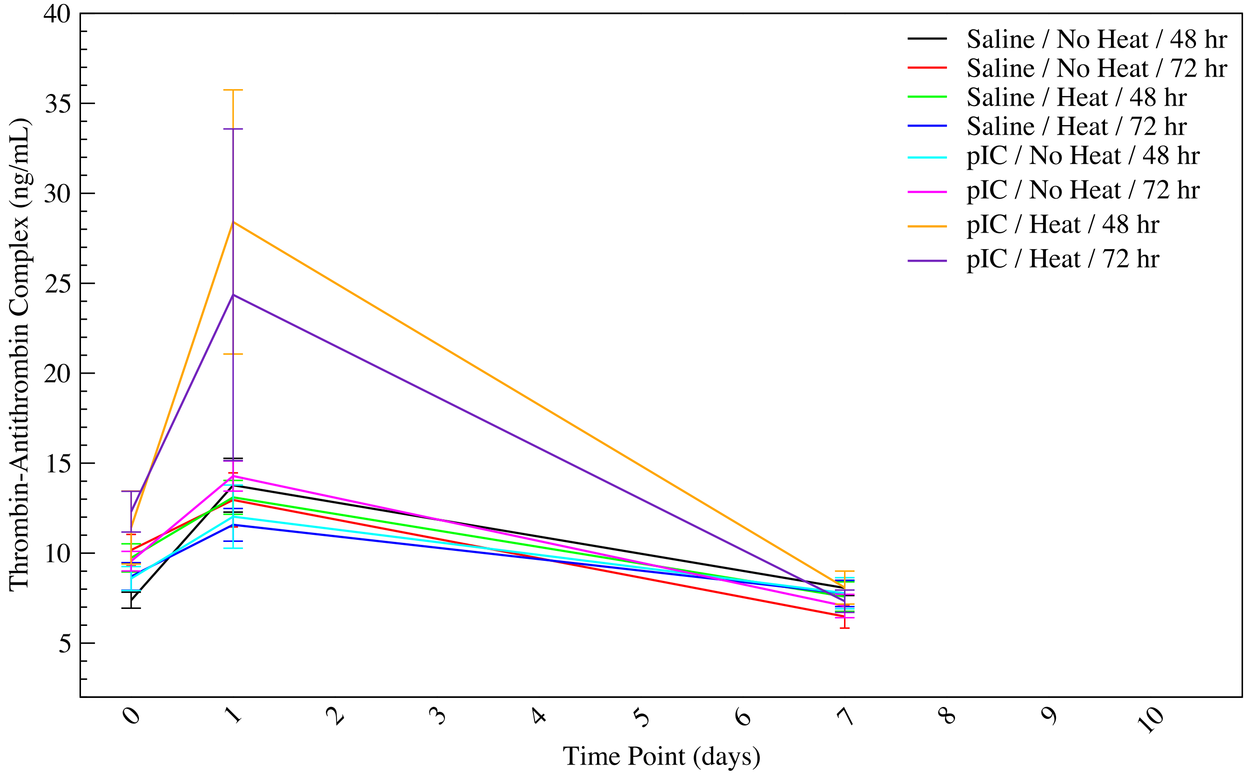

**AA. Antithrombin III concentration** is the concentration in micrograms per milliliter of a protein inhibitor of the coagulation system. Antithrombin is generally present at high levels in the blood stream. This measurement was not performed on LPS treated groups, as LPS is known to interfere with coagulation-related processes. Both heat challenge and pIC treatment decrease levels of this anticoagulant protein.

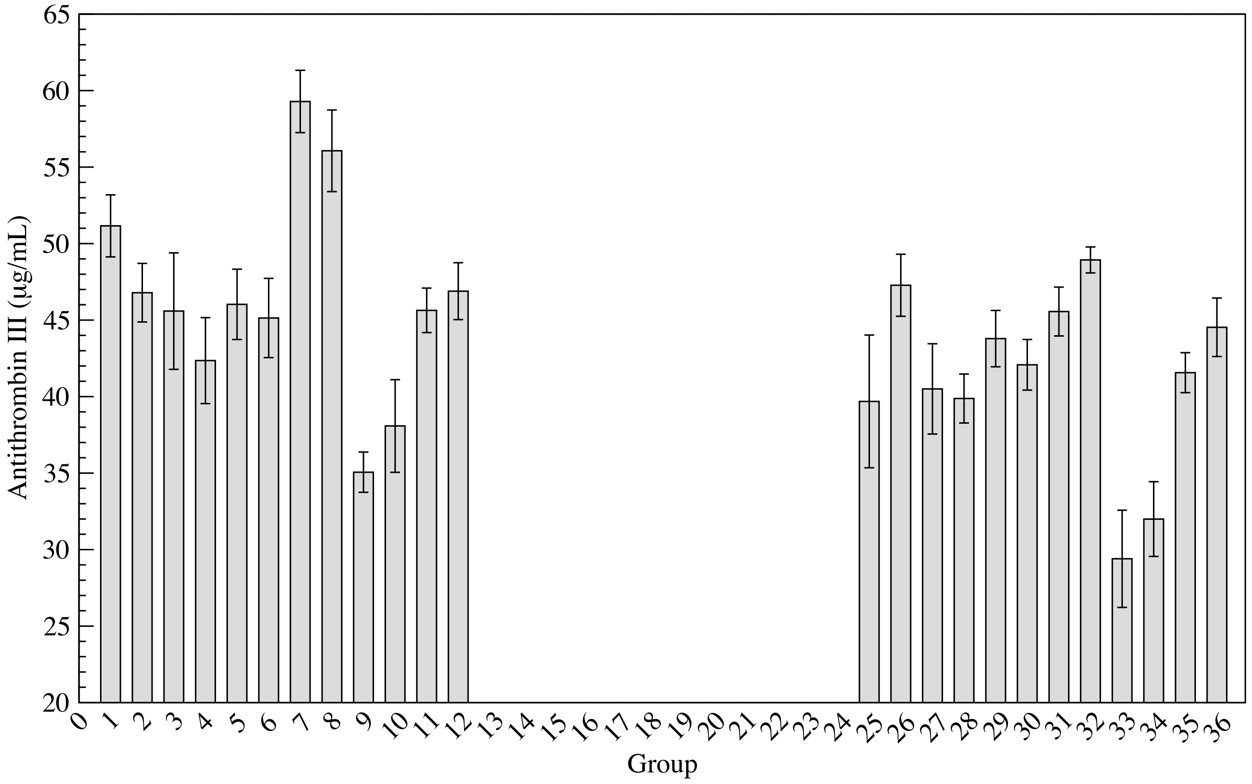


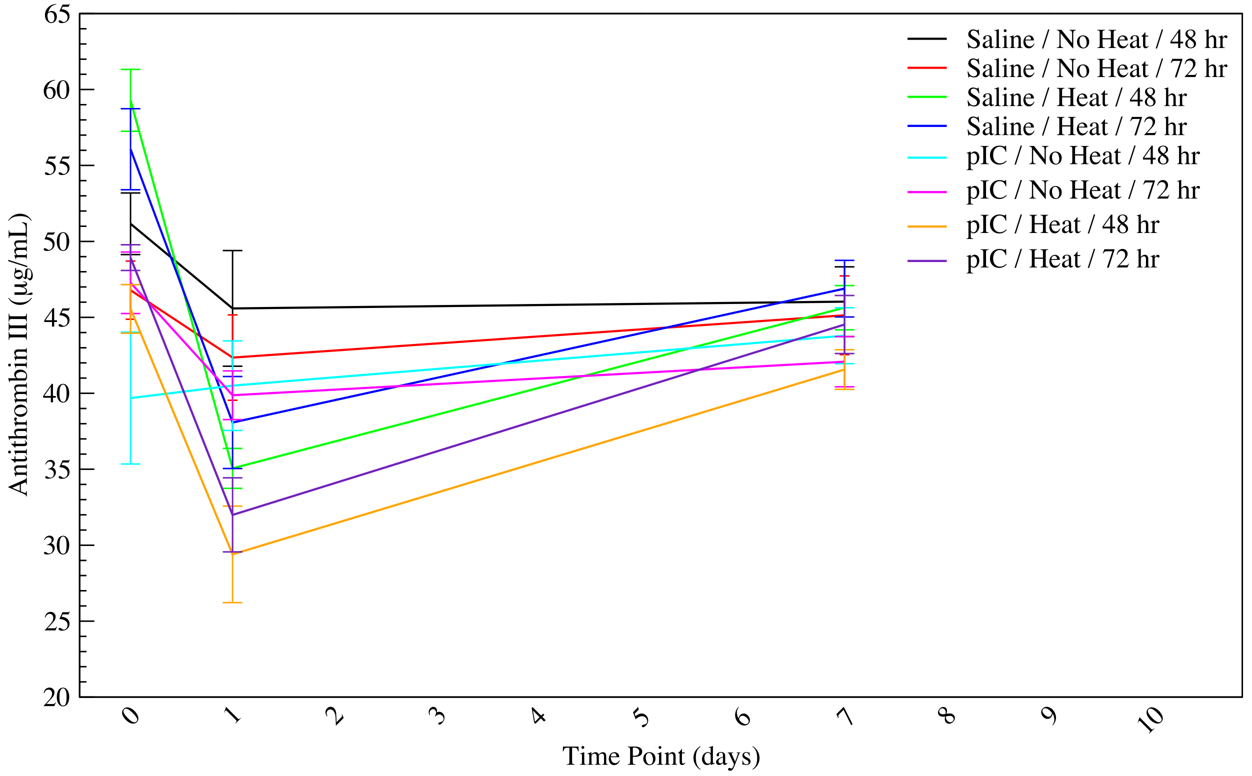

**AB. Thrombomodulin concentration** is the concentration in nanograms per milliliter of soluble thrombomodulin, a marker of endothelial damage and prothrombotic state. This measurement was not performed on LPS treated groups, as LPS is known to interfere with coagulation-related processes. Both heat challenge and pIC treatment increase levels of this coagulation marker.

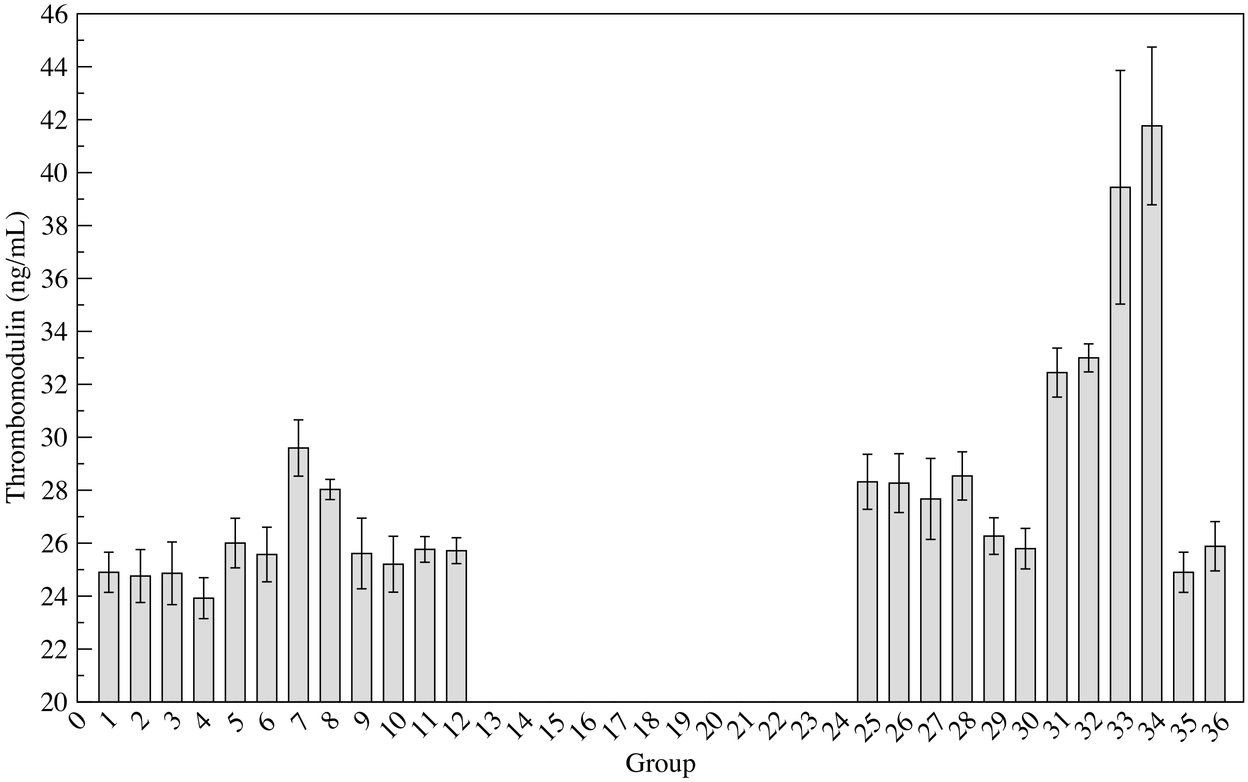


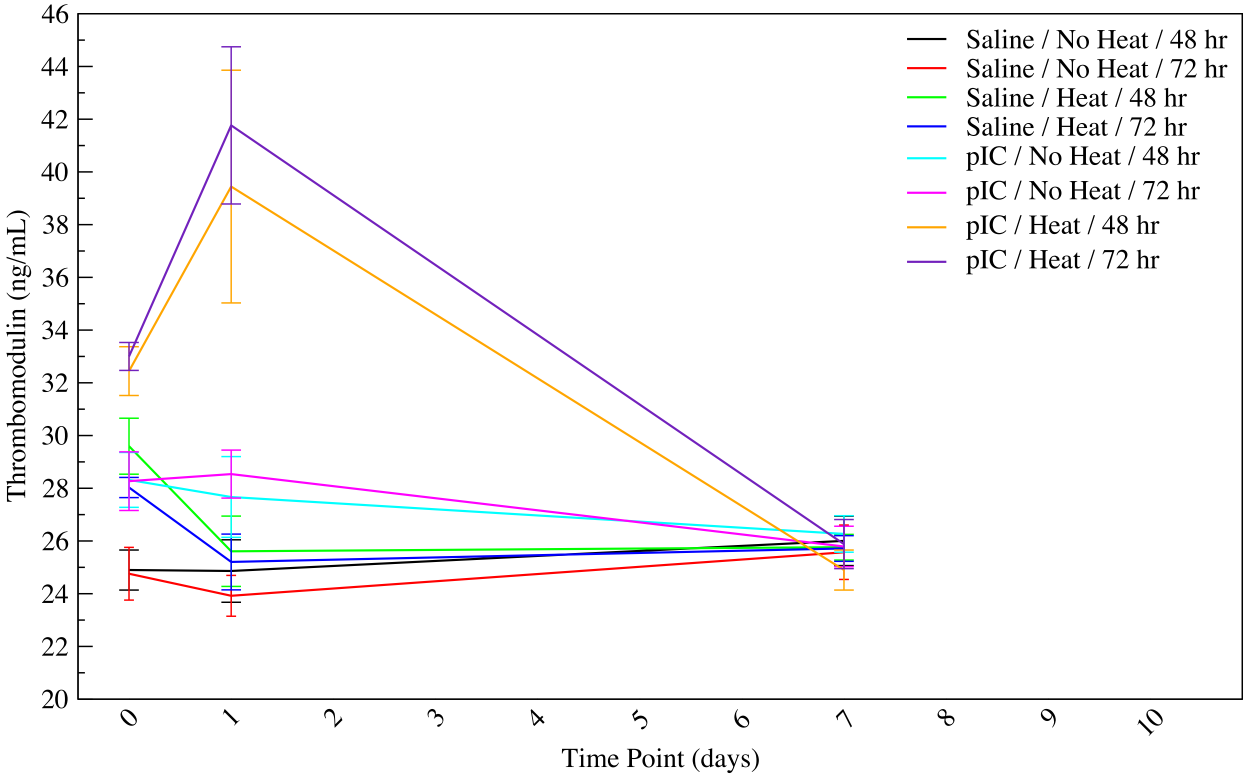

**AC. Tissue factor concentration** is the concentration in picograms per milliliter of a protein that participates in the initiation of the coagulation process. This measurement was not performed on LPS treated groups, as LPS is known to interfere with coagulation-related processes, and was only performed at $T_{c,max}$ time points due to limitations in the amount of blood required for all assays. Heat challenge increases levels of tissue factor, while pIC treatment decreases levels.

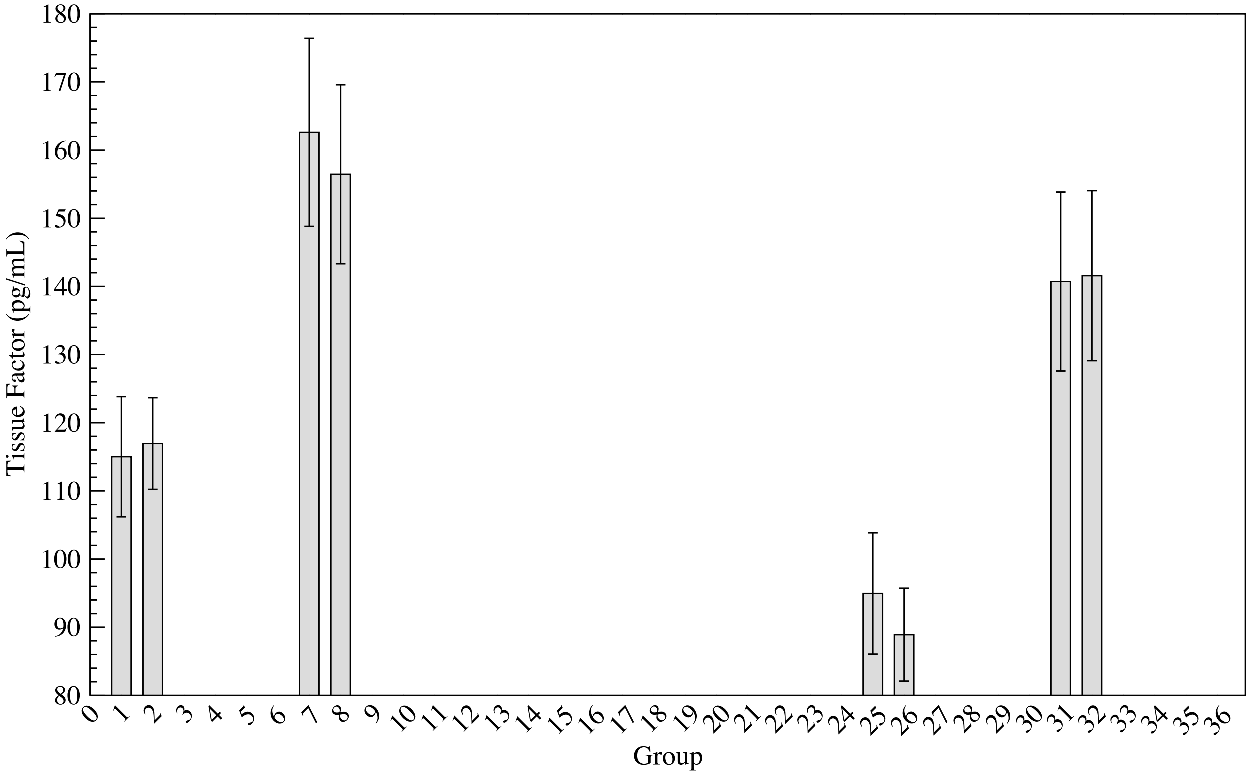


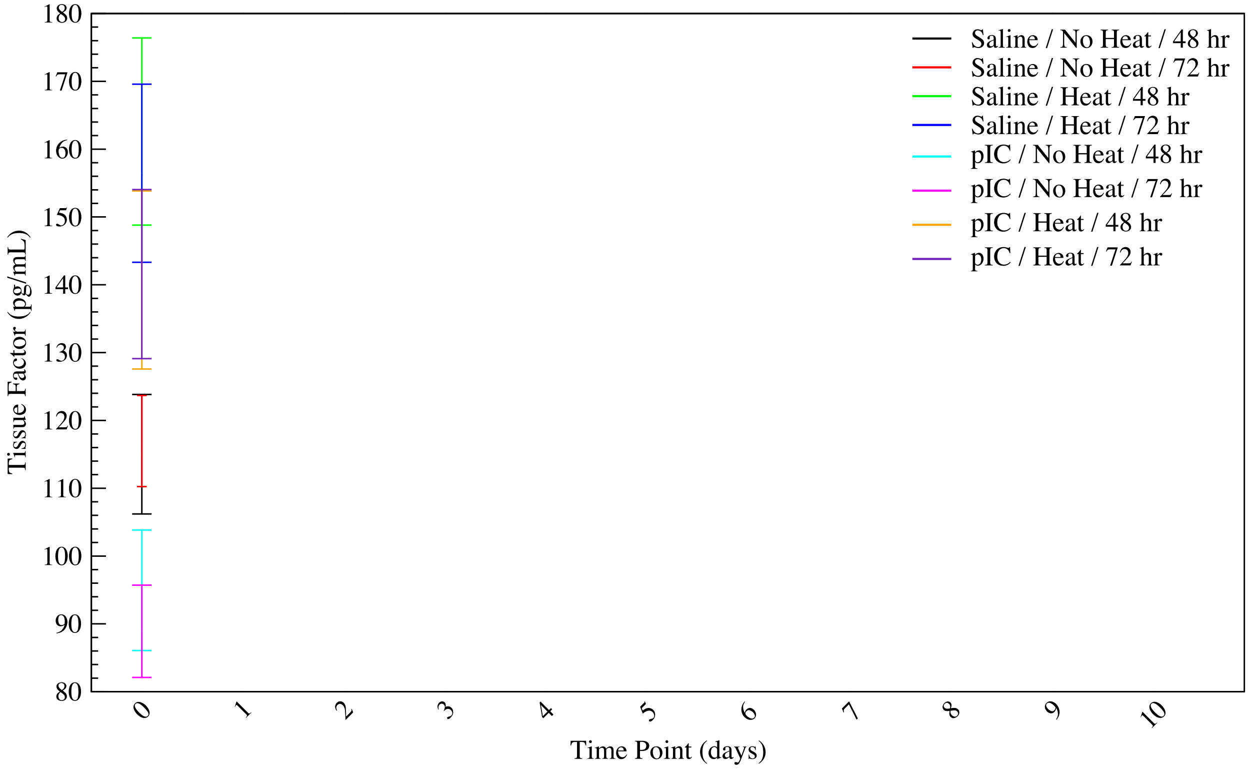

**Table S3. Enrichment of gene sets in severe heat stroke.** Hallmark gene sets with normalized enrichment scores (NES) and FDR q-values, as differentially present in poly I:C-treated, heat-challenged mice versus controls (saline-treated, unheated). Only gene sets with FDRq<0.25 are shown. Positive NES denotes up-regulation of transcription in the heat stroked mice, while negative NES denotes down-regulation of transcription.

| Gene Set | NES | FDR q-value |
|---|---|---|
| HALLMARK_TNFA_SIGNALING_VIA_NFKB | 6.3025413 | 0 |
| HALLMARK_INTERFERON_GAMMA_RESPONSE | 4.7531977 | 0 |
| HALLMARK_MYC_TARGETS_V1 | 3.9991922 | 0 |
| HALLMARK_HYPOXIA | 3.8294756 | 0 |
| HALLMARK_INTERFERON_ALPHA_RESPONSE | 3.721243 | 0 |
| HALLMARK_KRAS_SIGNALING_UP | 3.5039012 | 0 |
| HALLMARK_APOPTOSIS | 3.5034516 | 0 |
| HALLMARK_IL6_JAK_STAT3_SIGNALING | 3.414408 | 0 |
| HALLMARK_INFLAMMATORY_RESPONSE | 3.3083599 | 0 |
| HALLMARK_P53_PATHWAY | 3.1333308 | 0 |
| HALLMARK_E2F_TARGETS | 2.7685106 | 0 |
| HALLMARK_IL2_STAT5_SIGNALING | 2.7541401 | 0 |
| HALLMARK_ALLOGRAFT_REJECTION | 2.7365997 | 0 |
| HALLMARK_MYC_TARGETS_V2 | 2.6895933 | 0 |
| HALLMARK_UNFOLDED_PROTEIN_RESPONSE | 2.6467988 | 0 |
| HALLMARK_EPITHELIAL_MESENCHYMAL_TRANSITION | 2.6284373 | 0 |
| HALLMARK_G2M_CHECKPOINT | 2.5481465 | 7.93E-05 |
| HALLMARK_COMPLEMENT | 2.482341 | 3.28E-04 |
| HALLMARK_MTORC1_SIGNALING | 2.464753 | 3.88E-04 |
| HALLMARK_ESTROGEN_RESPONSE_EARLY | 2.2387335 | 0.0017964748 |
| HALLMARK_UV_RESPONSE_UP | 2.0070226 | 0.008110026 |
| HALLMARK_ESTROGEN_RESPONSE_LATE | 1.8552 | 0.016249517 |
| HALLMARK_MITOTIC_SPINDLE | 1.8096329 | 0.02029716 |
| HALLMARK_ANDROGEN_RESPONSE | 1.4919591 | 0.09301928 |
| HALLMARK_PI3K_AKT_MTOR_SIGNALING | 1.4709035 | 0.096499436 |
| HALLMARK_COAGULATION | 1.4387547 | 0.10824509 |
| HALLMARK_DNA_REPAIR | 1.3416716 | 0.15439369 |
| HALLMARK_PROTEIN_SECRETION | -1.2667292 | 0.22302693 |
| HALLMARK_GLYCOLYSIS | -1.3773451 | 0.16253032 |
| HALLMARK_CHOLESTEROL_HOMEOSTASIS | -1.4660783 | 0.13014264 |

| | | |
|---|---|---|
| HALLMARK_KRAS_SIGNALING_DN | -1.6269449 | 0.07398985 |
| HALLMARK_PEROXISOME | -1.6502489 | 0.07926317 |
| HALLMARK_BILE_ACID_METABOLISM | -1.8810419 | 0.029077718 |
| HALLMARK_ADIPOGENESIS | -2.268545 | 0.004759706 |
| HALLMARK_FATTY_ACID_METABOLISM | -2.7723553 | 0 |
| HALLMARK_OXIDATIVE_PHOSPHORYLATION | -4.2311616 | 0 |

**Table S4. Organ histopathology scores in heat-stroked mice.** Samples of liver, kidney, spleen, lung, and duodenum from the same 48 mice used for RNAseq analysis of kidney, with an additional 0-2 mice per group depending on availability. Severity is graded as: 0 = no significant change, 1 = minimal, 2 = mild, 3 = moderate, and 4 = severe, with the number of mice exhibiting each abnormality given in the first of the three columns for each group, followed by the mean score for the whole group and the mean score for the subset of affected mice (if different from the total number of mice in the group). Group 1: pIC/noheat/1d, Group 2: pIC/heat/1d, Group 3: pIC/noheat/Tcmax, Group 4: pIC/heat/Tcmax, Group 5: saline/noheat/1d, Group 6: saline/heat/1d, Group 7: saline/noheat/Tcmax, Group 8: saline/heat/Tcmax.

| | poly I:C, no heat, 1 day | | | poly I:C, heat, 1 day | | | poly I:C, no heat, $T_{c,max}$ | | | poly I:C, heat, $T_{c,max}$ | | |
|---|---|---|---|---|---|---|---|---|---|---|---|---|
| **Number** | **# Abnormal** | **Mean Score** | **Mean-Affected** | **# Abnormal** | **Mean Score** | **Mean-Affected** | **# Abnormal** | **Mean Score** | **Mean-Affected** | **# Abnormal** | **Mean Score** | **Mean-Affected** |
| | **n=** | **8** | | **n=** | **6** | | **n=** | **8** | | **n=** | **8** | |
| **1- Liver** | | | | | | | | | | | | |
| Extramedullary hematopoiesis | 0 | 0.0 | | 0 | 0.0 | | 2 | 0.3 | 1.0 | 0 | 0.0 | |
| Vacuolation, hepatocellular | 8 | 2.8 | 2.8 | 2 | 0.8 | 2.5 | 7 | 1.5 | 1.7 | 0 | 0.0 | |
| Inflammation, intrahepatic | 1 | 0.1 | 1.0 | 0 | 0.0 | | 3 | 0.6 | 1.7 | 6 | 0.9 | 1.2 |
| Inflammation, mesentery/ capsule | 1 | 0.3 | 2.0 | 0 | 0.0 | | 0 | 0.0 | | 0 | 0.0 | |
| Necrosis | 0 | 0.0 | | 0 | 0.0 | | 0 | 0.0 | | 0 | 0.0 | |
| Degeneration, hepatocellular | 0 | 0.0 | | 3 | 1.4 | 2.8 | 3 | 0.9 | 2.5 | 6 | 0.9 | 1.2 |
| Mineralization | 0 | 0.0 | | 0 | 0.0 | | 0 | 0.0 | | 0 | 0.0 | |
| Neoplasia | 0 | 0.0 | | 0 | 0.0 | | 0 | 0.0 | | 0 | 0.0 | |
| Hyperplasia, oval cell | 0 | 0.0 | | 1 | 0.2 | 1.0 | 1 | 0.1 | 1.0 | 1 | 0.1 | 1.0 |
| Vasculitis | 0 | 0.0 | | 0 | 0.0 | | 0 | 0.0 | | 0 | 0.0 | |
| Atrophy | 0 | 0.0 | | 0 | 0.0 | | 0 | 0.0 | | 0 | 0.0 | |
| Lymphoid Depletion | 0 | 0.0 | | 0 | 0.0 | | 0 | 0.0 | | 0 | 0.0 | |
| Artifact | 4 | 1.0 | 2.0 | 0 | 0.0 | | 2 | 0.4 | 1.5 | 1 | 0.3 | 2.0 |
| Sum - Scores: | 10 | 3.1 | 5.8 | 6 | 2.4 | 6.3 | 16 | 3.4 | 7.9 | 13 | 1.9 | 3.3 |
| No Significant Findings: | 0 | | | 1 | | | 0 | | | 1 | | |
| | **n=** | **8** | | **n=** | **6** | | **n=** | **8** | | **n=** | **8** | |
| **2- Kidney** | | | | | | | | | | | | |
| Hemorrhage | 0 | 0.0 | | 0 | 0.0 | | 0 | 0.0 | | 0 | 0.0 | |
| Edema | 0 | 0.0 | | 0 | 0.0 | | 0 | 0.0 | | 0 | 0.0 | |
| Inflammation | 0 | 0.0 | | 0 | 0.0 | | 0 | 0.0 | | 0 | 0.0 | |
| Degeneration | 0 | 0.0 | | 0 | 0.0 | | 0 | 0.0 | | 0 | 0.0 | |
| Necrosis | 0 | 0.0 | | 0 | 0.0 | | 0 | 0.0 | | 0 | 0.0 | |
| Autolysis | 0 | 0.0 | | 0 | 0.0 | | 0 | 0.0 | | 0 | 0.0 | |
| Mineralization | 0 | 0.0 | | 0 | 0.0 | | 0 | 0.0 | | 0 | 0.0 | |
| Neoplasia | 0 | 0.0 | | 0 | 0.0 | | 0 | 0.0 | | 0 | 0.0 | |

| | | | | | | | | | | | | |
|---|---|---|---|---|---|---|---|---|---|---|---|---|
| Hyperplasia | 0 | 0.0 | | 0 | 0.0 | | 0 | 0.0 | | 0 | 0.0 | |
| Vasculitis | 0 | 0.0 | | 0 | 0.0 | | 0 | 0.0 | | 0 | 0.0 | |
| Atrophy | 0 | 0.0 | | 0 | 0.0 | | 0 | 0.0 | | 0 | 0.0 | |
| Lymphoid Depletion | 0 | 0.0 | | 0 | 0.0 | | 0 | 0.0 | | 0 | 0.0 | |
| Artifact | 3 | 0.6 | 1.7 | 2 | 0.8 | 2.5 | 3 | 0.6 | 1.7 | 4 | 1.0 | 2.0 |
| Sum - Scores: | 0 | 0.0 | 0.0 | 0 | 0.0 | 0.0 | 0 | 0.0 | 0.0 | 0 | 0.0 | 0.0 |
| No Significant Findings: | 8 | | | 6 | | | 8 | | | 8 | | |
| | **n=** | **8** | | **n=** | **6** | | **n=** | **8** | | **n=** | **8** | |
| **3- Spleen** | | | | | | | | | | | | |
| Hemorrhage | 0 | 0.0 | | 0 | 0.0 | | 0 | 0.0 | | 0 | 0.0 | |
| Increased neutrophils, red pulp | 0 | 0.0 | | 0 | 0.0 | | 1 | 0.1 | 1.0 | 0 | 0.0 | |
| Inflammation, mesenteric | 4 | 0.8 | 1.5 | 1 | 0.2 | 1.0 | 4 | 0.8 | 1.5 | 0 | 0.0 | |
| Degeneration | 0 | 0.0 | | 0 | 0.0 | | 0 | 0.0 | | 0 | 0.0 | |
| Necrosis | 0 | 0.0 | | 0 | 0.0 | | 0 | 0.0 | | 0 | 0.0 | |
| Autolysis | 0 | 0.0 | | 0 | 0.0 | | 0 | 0.0 | | 0 | 0.0 | |
| Mineralization | 0 | 0.0 | | 0 | 0.0 | | 0 | 0.0 | | 0 | 0.0 | |
| Neoplasia | 0 | 0.0 | | 0 | 0.0 | | 0 | 0.0 | | 0 | 0.0 | |
| Hyperplasia | 0 | 0.0 | | 0 | 0.0 | | 0 | 0.0 | | 0 | 0.0 | |
| Vasculitis | 0 | 0.0 | | 0 | 0.0 | | 0 | 0.0 | | 0 | 0.0 | |
| Atrophy | 0 | 0.0 | | 0 | 0.0 | | 0 | 0.0 | | 0 | 0.0 | |
| Lymphoid Depletion | 0 | 0.0 | | 5 | 2.7 | 3.2 | 0 | 0.0 | | 0 | 0.0 | |
| Artifact | 0 | 0.0 | | 0 | 0.0 | | 0 | 0.0 | | 0 | 0.0 | |
| Sum - Scores: | 4 | 0.8 | 1.5 | 6 | 2.8 | 4.2 | 5 | 0.9 | 2.5 | 0 | 0.0 | 0.0 |
| No Significant Findings: | 4 | | | 1 | | | 4 | | | 8 | | |
| | **n=** | **8** | | **n=** | **6** | | **n=** | **8** | | **n=** | **8** | |
| **4- Lung** | | | | | | | | | | | | |
| Hemorrhage | 1 | 0.1 | 1.0 | 1 | 0.3 | 2.0 | 0 | 0.0 | | 0 | 0.0 | |
| Edema | 0 | 0.0 | | 0 | 0.0 | | 0 | 0.0 | | 0 | 0.0 | |
| Inflammation | 0 | 0.0 | | 0 | 0.0 | | 0 | 0.0 | | 0 | 0.0 | |
| Increased cellularity, alveolar | 5 | 0.6 | 1.0 | 5 | 0.8 | 1.0 | 4 | 0.5 | 1.0 | 5 | 0.6 | 1.0 |
| Necrosis | 0 | 0.0 | | 0 | 0.0 | | 0 | 0.0 | | 0 | 0.0 | |
| Autolysis | 0 | 0.0 | | 0 | 0.0 | | 0 | 0.0 | | 0 | 0.0 | |
| Mineralization | 0 | 0.0 | | 0 | 0.0 | | 0 | 0.0 | | 0 | 0.0 | |
| Neoplasia | 0 | 0.0 | | 0 | 0.0 | | 0 | 0.0 | | 0 | 0.0 | |
| Apoptotic bodies, alveolar | 0 | 0.0 | | 2 | 0.7 | 2.0 | 0 | 0.0 | | 0 | 0.0 | |
| Thrombosis | 0 | 0.0 | | 1 | 0.2 | 1.0 | 0 | 0.0 | | 0 | 0.0 | |
| Atrophy | 0 | 0.0 | | 0 | 0.0 | | 0 | 0.0 | | 0 | 0.0 | |
| Lymphoid Depletion | 0 | 0.0 | | 0 | 0.0 | | 0 | 0.0 | | 0 | 0.0 | |

| | | | | | | | | | | | | |
|---|---|---|---|---|---|---|---|---|---|---|---|---|
| Artifact | 0 | 0.0 | | 0 | 0.0 | | 0 | 0.0 | | 0 | 0.0 | |
| Sum - Scores: | 6 | 0.8 | 2.0 | 9 | 2.0 | 6.0 | 4 | 0.5 | 1.0 | 5 | 0.6 | 1.0 |
| No Significant Findings: | 3 | | | 1 | | | 4 | | | 3 | | |
| | **n=** | **8** | | **n=** | **6** | | **n=** | **8** | | **n=** | **8** | |
| **5- Duodenum** | | | | | | | | | | | | |
| Hemorrhage | 0 | 0.0 | | 0 | 0.0 | | 0 | 0.0 | | 0 | 0.0 | |
| Edema | 0 | 0.0 | | 2 | 0.3 | 1.0 | 1 | 0.1 | 1.0 | 0 | 0.0 | |
| Inflammation, mural | 0 | 0.0 | | 0 | 0.0 | | 0 | 0.0 | | 0 | 0.0 | |
| Degeneration | 0 | 0.0 | | 4 | 1.1 | 1.6 | 0 | 0.0 | | 8 | 1.6 | 1.6 |
| Necrosis | 0 | 0.0 | | 0 | 0.0 | | 0 | 0.0 | | 0 | 0.0 | |
| Inflammation, mesenteric | 0 | 0.0 | | 0 | 0.0 | | 0 | 0.0 | | 2 | 0.4 | 1.5 |
| Mineralization | 0 | 0.0 | | 0 | 0.0 | | 0 | 0.0 | | 0 | 0.0 | |
| Neoplasia | 0 | 0.0 | | 0 | 0.0 | | 0 | 0.0 | | 0 | 0.0 | |
| Hyperplasia | 0 | 0.0 | | 0 | 0.0 | | 0 | 0.0 | | 0 | 0.0 | |
| Vasculitis | 0 | 0.0 | | 0 | 0.0 | | 0 | 0.0 | | 0 | 0.0 | |
| Apoptotic bodies, villi | 0 | 0.0 | | 2 | 0.3 | 1.0 | 0 | 0.0 | | 4 | 0.5 | 1.0 |
| Lymphoid Depletion | 0 | 0.0 | | 1 | 0.7 | 4.0 | 0 | 0.0 | | 0 | 0.0 | |
| Artifact | 2 | 0.3 | 1.0 | 3 | 0.8 | 1.7 | 3 | 0.6 | 1.7 | 4 | 1.0 | 2.0 |
| Sum - Scores: | 0 | 0.0 | 0.0 | 9 | 2.4 | 7.6 | 1 | 0.1 | 1.0 | 14 | 2.5 | 4.1 |
| No Significant Findings: | 8 | | | 1 | | | 7 | | | 0 | | |

| | **saline, no heat, 1 day** | | | **saline, heat, 1 day** | | | **saline, no heat, $T_{c,max}$** | | | **saline, heat, $T_{c,max}$** | | |
|---|---|---|---|---|---|---|---|---|---|---|---|---|
| **Number** | **# Abnormal** | **Mean Score** | **Mean-Affected** | **# Abnormal** | **Mean Score** | **Mean-Affected** | **# Abnormal** | **Mean Score** | **Mean-Affected** | **# Abnormal** | **Mean Score** | **Mean-Affected** |
| | **n=** | **8** | | **n=** | **8** | | **n=** | **8** | | **n=** | **8** | |
| **1- Liver** | | | | | | | | | | | | |
| Extramedullary hematopoiesis | 4 | 0.5 | 1.0 | 0 | 0.0 | | 2 | 0.3 | 1.0 | 0 | 0.0 | |
| Vacuolation, hepatocellular | 8 | 2.6 | 2.6 | 8 | 2.8 | 2.8 | 8 | 1.9 | 1.9 | 1 | 0.1 | 1.0 |
| | 5 | 0.6 | 1.0 | 6 | 0.8 | 1.0 | 3 | 0.4 | 1.0 | 7 | 1.4 | 1.6 |

| | | | | | | | | | | | | |
|---|---|---|---|---|---|---|---|---|---|---|---|---|
| Inflammation, intrahepatic | | | | | | | | | | | | |
| Inflammation, mesentery/ capsule | 1 | 0.3 | 2.0 | 0 | 0.0 | | 1 | 0.1 | 1.0 | 1 | 0.1 | 1.0 |
| Necrosis | 0 | 0.0 | | 0 | 0.0 | | 0 | 0.0 | | 0 | 0.0 | |
| Degeneration, hepatocellular | 0 | 0.0 | | 6 | 1.5 | 2.0 | 1 | 0.1 | 1.0 | 4 | 0.8 | 1.6 |
| Mineralization | 0 | 0.0 | | 0 | 0.0 | | 0 | 0.0 | | 0 | 0.0 | |
| Neoplasia | 0 | 0.0 | | 0 | 0.0 | | 0 | 0.0 | | 0 | 0.0 | |
| Hyperplasia, oval cell | 1 | 0.1 | 1.0 | 0 | 0.0 | | 0 | 0.0 | | 1 | 0.1 | 1.0 |
| Vasculitis | 0 | 0.0 | | 0 | 0.0 | | 0 | 0.0 | | 0 | 0.0 | |
| Atrophy | 0 | 0.0 | | 0 | 0.0 | | 0 | 0.0 | | 0 | 0.0 | |
| Lymphoid Depletion | 0 | 0.0 | | 0 | 0.0 | | 0 | 0.0 | | 0 | 0.0 | |
| Artifact | 1 | 0.3 | 2.0 | 3 | 0.8 | 2.0 | 4 | 0.9 | 1.8 | 3 | 0.6 | 1.7 |
| Sum - Scores: | 19 | 4.1 | 7.6 | 20 | 5.0 | 5.8 | 15 | 2.8 | 5.9 | 14 | 2.6 | 6.3 |
| No Significant Findings: | 0 | | | 0 | | | 0 | | | 0 | | |
| | **n=** | **8** | | **n=** | **8** | | **n=** | **8** | | **n=** | **8** | |
| **2- Kidney** | | | | | | | | | | | | |
| Hemorrhage | 0 | 0.0 | | 0 | 0.0 | | 0 | 0.0 | | 0 | 0.0 | |
| Edema | 0 | 0.0 | | 0 | 0.0 | | 0 | 0.0 | | 0 | 0.0 | |
| Inflammation | 0 | 0.0 | | 0 | 0.0 | | 0 | 0.0 | | 0 | 0.0 | |
| Degeneration | 0 | 0.0 | | 0 | 0.0 | | 0 | 0.0 | | 0 | 0.0 | |
| Necrosis | 0 | 0.0 | | 0 | 0.0 | | 0 | 0.0 | | 0 | 0.0 | |
| Autolysis | 0 | 0.0 | | 0 | 0.0 | | 0 | 0.0 | | 0 | 0.0 | |
| Mineralization | 0 | 0.0 | | 0 | 0.0 | | 0 | 0.0 | | 0 | 0.0 | |
| Neoplasia | 0 | 0.0 | | 0 | 0.0 | | 0 | 0.0 | | 0 | 0.0 | |
| Hyperplasia | 0 | 0.0 | | 0 | 0.0 | | 0 | 0.0 | | 0 | 0.0 | |
| Vasculitis | 0 | 0.0 | | 0 | 0.0 | | 0 | 0.0 | | 0 | 0.0 | |
| Atrophy | 0 | 0.0 | | 0 | 0.0 | | 0 | 0.0 | | 0 | 0.0 | |
| Lymphoid Depletion | 0 | 0.0 | | 0 | 0.0 | | 0 | 0.0 | | 0 | 0.0 | |
| Artifact | 1 | 0.3 | 2.0 | 7 | 2.4 | 2.8 | 3 | 0.8 | 2.0 | 2 | 0.5 | 2.0 |
| Sum - Scores: | 0 | 0.0 | 0.0 | 0 | 0.0 | 0.0 | 0 | 0.0 | 0.0 | 0 | 0.0 | 0.0 |
| No Significant Findings: | 8 | | | 8 | | | 8 | | | 8 | | |
| | **n=** | **8** | | **n=** | **8** | | **n=** | **8** | | **n=** | **8** | |
| **3- Spleen** | | | | | | | | | | | | |
| Hemorrhage | 0 | 0.0 | | 0 | 0.0 | | 0 | 0.0 | | 0 | 0.0 | |
| Increased neutrophils, red pulp | 0 | 0.0 | | 0 | 0.0 | | 0 | 0.0 | | 0 | 0.0 | |
| Inflammation, mesenteric | 3 | 0.5 | 1.3 | 3 | 0.4 | 1.0 | 2 | 0.5 | 2.0 | 3 | 0.6 | 1.7 |
| Degeneration | 0 | 0.0 | | 0 | 0.0 | | 0 | 0.0 | | 0 | 0.0 | |
| Necrosis | 0 | 0.0 | | 0 | 0.0 | | 0 | 0.0 | | 0 | 0.0 | |
| Autolysis | 0 | 0.0 | | 0 | 0.0 | | 0 | 0.0 | | 0 | 0.0 | |

| | | | | | | | | | | | | |
|---|---|---|---|---|---|---|---|---|---|---|---|---|
| Mineralization | 0 | 0.0 | | 0 | 0.0 | | 0 | 0.0 | | 0 | 0.0 | |
| Neoplasia | 0 | 0.0 | | 0 | 0.0 | | 0 | 0.0 | | 0 | 0.0 | |
| Hyperplasia | 0 | 0.0 | | 0 | 0.0 | | 0 | 0.0 | | 0 | 0.0 | |
| Vasculitis | 0 | 0.0 | | 0 | 0.0 | | 0 | 0.0 | | 0 | 0.0 | |
| Atrophy | 0 | 0.0 | | 0 | 0.0 | | 0 | 0.0 | | 0 | 0.0 | |
| Lymphoid Depletion | 0 | 0.0 | | 1 | 0.1 | 1.0 | 0 | 0.0 | | 0 | 0.0 | |
| Artifact | 0 | 0.0 | | 0 | 0.0 | | 0 | 0.0 | | 0 | 0.0 | |
| Sum - Scores: | 3 | 0.5 | 1.3 | 4 | 0.5 | 2.0 | 2 | 0.5 | 2.0 | 3 | 0.6 | 1.7 |
| No Significant Findings: | 5 | | | 5 | | | 6 | | | 5 | | |
| | **n=** | **8** | | **n=** | **8** | | **n=** | **8** | | **n=** | **8** | |
| **4- Lung** | | | | | | | | | | | | |
| Hemorrhage | 0 | 0.0 | | 0 | 0.0 | | 0 | 0.0 | | 1 | 0.1 | 1.0 |
| Edema | 0 | 0.0 | | 0 | 0.0 | | 0 | 0.0 | | 0 | 0.0 | |
| Inflammation | 0 | 0.0 | | 0 | 0.0 | | 0 | 0.0 | | 0 | 0.0 | |
| Increased cellularity, alveolar | 8 | 1.3 | 1.3 | 6 | 0.8 | 1.0 | 4 | 0.6 | 1.3 | 4 | 0.6 | 1.3 |
| Necrosis | 0 | 0.0 | | 0 | 0.0 | | 0 | 0.0 | | 0 | 0.0 | |
| Autolysis | 0 | 0.0 | | 0 | 0.0 | | 0 | 0.0 | | 0 | 0.0 | |
| Mineralization | 0 | 0.0 | | 0 | 0.0 | | 0 | 0.0 | | 0 | 0.0 | |
| Neoplasia | 0 | 0.0 | | 0 | 0.0 | | 0 | 0.0 | | 0 | 0.0 | |
| Apoptotic bodies, alveolar | 0 | 0.0 | | 0 | 0.0 | | 0 | 0.0 | | 0 | 0.0 | |
| Thrombosis | 0 | 0.0 | | 0 | 0.0 | | 0 | 0.0 | | 1 | 0.1 | 1.0 |
| Atrophy | 0 | 0.0 | | 0 | 0.0 | | 0 | 0.0 | | 0 | 0.0 | |
| Lymphoid Depletion | 0 | 0.0 | | 0 | 0.0 | | 0 | 0.0 | | 0 | 0.0 | |
| Artifact | 0 | 0.0 | | 1 | 0.4 | 3.0 | 0 | 0.0 | | 1 | 0.1 | 1.0 |
| Sum - Scores: | 8 | 1.3 | 1.3 | 6 | 0.8 | 1.0 | 4 | 0.6 | 1.3 | 6 | 0.9 | 3.3 |
| No Significant Findings: | 0 | | | 2 | | | 4 | | | 4 | | |
| | **n=** | **8** | | **n=** | **8** | | **n=** | **8** | | **n=** | **8** | |
| **5- Duodenum** | | | | | | | | | | | | |
| Hemorrhage | 0 | 0.0 | | 0 | 0.0 | | 0 | 0.0 | | 0 | 0.0 | |
| Edema | 0 | 0.0 | | 0 | 0.0 | | 0 | 0.0 | | 0 | 0.0 | |
| Inflammation, mural | 0 | 0.0 | | 0 | 0.0 | | 0 | 0.0 | | 0 | 0.0 | |
| Degeneration | 0 | 0.0 | | 0 | 0.0 | | 0 | 0.0 | | 0 | 0.0 | |
| Necrosis | 0 | 0.0 | | 0 | 0.0 | | 0 | 0.0 | | 0 | 0.0 | |
| Inflammation, mesenteric | 1 | 0.1 | 1.0 | 0 | 0.0 | | 0 | 0.0 | | 0 | 0.0 | |
| Mineralization | 0 | 0.0 | | 0 | 0.0 | | 0 | 0.0 | | 0 | 0.0 | |
| Neoplasia | 0 | 0.0 | | 0 | 0.0 | | 0 | 0.0 | | 0 | 0.0 | |
| Hyperplasia | 0 | 0.0 | | 0 | 0.0 | | 0 | 0.0 | | 0 | 0.0 | |
| Vasculitis | 0 | 0.0 | | 0 | 0.0 | | 0 | 0.0 | | 0 | 0.0 | |

| | | | | | | | | | | | | |
|---|---|---|---|---|---|---|---|---|---|---|---|---|
| Apoptotic bodies, villi | 0 | 0.0 | | 0 | 0.0 | | 0 | 0.0 | | 5 | 0.6 | 1.0 |
| Lymphoid Depletion | 0 | 0.0 | | 0 | 0.0 | | 0 | 0.0 | | 0 | 0.0 | |
| Artifact | 0 | 0.0 | | 2 | 0.3 | 1.0 | 5 | 1.1 | 1.8 | 8 | 2.0 | 2.0 |
| Sum - Scores: | 1 | 0.1 | 1.0 | 0 | 0.0 | 0.0 | 0 | 0.0 | 0.0 | 5 | 0.6 | 1.0 |
| No Significant Findings: | 7 | | | 8 | | | 8 | | | 3 | | |
